# Distinct modes of folding assistance in cells by protein- and RNA-based chaperones

**DOI:** 10.64898/2026.04.15.718697

**Authors:** Ahhyun Son, Catherine Durso, Timothy A. Whitehead, Scott Horowitz

**Affiliations:** Department of Chemistry & Biochemistry and the Knoebel Institute for Healthy Aging, University of Denver; Denver, CO 80208, USA; Daniel Felix Ritchie School of Engineering and Computer Science, University of Denver; Denver, CO 80208, USA; Department of Chemical and Biological Engineering, University of Colorado; Boulder, CO 80305, USA

**Author notes:** Corresponding authors: Scott Horowitz and Ahhyun Son, **Email:** and.

**Keywords:** protein folding, molecular chaperones, proteostasis, G-quadruplex, CHAP-SEQ

## Abstract

How chaperones mediate protein folding in the crowded cell environment remains poorly understood. To gain insight, we developed CHAP-SEQ, a high-throughput approach for examining how chaperones affect protein folding in cells at amino acid resolution. Performing CHAP-SEQ using three protein-based chaperones and one RNA-based chaperone revealed distinct modes of folding assistance. The protein-based chaperones preferentially act in a partially redundant manner on hydrophobic core residues, whereas the RNA-based chaperone primarily targets structural or dynamic features of the client protein. Furthermore, while the chaperone RNA has little preference for client variants’ baseline stability, the protein-based chaperones preferentially assisted clients with greater intrinsic foldability. Together, these findings demonstrated that protein– and RNA-based chaperones promote protein folding through mechanistically distinct modes of action.

**Significance Statement:** Determining the structural basis of how individual chaperones act on client proteins remains challenging, particularly within living cells. This knowledge gap is especially pronounced for RNA-based chaperones, whose mechanisms of action remain largely unknown. Here we introduce CHAP-SEQ, a technique that maps chaperone effects across a client protein at the level of individual amino acid residues. CHAP-SEQ revealed distinct residue-level patterns of folding assistance by protein– and RNA-based chaperones, and provided insight into both how different chaperones act in a partially redundant, partially complementary manner to aid folding in cells.

## Introduction

Chaperones are central regulators of protein folding and protein homeostasis in cells (1–5). Because nearly all cellular processes depend on proteins adopting and maintaining functional three-dimensional structures, understanding how chaperones influence protein folding is essential for explaining both normal cellular function and the loss of protein homeostasis associated with stress, aging, and disease. Protein-based chaperones have therefore been studied for decades. However, much of our mechanistic understanding has been derived from biochemical and structural studies in vitro (6–9). Although these studies have established important models of chaperone action, it remains difficult to assess how these mechanisms operate within the crowded and heterogeneous intracellular environment.

A major limitation has been the lack of approaches that directly measure chaperone effects on protein folding in cells at high resolution, making it difficult to compare chaperones with fundamentally different modes of action under a common experimental framework. As an example, although accumulating evidence indicates that RNA molecules can modulate protein folding and aggregation (10–12), their residue-level mechanisms and their similarities to or differences from protein-based chaperones remain largely unknown.

To address these challenges, we introduce CHAP-SEQ, a high-throughput approach adapted from fluorescence-based sort-and-sequence technologies (13–16) to determine how different chaperones affect protein folding in cells at residue-level resolution. CHAP-SEQ uses the folding biosensor protein TagRFP675, which requires overexpressed chaperones to fold and fluoresce appreciably in *E. coli*; its fluorescence in *E. coli* is strongly dependent on its folding quality (10, 12). In addition, we previously showed that under conditions of insufficient chaperoning capacity, poorly folded TagRFP675 is predominantly degraded rather than accumulating as stable intermediates or insoluble aggregates (12). TagRFP675 therefore provides a robust folding-dependent cellular readout and its responses to multiple protein– and RNA-based chaperones have been established in both cells and purified-protein assays.

Here, we generated an alanine-scanning mutational library spanning the TagRFP675 sequence and expressed the library in *E. coli* MC4100(DE3) together with either an empty vector or one of four chaperones. An alanine-scanning design was selected to provide a readily interpretable perturbation at each position, thereby facilitating direct residue-level comparisons across chaperone conditions and structural interpretation of native side-chain contributions. Fluorescence-based sorting followed by deep sequencing (17) then allowed us to quantify how each alanine substitution altered folding under each chaperone condition.

The four chaperones were deliberately selected to span distinct mechanistic classes of folding assistance. GroEL, the bacterial Hsp60 chaperonin, uses ATP cycling to encapsulate client proteins (18, 19). DnaK, the bacterial Hsp70, recognizes exposed peptide segments and uses ATP-dependent cycles of binding and release to limit nonproductive interactions and facilitate folding (20, 21); Spy is an ATP-independent chaperone normally localized to the periplasm that provides a dynamic protective environment in which client proteins can remain partially folded or undergo refolding from stresses (22, 23); here, Spy was expressed in the cytoplasm to enable a direct comparison under the same cellular conditions. The RNA G-quadruplex (G4) Seq576 is an RNA-based chaperone that promotes protein folding in vitro and in cells while remaining associated with its client (10, 12, 24). All four chaperones were previously shown to enhance TagRFP675 folding in *E. coli* (10, 12), making them suitable for a controlled comparison of ATP-dependent, ATP-independent, and RNA-mediated folding assistance.

This experimental design enabled us to ask whether chaperones with distinct chemical mechanisms act on shared or different features of the same client protein and whether protein– and RNA-based chaperones act differently upon clients in cells. The resulting maps uncover previously unrecognized distinctions in how diverse chaperone systems navigate the same folding landscape in cells.

## Results

### CHAP-SEQ Enables Quantitative, Residue-Level Mapping of Chaperone-Dependent Folding in Cells

CHAP-SEQ uses fluorescence-activated cell sorting (FACS) to separate an alanine-scanning library according to TagRFP675 fluorescence, which serves as a readout of folding quality (10, 12). DAPI was added to exclude dead or dying cells, and the remaining population was divided into low– and high-fluorescence gates separated by a common fluorescence threshold, with approximately half of the parent population assigned to each gate. The sorted libraries were then deep sequenced to determine relative abundance of each single-mutant variant in the low– or high-fluorescence populations (Fig. 1A and B).

**Figure 1.**
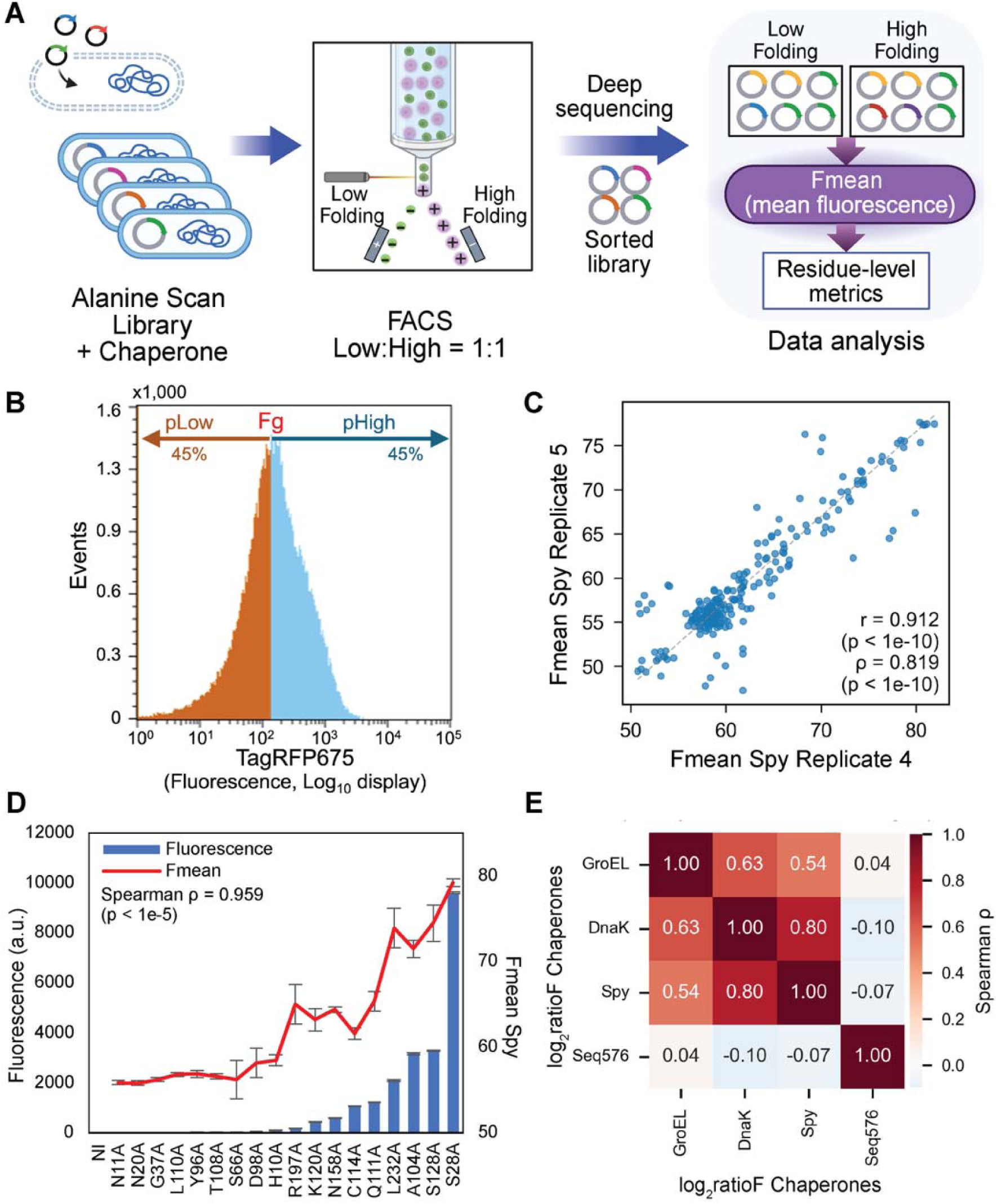
CHAP-SEQ workflow and validation. (A) Schematic overview of the CHAP-SEQ workflow. An alanine-scanning library of the folding biosensor TagRFP675 was expressed in *E. coli* together with individual chaperones. Cells were separated by fluorescence using FACS into low– and high-folding populations. Deep sequencing of sorted libraries was used to estimate variant frequencies and compute mean fluorescence (Fmean) values for each variant. This approach enables residue-level comparison of how different chaperones act on protein folding, providing insight into their underlying mechanisms. (B) Representative fluorescence-activated cell sorting (FACS) histogram from an experimental dataset, showing gating used to separate low– and high-folding variants at a 1:1 gate boundary. Approximately 10-20% of high DAPI (cell death) events were excluded, resulting in 40-45% of cells in each gate. Full FACS datasets and sorting images are shown in Figs. S2 and S3. (**C**) Reproducibility across biological replicates. Scatter plot of residue-level Fmean values from two representative Spy replicates (Spy Rep 4 and 5) shows strong agreement. Each point represents a single-residue substitution. Full plots are shown in Fig. S7. (**D**) Orthogonal validation of CHAP-SEQ measurements. Plate reader measurements of fluorescence for individual variants (at OD_600_ = 1.0) correlate with CHAP-SEQ-derived Fmean values (Spy condition, right y-axis). Background fluorescence was minimal based on non-induced (NI) controls. Full plots are shown in Fig. S8. (**E**) Condition-level correlation of residue-specific folding effects. Spearman correlation matrix of residue-level log_2_ratioF values (log_2_(Fmean_Chaperone_/Fmean_Empty_)) across chaperone conditions. Protein chaperones (GroEL, DnaK, Spy) show strong inter-correlation, whereas Seq576 displays minimal correlation with the protein chaperones, indicating a distinct mode of folding modulation. All pairwise scatter plots and Pearson correlation heatmap are shown in Fig. S10.

We implemented a multistep quality-control workflow to verify that FACS reproducibly separated libraries with distinct folding phenotypes. The initially sorted low– and high-fluorescence populations were then reanalyzed by FACS to determine whether they retained their expected fluorescence distributions. DNA libraries were subsequently purified from the sorted populations, re-transformed into fresh cells, and evaluated again by FACS and bulk fluorescence measurements. Libraries that reproducibly retained their respective low– or high-fluorescence phenotypes were advanced for deep sequencing (Figs. S1-S3; Table S2). Raw sequencing reads were then processed using a strict codon-level variant-calling pipeline designed specifically for the GCG alanine-scanning library (Fig. S4). At each targeted position, the pipeline evaluated positional coverage and complete GCG substitution evidence; incomplete or unsupported variant calls were explicitly flagged and prevented from contributing valid variant counts. Subsequent deep sequencing quality-control analyses further confirmed sufficient read recovery and consistent variant representation across the retained biological replicates and experimental conditions (Table S3-S4). Together, these analyses demonstrate that the CHAP-SEQ workflow reliably isolated and recovered mutant libraries with reproducibly distinct folding outcomes, and generated high-quality variant-frequency data for quantitative analysis.

For each variant, deep sequencing read counts from the low– and high-fluorescence populations were converted to variant frequencies and used to calculate pHigh, the probability of the variant falling above the FACS gate threshold. Using the previously established log-normal fluorescence distribution model (15), we combined pHigh with the gate threshold and the empirically determined distribution-width parameter σ derived from the FACS data. Numerical inversion of this relationship yielded an estimated mean fluorescence value (Fmean) for each variant (Fig. S5 and Table S5; Supplemental Materials and Methods). Thus, Fmean is a sequencing-based estimate of cellular fluorescence rather than a direct fluorescence measurement. This conversion enabled quantitative comparisons of folding outcomes among variants and chaperone conditions.

We next evaluated the reproducibility and accuracy of the calculated Fmean values using three complementary approaches. First, we compared variant-level Fmean values across three biological replicates for each chaperone and Empty condition. The replicates showed highly significant pairwise correlations and low dispersion, with Pearson r = 0.644*–*0.940 (all p < 1×10^-10^) (Figs. 1C and S6-S7; Table S6). Second, we determined whether the sequencing-derived Fmean values agreed with fluorescence measured directly in cells. Eighteen individual TagRFP675 variants were expressed separately across each chaperone condition, and their fluorescence was measured using a plate reader. The directly measured fluorescence values were strongly correlated with the corresponding CHAP-SEQ-derived Fmean estimates, with Spearman correlation coefficients (ρ) ranging from 0.845 to 0.959 (all p < 1×10^-4^) (Figs. 1D and S8). Finally, we examined whether the chaperone-dependent fluorescence effects observed in cells were consistent with direct effects on protein folding. Two individual variants, TagRFP675(Q111A) and TagRFP675(L232A), were purified and subjected to *in vitro* refolding assays in the presence or absence of G4 Seq576. Seq576 enhanced the refolding of both variants, and the direction of these effects was consistent with the corresponding changes observed in *E. coli* by CHAP-SEQ (Fig. S9). Together, the reproducibility across biological replicates, agreement with orthogonal cellular fluorescence measurements, and consistency with *in vitro* refolding establish CHAP-SEQ as a robust approach for quantifying chaperone-dependent folding effects in cells at residue-level resolution.

### Protein– and RNA-Based Chaperones Exhibit Distinct Residue-Level Folding Signatures

With CHAP-SEQ validated, we next analyzed chaperone-assisted folding of TagRFP675 in *E. coli* at residue-level resolution by comparing each alanine variant under a chaperone condition with the same variant in the matched Empty condition. This within-variant normalization was designed to isolate the contribution of chaperone expression while controlling for the baseline stability and fluorescence of each mutant. We therefore defined a folding-effect score, calculated as log_2_(Fmean_Chaperone_ / Fmean_Empty_), hereafter referred to as log_2_ratioF. Because the same alanine variant was compared between matched conditions, log_2_ratioF reflects the magnitude and direction of chaperone-dependent folding assistance at each amino acid residue rather than absolute fluorescence. To interpret these variant-level effects in terms of the WT sequence, we took advantage of using alanine substitutions that remove most native side-chain functionality while retaining the peptide backbone. Therefore, a relatively lower log_2_ratioF indicates that the alanine-substituted variant at that position showed a weaker chaperone-dependent folding response than variants with higher scores. When mapped back to the WT sequence, this reduced response is interpreted as evidence that the corresponding native side chain in TagRFP675 makes a greater contribution to chaperone-assisted folding of TagRFP675. Conversely, a relatively higher log_2_ratioF indicates that the alanine-substituted variant exhibited a stronger chaperone-dependent folding response, suggesting that the corresponding native side is less important for chaperone-assisted folding. This WT-oriented interpretation therefore enables direct comparison of relative chaperone dependence across conditions at residue-level resolution.

We first examined whether residue-level folding-effect scores were correlated across chaperone conditions. Pairwise comparisons of residue-level effects revealed moderate correlations (Spearman ρ = 0.537*–*0.797, Pearson r = 0.453*–*0.770) among protein-based chaperones, indicating partially shared patterns of folding assistance in cells. Interestingly, there was little to no correlation between the protein-based chaperones and Seq576 (Spearman ρ = – 0.097*–*0.043, Pearson r = –0.058*–*0.097), revealing a distinct residue-level folding signature for the RNA-based chaperone Seq576 as compared to the protein-based chaperones (Figs. 1E and S10).

Given the distinct residue-level folding signatures observed across chaperone conditions, we next examined which classes of client amino acids contributed most strongly to chaperone-assisted folding (Figs. 2 and S11). All three protein-based chaperones showed substantial selectivity based on amino acid properties, with hydrophobic residues consistently enriched among residues important for chaperone-dependent folding (Figs. 2A*–*C and S11A*–*C). For DnaK and GroEL especially, these observations are consistent with previous studies showing that protein-based chaperones preferentially recognize or act on exposed hydrophobic segments of client proteins during folding (25–27). On the other hand, Seq576 showed no significant overall preference among amino acid classes (Figs. 2D and S11D), indicating that amino acid identity alone does not explain the G4’s folding effects.

**Figure 2.**
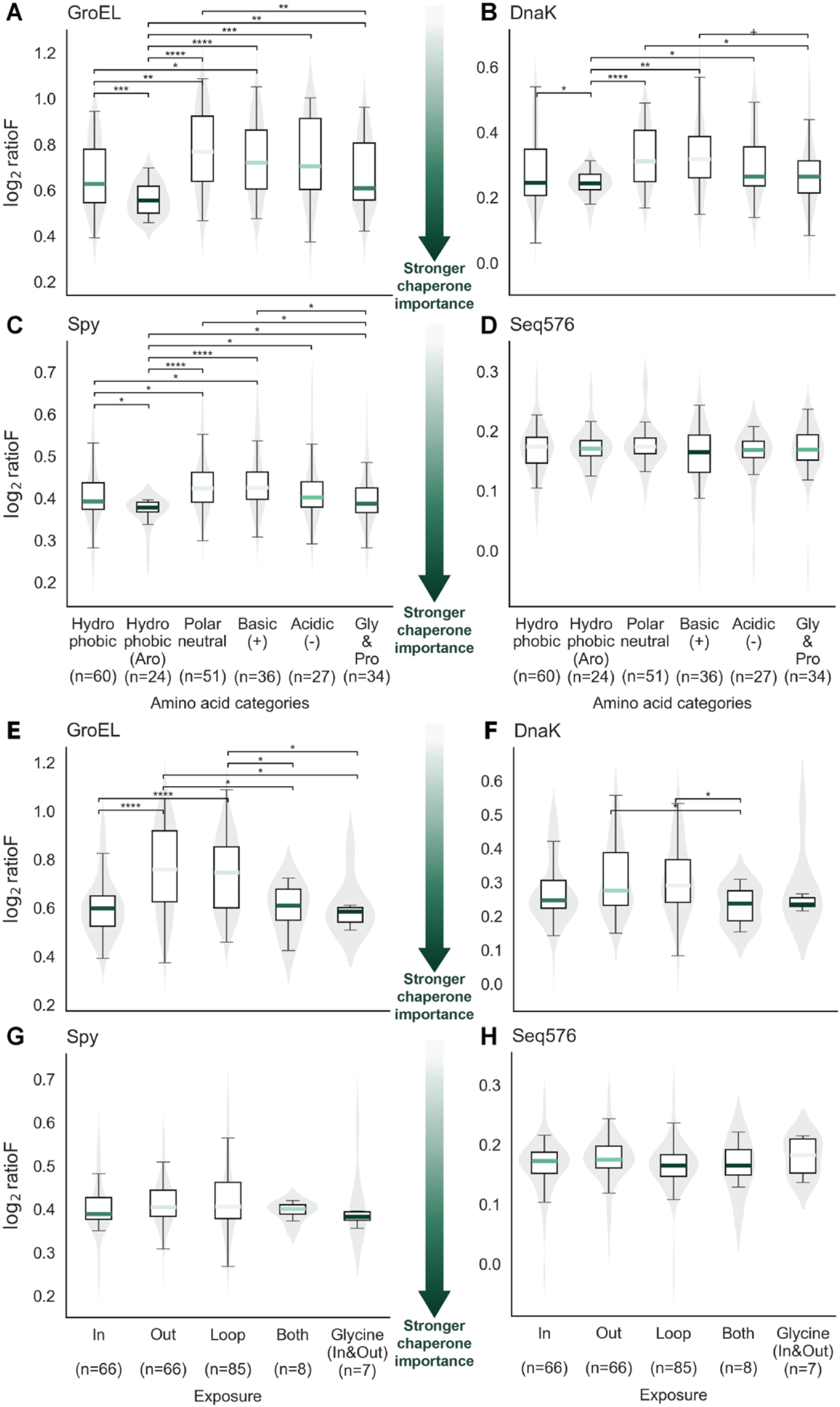
Amino acid and structural preferences of chaperone-dependent folding effects. WT-oriented residue-level log_2_ratioF distributions were grouped by amino acid class and structural exposure. Lower values indicate residues more important for chaperone function under the WT-oriented interpretation, as mutations at these positions more strongly impair chaperone-assisted folding. (**A–D**) Distribution of log_2_ratioF values across amino acid categories for GroEL (**A**), DnaK (**B**), Spy (**C**), and Seq576 (**D**). Lower values indicate residues more important for chaperone-dependent folding under the WT-oriented interpretation. Violin plots show distribution density, with boxplots indicating medians and interquartile ranges. Pairwise comparisons were performed using Welch’s t-test with Benjamini-Hochberg FDR correction. (**E–H**) Distribution of log_2_ratioF values grouped by structural exposure class (In, Out, Loop, Both, Glycine) for the same conditions in the same order as in (**A–D**). Combined heatmaps summarizing both amino acid categories and structural exposure class are shown in Fig. S11.

We next examined whether chaperone dependence differed between buried and surface-exposed residues (Figs. 2E*–*H and S11). The three protein-based chaperones showed clear residue– and structure-dependent preferences, with stronger effects generally associated with hydrophobic and/or buried residues, whereas Seq576 showed comparatively little overall preference across these categories. For DnaK and GroEL this was expected based on previous analyses, but more surprising for Spy as an ATP-independent chaperone that cannot cycle ATP to work at the interior of the protein. Despite Seq576’s lack of overall preference (Fig 2D), stratifying residues by both amino acid class and structural context revealed some context-dependent preferences for Seq576 (Fig. S11D). Within the β-barrel interior, basic residues showed the strongest dependence on Seq576. Within loop regions, an unbiased comparison across all six amino acid classes showed a loop-specific class-dependent pattern in the relative Seq576 versus protein-based chaperone response (Kruskal–Wallis H = 11.43, *p* = 0.0435; permutation *p* = 0.0502). Based on this pattern, a follow-up comparison showed that charged loop residues exhibited greater relative Seq576 dependence than non-charged loop residues (Mann–Whitney *p* = 0.0021; permutation *p* = 0.0019). Together, these results suggest that, unlike protein-based chaperones that preferentially depend on buried hydrophobic residues to aid protein folding, Seq576 exhibits more context-dependent residue preferences that are shaped by the local structural environment rather than by amino acid class or surface exposure alone.

### Chaperones Exhibit Distinct Structural Patterns of Client Residue Dependence

We next mapped condition-specific log_2_ratioF values onto the three-dimensional structure of TagRFP675 to determine whether each chaperone depended on distinct structural features of the client (Figs. 3 and S12; Movies S1*–*S4). As expected from the amino acid class analysis, GroEL, DnaK, and Spy primarily use residues distributed throughout the interior of the β-barrel to promote folding.

**Figure 3.**
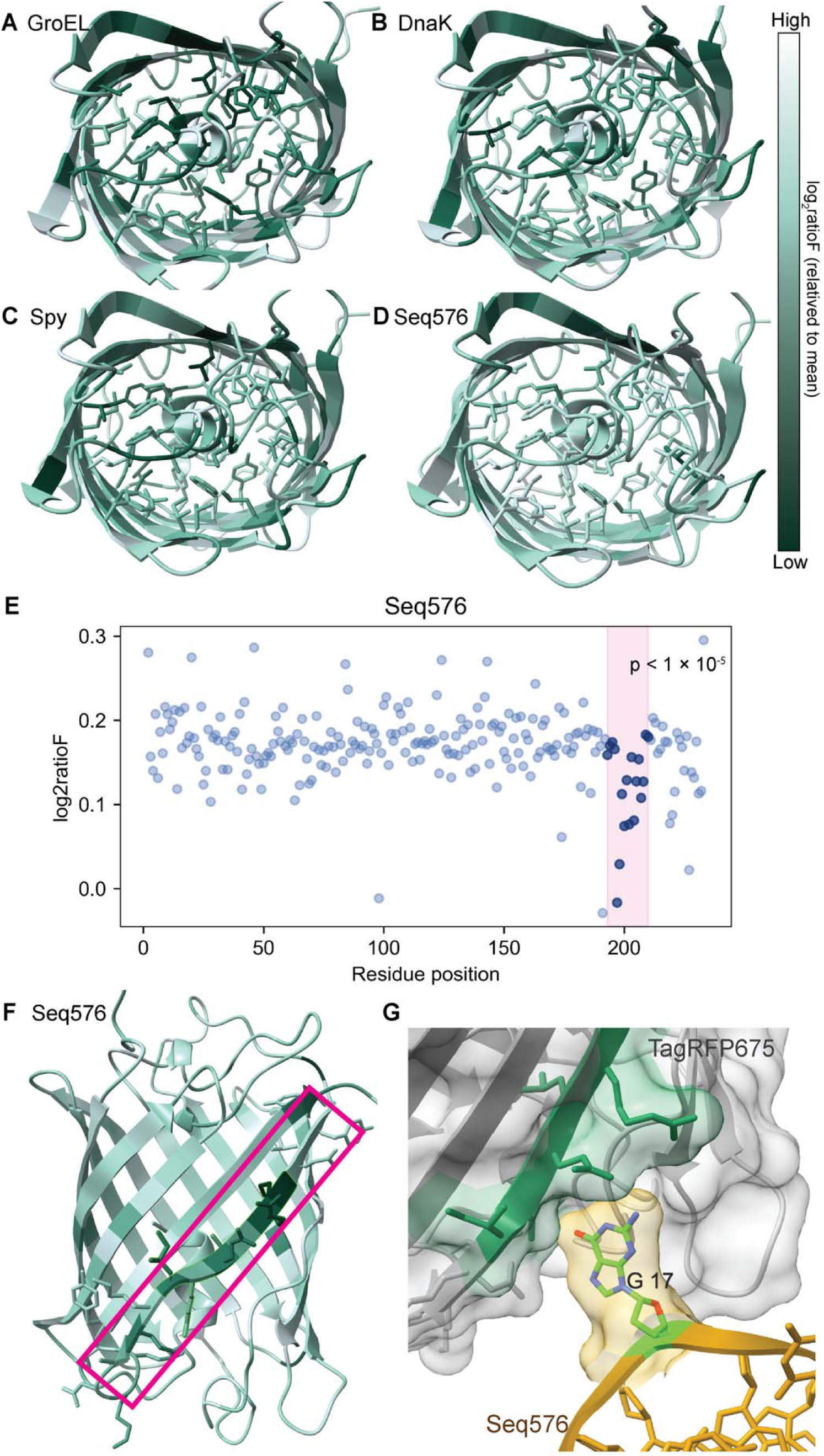
Structural mapping reveals a Seq576-specific β-strand hotspot. (A-D) Projection of residue-level log_2_ratioF values onto the TagRFP675 structure under GroEL (**A**), DnaK (**B**), Spy (**C**), and Seq576 (**D**) conditions. Protein chaperones show broadly distributed effects across the β-barrel interior, whereas Seq576 exhibits a localized signal concentrated on a single β-strand region. log_2_ratioF values are interpreted in a WT-oriented framework: because these values are derived from single-point substitutions, lower values indicate positions where mutation more strongly impairs chaperone-assisted folding, and thus identify WT residues that are more important for chaperone function. Values are scaled independently within each condition relative to the mean (see Methods), and color intensity reflects relative deviation within each condition. (**E, F**) Sliding-window and permutation analysis (window size = 10 residues; lower 15% log_2_ratioF threshold) identifies a statistically significant contiguous cluster uniquely under the Seq576 condition (permutation: residues 197-202, p = 0.0021; significant windows spanning residues 193-210, minimum window p < 1 × 10^-^). The identified region (residues 193-210) is highlighted in magenta, shown as a 1D log_2_ratioF plot (**E**) and mapped onto the 3D structure (**F**). No significant clustering was detected for GroEL, DnaK, or Spy. Full scatter plots are shown in Fig. S12, and full structures and movies are shown in Fig. S13 and Movies S1-S4, respectively. (**G**) Alphafold3-predicted model of TagRFP675 in complex with Seq576 G4 DNA. The Seq576 hotspot β-strand (green) spatially overlaps with the predicted G4 interaction interface (gold). The predicted complex shows high confidence (ipTM = 0.91, pTM = 0.93), supporting the relative positioning of the G4 near the hotspot region, although local interactions are less well resolved due to lower pLDDT of the nucleic acid component, consistent with known challenges in modeling G-quadruplex structures. Nevertheless, the nucleic acid adopts a G4-like stacked conformation.

Seq576 showed a markedly different structural pattern. Important residues were distributed both inside and outside the β-barrel, with a prominent concentration within residues 193*–*210 encompassing the C-terminal β12 region (Figs. 3E and 3F, and S13). Two independent analyses designed to detect contiguous sequence enrichment identified this region as a statistically significant Seq576-specific hotspot, whereas protein-based chaperones showed no significantly enriched contiguous regions (Table S7).

We next examined whether this hotspot for Seq576 was associated with local structural dynamics by ensemble refinement of the TagRFP675 crystal structure (28, 29). This analysis revealed elevated structural variability within the same β12 region (Fig. S14; Movies S5 and S6), and the dynamically variable residues significantly overlapped with residues strongly affected by Seq576 (Table S8). Independently, AlphaFold3 modeling (30) predicted that Seq576 binds to TagRFP675 near the β12 region, with G17 of Seq576 contributing prominently to the predicted interface (Fig. 3G). Notably, G17 was previously identified as the nucleotide most important for the protein folding activity of Seq576 *in vitro* (31). These findings support a model in which G4 Seq576 preferentially assists a dynamic C-terminal β-strand through base-mediated contacts, in contrast to the more broadly distributed interior residue dependencies observed for the protein-based chaperones. The predicted position of G17 further raises the possibility of a base-insertion mechanism.

We next asked whether the broad similarities among the protein-based chaperones reflected shared residue-level dependencies. Because the distributions of log_2_ratioF values differed across conditions, values were converted to z-scores separately within each condition, and negative-z outliers were compared across chaperones. No outlier was shared across Seq576 and all three protein-based chaperones, further supporting the distinct residue-level dependence of the RNA-based chaperone (Fig. 4A and Movie S7). Interestingly, despite the broad similarities and correlations observed among the protein-based chaperones, only one residue, R157, was identified as a concordant outlier across GroEL, DnaK and Spy (Fig. 4B and Movie S8). Additional characterization of R157 is provided in Fig. S15. Chaperone-specific outliers showed distinct structural distributions: GroEL-specific residues were concentrated toward the upper β-barrel and adjacent loops, DnaK-specific residues toward the lower barrel, and Spy-specific residues primarily within loop regions (Fig. 4B and Movie S8). Thus, although the protein-based chaperones share broad preferences for client features, they depend on largely distinct residue sets in structurally defined regions, indicating partially redundant but also complementary modes of folding assistance.

**Figure 4.**
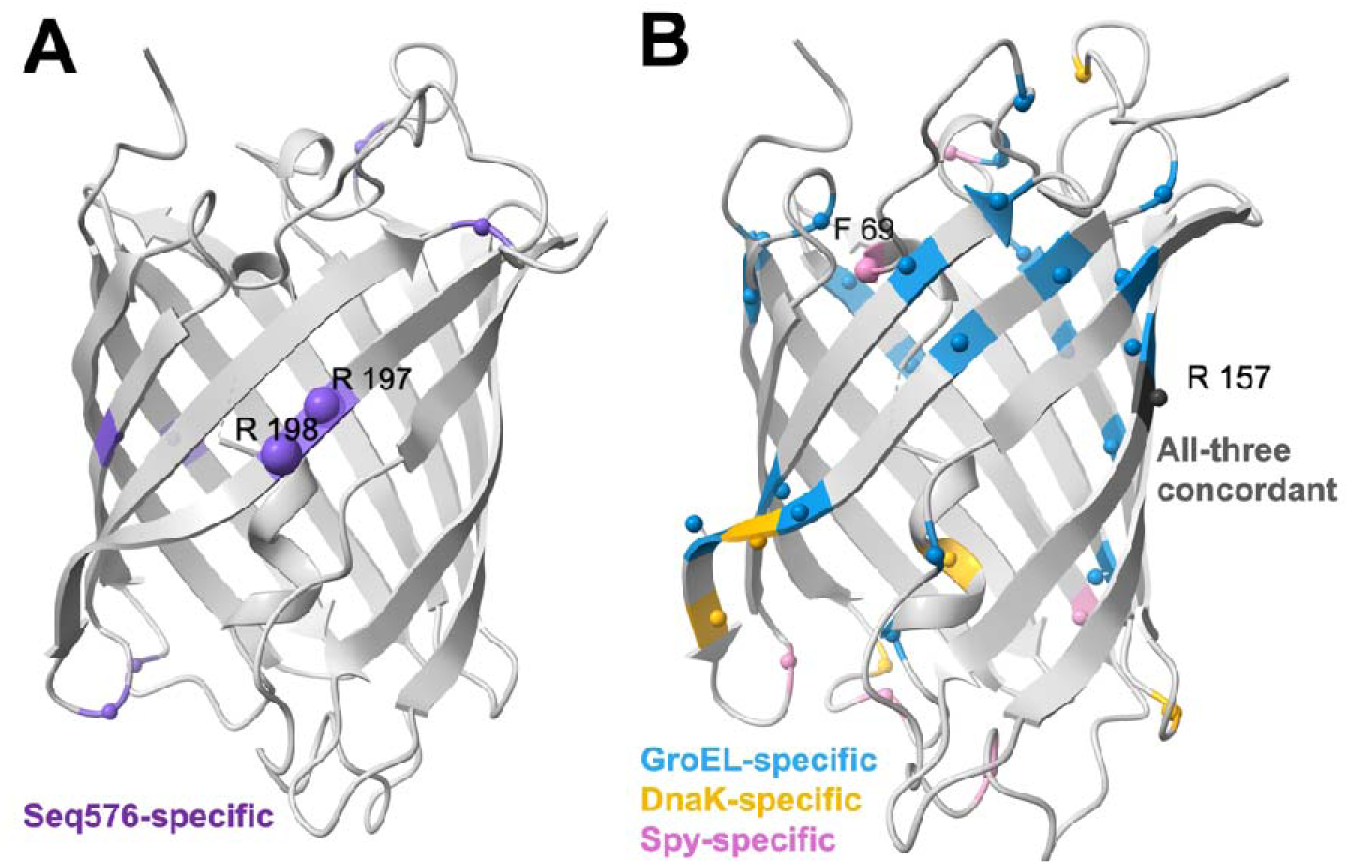
Comparison of log_2_ratioF outliers across chaperone conditions. To compare relative residue-level effectiveness patterns across conditions while accounting for differences in the distributions of log_2_ratioF values, log_2_ratioF values were converted to z scores separately within each chaperone condition. Only negative-z outliers, considered candidate residues important for chaperone function because of their relatively low log2ratioF values, are shown. **(A)** Seq576 (G4)-specific versus protein-based-chaperone-specific outliers. Purple spheres indicate residues classified as Seq576-specific log_2_ratioF outliers, whereas gray spheres indicate log_2_ratioF outliers specific to the protein-based chaperones. **(B)** Protein-chaperone-specific and concordant outliers. Blue, yellow, and pink spheres indicate residues classified as GroEL-, DnaK-, and Spy-specific log_2_ratioF outliers, respectively. The dark gray sphere indicates R157, the only log_2_ratioF outlier concordantly identified across all three protein chaperones. Sphere size reflects the effectiveness-score (ES) category, with the largest spheres representing ES5 residues (z ≤ –3), intermediate-sized spheres representing ES4 residues (–3 < z ≤ –2) and the smallest spheres representing ES3 residues (–2 < z ≤ –1). For residues identified in multiple chaperone conditions, the highest ES category observed in any contributing condition was used. Rotating structural visualizations of the outlier mappings are provided in Movies S7–S8, respectively.

We also examined propagated SD across biological replicates to identify positions with greater variability in chaperone-dependent folding responses (Figs. S16 and S17; Movies S9– S16). Twelve high-SD outliers were shared among the three protein-based chaperones, with the strongest shared outliers located primarily in loops or near β-strand termini rather than in the buried protein interior. GroEL-specific variability extended most prominently into internally packed regions, whereas DnaK– and Spy-specific high-SD outliers were more frequently associated with β-strand termini or loops. These patterns indicate greater variability in chaperone-dependency at flexible or edge-proximal regions than in regularly structured regions. Notably, there were no significant differences in sequencing quality and depth (Tables S3 and S4), indicating that these patterns did not result from uneven sequencing coverage or sample quality.

Taken together, these results reveal broad class-level similarities but distinct residue-level dependencies among the protein-based chaperones, together with a unique β12-centered folding signature for Seq576.

### Protein-Based Chaperones Preferentially Amplify High-Foldability Variants, Unlike Seq576

This dataset afforded the opportunity to investigate whether the chaperones displayed any preference in client foldability. In the absence of additional chaperones, Fmean (Fmean_Empty_) reflects the steady-state folded population of each variant in the cellular environment, integrating intrinsic folding stability, folding kinetics, and degradation (32). Thus, Fmean_Empty_ serves as a proxy for baseline foldability across variants.

Plotting Fmean_Empty_ against Fmean under each chaperone condition provides a framework to assess selectivity for baseline foldability. If the chaperone acts uniformly across variants (*i.e.*, independent of baseline foldability), a slope near 1 is expected. In contrast, slopes greater than 1 indicate that the chaperone preferentially enhances the folding of variants with higher baseline foldability.

Using this framework, the G4 displayed a slope of ∼1.0, with relatively small dispersion, while the protein-based chaperones all displayed slopes greater than 2.0, with considerably greater dispersion (Fig. 5A*–*5D). The relationship between residue-level log_2_ratioF and baseline folding levels (Fmean_Empty_) further confirmed a clear baseline-dependent trend for protein-based chaperones (Fig. S18). The slope near 1.0 for the G4 indicates that it does not show a preference for client foldability, acting similarly across variants with both high and low baseline foldability. In contrast, the protein-based chaperones, which all had higher slopes, showed a preference for clients with higher foldability. Additionally, the protein-based chaperones displayed considerably greater dispersion away from their trendlines than the G4. This broader dispersion indicates that additional variant-specific factors contribute to protein-chaperone-dependent responses beyond baseline foldability alone (Figs. S19–S21). These results indicate that the protein-based chaperones exhibited preferential amplification of folding for high-folding variants.

**Figure 5.**
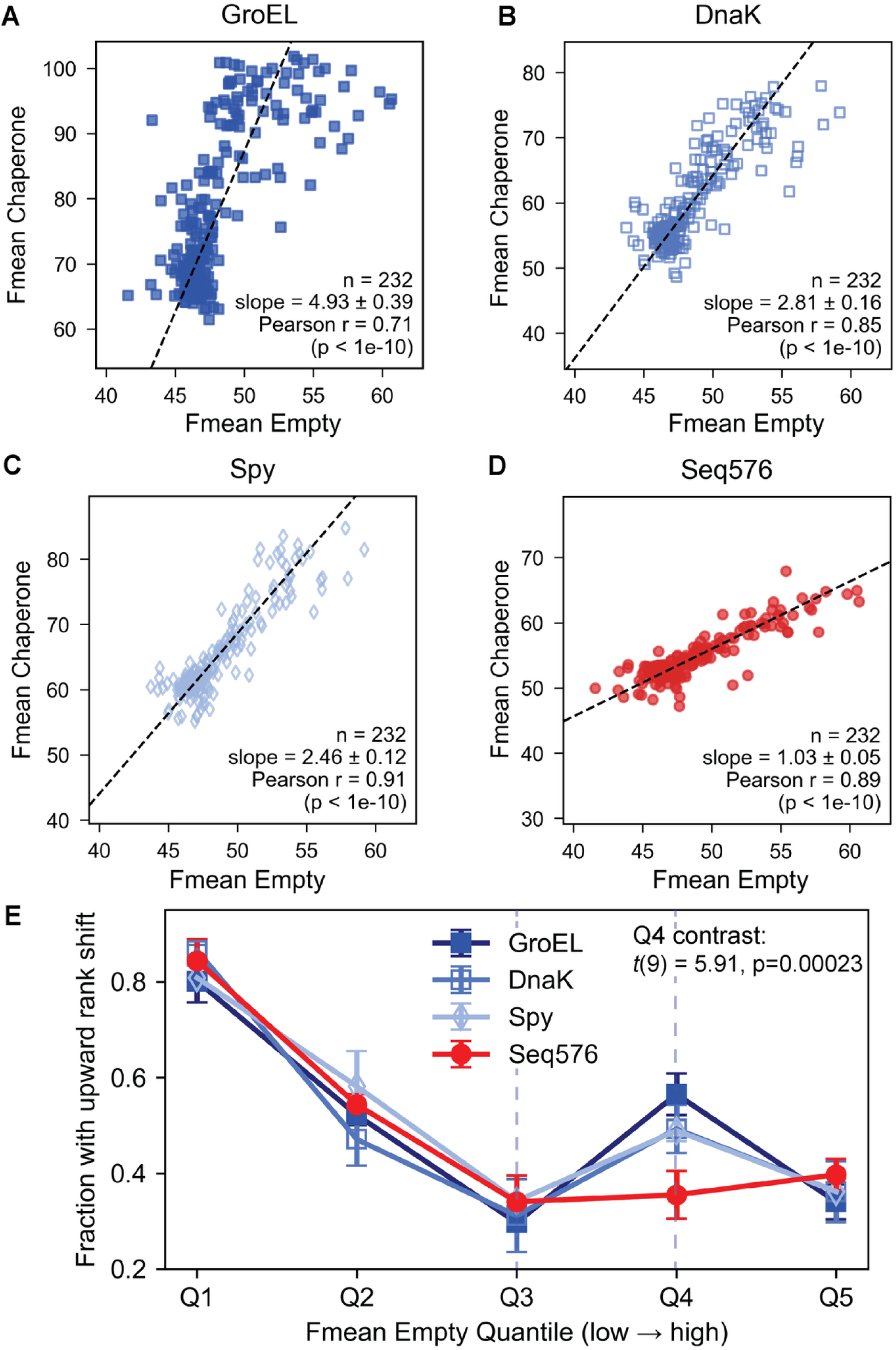
Stability-dependent selection and folding amplification landscape. (**A-D**) Relationship between baseline folding and chaperone-assisted folding. Scatter plots show Fmean values under each chaperone condition plotted against baseline folding levels measured under the Empty condition for GroEL (**A**), DnaK (**B**), Spy (**C**), and Seq576 (**D**). Protein chaperones exhibit slopes substantially greater than 1, indicating stronger folding enhancement for variants with higher baseline folding levels, whereas Seq576 shows a slope near 1, indicating minimal dependence on baseline stability. (**E**) Quantile-based rank-shift analysis of folding redistribution. Residues were grouped by baseline folding quantiles based on the Empty condition. Protein chaperones show a marked increase in upward redistribution between the Q3 (40^th^–60^th^ percentile) and Q4 (60^th^–80^th^ percentile) quantiles, consistent with a stability-dependent amplification effect, whereas Seq576 shows a more uniform redistribution pattern. At Q4, the contrast between protein-based chaperones and Seq576 was significant (*t*(9) = 5.91, *p* = 0.00023). Points represent means across replicates; error bars indicate SD.

To gain deeper insight into this selectivity, we next analyzed how residues changed their relative folding rank under each chaperone condition. Variants were first grouped into five quantiles based on their Fmean_Empty_ values, from lowest foldability (Q1) to highest foldability (Q5). For each biological replicate and chaperone condition, variants were re-ranked according to their Fmean values. We then calculated the fraction within each chaperone quantile that shifted to a higher relative folding rank than in the matched Empty condition.

This analysis revealed a distinct increase for the three protein-based chaperones: the fraction of residues showing upward rank shifts increased sharply from Q3 (40^th^–60^th^ percentile) to Q4 (60^th^–80^th^ percentile), whereas no comparable transition was observed for Seq576 (Fig. 5E). At Q4, the mean fraction under GroEL, DnaK, and Spy was significantly greater than that under Seq576 (difference = 0.161, 95% CI = 0.099–0.222, *t*(9) = 5.91, *p* = 0.00023). A higher-resolution ten-quantile analysis further showed that the overall redistribution profiles differed significantly among chaperone conditions (condition × quantile interaction, F = 2.45, p = 0.0011), and confirmed the difference between protein-based chaperones and Seq576 across Q7–Q8 (Holm-adjusted *p* = 0.00068; Fig. S22; Table S9). Thus, variants with intermediate-to-high baseline foldability were more likely to improve when additional protein-based chaperones were present, consistent with foldability-dependent amplification. In contrast, Seq576 produced a more even redistribution across quantiles, indicating little preference based on baseline foldability.

Predicted mutation stability from ThermoMPNN (33) did not correlate with chaperone-mediated folding rescue (Spearman ρ = –0.11 to 0.08). Because ThermoMPNN provides predicted rather than experimentally measured stability changes, the lack of correlation may partly reflect limitations in prediction accuracy. These results therefore do not exclude a contribution from thermodynamic stability but also suggest that predicted stability alone does not account for the observed chaperone responses (Fig. S23). Instead, cellular folding outcomes likely reflect combined contributions from thermodynamic stability, folding kinetics, and degradation (25, 34). Similarly, synonymous codon usage showed no detectable relationship with folding rescue (Fig. S24). Only weak positional trends were observed along the sequence, indicating minimal sequence-position effects overall (Fig. S25).

Together, these results reveal a “rich-getting-richer” pattern of foldability-dependent amplification by the three protein-based chaperones, whereas Seq576 acts more uniformly across variants with different baseline foldabilities. These patterns could not be explained by sequencing depth, codon usage, or predicted thermodynamic stability alone.

## Discussion

Together, the CHAP-SEQ data reveal both shared features among the protein-based chaperones and clear distinctions between protein– and RNA-based chaperone action. Structural and amino acid class analyses showed that the protein-based chaperones preferentially depended on buried hydrophobic residues within the protein interior. In contrast, Seq576 showed little overall preference for amino acid class or surface exposure, but instead displayed a localized dependence on the most dynamic C-terminal β12 region of the protein (Figs. 2 and 4). This observation is reminiscent of G4s’ broad binding ability with intrinsically disordered proteins (11, 35–37). Although β12 is structured in the mature protein, its elevated dynamics and position may render this region more accessible to Seq576 during folding. Therefore, at least for this client, G4 and protein-based chaperones appear to promote folding through largely complementary mechanisms in cells. Comparing the outliers among the protein-based chaperones further showed that despite their shared preference for hydrophobic core residues, they displayed distinct structural patterns of residue dependence.

A second major distinction is their preference based on client baseline foldability (Fig. 5). Previous models of genetic buffering generally propose that chaperones preferentially rescue poorly folded or deleterious variants, although this remains controversial (3). Surprisingly, in this work the protein-based chaperones examined here exhibited a “rich-getting-richer” pattern, preferentially amplifying variants with greater baseline foldability, whereas the RNA-based chaperone Seq576 acted more uniformly across variants. These results suggest that protein-based chaperones may most effectively promote folding of intermediates that are already competent to enter a productive folding pathway, rather than universally rescuing severely compromised variants. Variants with low baseline foldability may fail to reach a chaperone-accessible intermediate state, or may undergo irreversible misfolding and degradation. Further studies will be required to determine whether this foldability-dependent amplification extends to other client proteins and cellular systems.

The alanine-scanning design used here prioritized mechanistic interpretability and direct comparison across all chaperone conditions over maximizing the range of folding perturbations. However, broader mutagenesis strategies incorporating chemically diverse substitutions could reveal a wider spectrum of folding defects and chaperone responses. Similarly, the binary high– and low-fluorescence sorting strategy was selected for experimental simplicity and robust sample recovery but is not an inherent requirement of CHAP-SEQ. Future applications could incorporate multi-bin sorting to improve reconstruction of variant fluorescence distributions or focus on specific regions of the folding landscape. Because these conclusions are based on a single fluorescent reporter, they should not yet be generalized to all client proteins. However, CHAP-SEQ can be extended to diverse client proteins according to the experimental questions.

The distinct residue-level preferences of Seq576 and the protein-based chaperones, along with the foldability preference of chaperone proteins but not Seq576, raise potential evolutionary implications. In the context of the RNA world hypothesis (38–40), RNA-based chaperones likely pre-existed protein-based chaperones, and not just from quadruplexes; even the protein-folding activity of the ribosome has been attributed to RNA components (40). The results here suggest that in an RNA-peptide world, the RNA-based chaperones may have been able to provide relatively broad assistance for disordered peptides before the emergence of modern protein-based chaperone systems that favor clients capable of forming stable hydrophobic cores. This possibility is consistent with computational models linking protein evolution to folding stability, aggregation propensity, and chaperone activity (41, 42).

Taken together, CHAP-SEQ provides a flexible framework that can be adapted to different mutational libraries, sorting strategies, and client proteins to investigate diverse forms of chaperone-assisted folding.

## Materials and Methods

### Construction of expression vectors and alanine-scanning library

Expression vectors for molecular chaperones (GroEL, DnaK, and Spy), RNA G4 (Seq576), and corresponding empty controls were based on previously described pBAD33-derived backbones (10). Briefly, chaperone expression constructs were generated in the pBAD33 vector using SacI and HindIII restriction sites, and RNA expression constructs were cloned into the modified pBAD33mut backbone using SacI and SpeI sites, as described previously (10). pBAD/HisD-TagRFP675 was used as a folding biosensor (43). The reporter plasmid pBAD/HisD-TagRFP675 carries a pBR322 origin, whereas the pBAD33-derived protein-chaperone and RNA-expression plasmids carry a p15A origin. Only the indicated protein chaperone was plasmid-overexpressed; endogenous cochaperone systems, including GroES and DnaJ/GrpE, remained available in the host cells. A single-amino-acid alanine substitution library of TagRFP675 was newly generated by substituting each amino acid residue with alanine (GCG codon), spanning residues 2–233. The expression library was synthesized and pooled (GenScript). Strains and plasmids used in this study are listed in Table S1.

### Co-expression and fluorescence assay

Expression vectors encoding molecular chaperones and either pBAD/HisD-TagRFP675 (WT) or the pBAD/HisD-TagRFP675 mutation library were co-transformed into MC4100(DE3) (Δ*(argF-lac) U169 araD139 rpsLS50 reLAl deoCI ptsF25*) by electroporation. Transformants were selected on LB agar supplemented with ampicillin (200 μg/mL) and chloramphenicol (50 μg/mL). Protein and RNA expression was induced by plating cells on LB agar containing 0.2% L-arabinose and antibiotics, followed by overnight incubation at 42 °C. Non-induced control plates containing antibiotics only were used to confirm tight regulation of the pBAD promoter.

Cells were scraped from plates, resuspended in 170 mM NaCl, and harvested by centrifugation at 10,000 × g for 2 min at 4 °C. Cell pellets were imaged immediately after harvesting to qualitatively compare fluorescence. Cells were then resuspended in 170 mM NaCl and diluted to OD600 = 0.1 for bulk fluorescence measurements and preparation for cell sorting. Fluorescence emission was measured using a microplate reader (Infinite M200 Pro, Tecan) in black 96-well plates (Corning Black NBS 3991), with excitation at 598 nm and emission detection at 675 nm.

### Fluorescence-activated cell sorting (FACS)

For fluorescence-activated cell sorting (FACS), cell suspensions prepared at OD600 = 0.1 were further diluted 1:10 to a final OD600 of approximately 0.01, corresponding to ∼10^6^ cells/mL. At this dilution step, DAPI was added to assess cell viability, and parallel samples without DAPI were prepared as negative controls. Cells were analyzed and sorted on a Sony SH800 cell sorter (Sony Biotechnology, San Jose, CA) equipped with 405/488/561/638 nm lasers. DAPI fluorescence was detected in the FL1 channel using a 450/50 nm bandpass filter, and red fluorescent protein signals (TagRFP675/mPlum range) were collected in the FL4 channel using a 665/30-nm bandpass filter. Cell viability during sorting was assessed by DAPI staining, and cells exhibiting high DAPI fluorescence were excluded from all sorting gates.

Gating thresholds were defined using non-induced control cells. Viable cells were subsequently divided into low– and high-folding populations based on TagRFP675 fluorescence intensity. The low– and high-folding fractions were collected at approximately equal ratios, with each fraction typically representing 40–48% of viable cells after exclusion of DAPI-positive events. For each sample, approximately 100,000 events were acquired and analyzed using Cell Sorter Software (version 1.7, LE-SH800 series).

In Round 1, the TagRFP675 mutation library was sorted in the presence of Empty vector, GroEL, or Seq576. In Round 2, the same library was sorted in the presence of Empty vector, DnaK, or Spy. Three biological replicates were processed independently for each condition. After initial sorting, the low– and high-fluorescence populations were reanalyzed by FACS, and recovered plasmid libraries were re-transformed into fresh cells and re-evaluated by FACS and bulk fluorescence. Only libraries that retained the expected fluorescence phenotype were advanced to sequencing.

### NGS processing and inference of folding values

NGS was performed by Azenta Life Sciences using 2 × 150 bp paired-end Illumina sequencing. Sample-level read counts and sequencing-quality metrics are provided in Tables S3 and S4. To ensure robust alanine substitution quantification, a strict codon-level filtering strategy was applied (Fig. S4). Only variants containing complete and unambiguous GCG codon evidence across all required nucleotide positions were retained for downstream analyses.

To infer quantitative folding values from sorting data, fluorescence distributions were modeled using a log-normal framework (Fig. S5). For each variant, pHigh = *freq*_High_ / (*freq*_High_ + *freq*_Low_), where *freq*_High_ and *freq*_Low_ represent variant frequencies in the respective sorted populations. Mean fluorescence values (Fmean) were obtained by numerical inversion of the previously established log-normal gating model using pHigh, the shared gate threshold, and the empirically determined σ (15). Numerical stabilization and replicate normalization were applied prior to downstream analyses.

### Residue-level and structural analyses

Chaperone-dependent folding effects were quantified as log_2_ratioF = log_2_(Fmean_Chaperone_/Fmean_Empty_), using the corresponding Empty control from the same experimental round. Residue-level effects were analyzed using condition-normalized outlier comparisons, rank-based redistribution analyses, and sliding-window enrichment analyses. Structural visualization and mapping were performed using the TagRFP675 crystal structure (PDB: 4KGF). Additional structural analyses included AlphaFold3-based structure prediction (30), ensemble RMSD analysis (28, 29), and ThermoMPNN-based stability prediction (33).

## Statistical analyses

Statistical analyses were performed at the biological-replicate level, with methods selected according to the structure of each dataset and the specific hypothesis being tested. Statistical methods were selected according to the structure of each dataset and the specific hypothesis being tested. Analyses included correlation and regression models, ANOVA and planned contrasts, rank– and permutation-based tests, outlier detection, and enrichment analyses. Multiple-comparison corrections were applied where appropriate. Full model specifications, analysis units, thresholds, permutation procedures, and correction methods are provided in the Supporting Methods and corresponding figure legends.

## Computational analysis and code availability

Core CHAP-SEQ processing steps, including fluorescence inference modeling, numerical stabilization, and downstream residue-level analyses, were implemented in Python (v3.13) using NumPy, SciPy, and pandas. Analysis scripts and example datasets are available at https://github.com/AhhyunSon/CHAP-SEQ-analysis.

*Additional experimental procedures, equations, statistical analyses, and computational details are provided in the Supporting Information*.

## Author Contributions

Conceptualization: SH, AS

Methodology: SH, AS, TAW, CD

Investigation: AS

Visualization: AS

Funding acquisition: SH

Writing – original draft: SH, AS

Writing – review & editing: SH, AS, TAW, CD

## Classification

Biological Sciences; Biochemistry

