## Supplemental Information for "Distinct modes of folding assistance in cells by protein- and RNA-based chaperones"

PNAS

**This PDF file includes:**

Supporting text
Figures S1 to S25
Tables S1 to S9
Legends for Movies S1 to S16
Legends for Datasets S1 to S2
SI References

**Other supporting materials for this manuscript include the following:**

Movies S1 to S16
Datasets S1 to S2

### Supporting Information Text

### SI Results

#### DAPI-based assessment of cell viability during FACS sorting

Cell viability during FACS was monitored using DAPI staining, and cells exhibiting high DAPI fluorescence were excluded from all sorting gates. To assess whether chaperone expression or library complexity affected cell viability, DAPI fluorescence distributions were compared across multiple experimental conditions, including pBAD33 vector only controls, Wild-type (WT) TagRFP675 expressed with individual chaperones, and TagRFP675 alanine mutation libraries expressed with chaperones (Fig. S1).

Across all tested conditions, the fraction of cells falling into the high-DAPI population ranged from approximately 20% to 40%. Importantly, no substantial differences in DAPI-positive fractions were observed between empty vector controls and samples expressing chaperones, nor between WT TagRFP675 and alanine scanning libraries. These results indicate that neither chaperone expression nor mutational library expression led to a condition-specific increase in cell death during sorting, supporting the robustness and comparability of the FACS-based folding measurements.

#### Validation of FACS-based separation of sample selection

To verify robust separation of low- and high-folding populations and exclude phenotypes arising from chromosomal alterations, sorted populations were evaluated by cell-pellet appearance, bulk fluorescence measurements, and FACS reanalysis (Figs S2 and S3). In Round 1, WT TagRFP675 and alanine mutation libraries expressed with Empty, GroEL, or Seq576 were initially separated into approximately equal low- and high-fluorescence populations. Plasmids recovered from the sorted populations were re-transformed into fresh E. coli, and the resulting populations were re-evaluated by bulk fluorescence and FACS. High-folding populations consistently exhibited increased fluorescence and right-shifted FACS distributions relative to both the original mutation library and low-folding populations, whereas low-folding populations showed reduced fluorescence. These differences were retained through repeated sorting and re-transformation, supporting enrichment of plasmid-encoded folding phenotypes rather than chromosomal effects.

Round 2 libraries expressed with Empty, DnaK, or Spy were validated using the same cell-pellet, bulk-fluorescence, and FACS-based criteria. Sorted populations were first evaluated before re-transformation, after which plasmids from selected populations were re-transformed into fresh cells in two independent batches. Re-transformed populations retained clear separation between low- and high-folding fractions. Samples showing inconsistent bulk-fluorescence separation or discordant FACS population shifts were excluded from downstream analysis. Following quality control, three biological replicates were selected for each condition, except for Spy, for which four biological replicates were selected. Validation outcomes and the samples selected for sequencing are summarized in Table S2.

#### Sequencing yield and quality

Sequencing quality was assessed using the quality-control reports provided by Azenta Life Sciences. Vendor-provided sequencing quality metrics were examined for all libraries retained after FACS-based quality control (Table S3). The 18 Round 1 libraries generated a total of 85.96 million read pairs and 25.79 Gb of sequence. Individual libraries yielded 3.77–5.67 million read pairs, with an aggregate mean quality score of 34.90 and 88.07% of bases at or above Q30. The 20 Round 2 libraries generated a total of 53.05 million read pairs and 15.92 Gb of sequence. Individual libraries yielded 2.30–3.02 million read pairs, with an aggregate mean quality score of 37.84 and 89.37% of bases at or above Q30. These results indicate consistent sequencing yield and base-calling quality across the libraries used for downstream CHAP-SEQ analysis.

#### Sequencing depth stability and numerical robustness of Fmean inference

We evaluated sequencing-depth stability across rounds and conditions (Table S4). Depth ratios ($DP_H/DP_L$) were centered near unity in both rounds (median = 1.002 in R1; 0.999 in R2). The $5^{th}$–95th percentile ranges were 0.993–1.046 in R1; 0.911–1.189 in R2, indicating generally balanced sampling between high- and low-fluorescence bins. The fraction of variants with zero

effective depth (DP=0) varied among conditions and was higher in some chaperone backgrounds including GroEL in R1 and DnaK in R2. These variants were processed using the same numerical-stabilization procedure applied uniformly across conditions.
We next examined the effect of ε-based numerical stabilization on probability estimates (Tables S5–S6). Across rounds and conditions, stabilized and raw estimates were highly concordant (Spearman $\rho = 0.985–0.997$; Pearson $r = 0.973–0.985$). And mean absolute differences were small relative to the overall dynamic range. Sensitivity analysis across quantile thresholds demonstrated that Q5 provided an optimal balance between minimizing deviation from raw probabilities and improving replicate consistency. The Q5 threshold was therefore used for all downstream Fmean inference and validation analyses.

**Analytical consistency and replicate reproducibility of inferred Fmean values**

Using the selected Q5 stabilization threshold, we next evaluated analytical consistency of the inversion framework. Within each replicate, inferred Fmean values were compared to their corresponding pHigh estimates (Fig. S6). As expected from the analytical inversion of the log-normal gating model, Fmean exhibited a smooth nonlinear relationship with pHigh, with $R^2$ values ranging from 0.984 to 0.994 across conditions. The slight curvature reflects the exponential form of the inversion and is fully consistent with model expectations.

We then assessed biological reproducibility by performing pairwise comparisons of inferred Fmean values between replicates for each condition and round (Fig. S7). Strong linear agreement was observed across replicate pairs, with consistently high Pearson correlation coefficients across conditions. Linear regression analyses demonstrated near-proportional concordance between replicates, indicating that sequencing-based inference yields quantitatively stable and reproducible estimates of variant-level folding output.

Collectively, these results demonstrate that Fmean inference is analytically coherent and reproducible across biological replicates, supporting its use for downstream comparative analyses.

**Orthogonal validation of inferred Fmean values**

To independently validate CHAP-SEQ-inferred folding outputs, we generated 18 individual alanine substitution variants selected from low- and high-bin populations and quantified their bulk fluorescence using a plate reader under matched expression conditions (Fig. S8). Across chaperone conditions, plate reader fluorescence exhibited strong monotonic agreement with inferred Fmean values, with Spearman correlation coefficients ranging from $\rho = 0.85$ (Empty) to $\rho = 0.96$ (Spy) ($p \leq 10^{-5}$ for all conditions). Concordance was observed across mechanistically distinct folding environments, including GroEL ($\rho = 0.91$), DnaK ($\rho = 0.91$), Spy ($\rho = 0.96$), and Seq576 ($\rho = 0.85$). These results provide independent experimental support for the sequencing-based inference framework and indicate that inferred Fmean values capture relative differences in variant-level fluorescence output across diverse chaperone backgrounds.

While orthogonal plate reader measurements confirmed the accuracy of CHAP-SEQ-inferred Fmean values *in vivo*, they did not directly determine whether Seq576 modulate the intrinsic folding of TagRFP675, independently of cellular expression effects. To address this question, we performed *in vitro* refolding assays using chemically denatured TagRFP675 variants in the presence of the G4-forming oligonucleotide Seq576 or a non-G4 single-stranded DNA control Seq42 (Fig. S9). Under refolding conditions (0.5 μM protein; 1:2 protein:DNA molar ratio), Seq576 consistently enhanced native fluorescence recovery relative to control reactions. Increased recovery was observed for WT TagRFP675 as well as selected variants Q111A and L232A. These results demonstrate that Seq576 can directly enhance recovery of the native folded state *in vitro.* Importantly, the direction of refolding enhancement observed *in vitro* was consistent with folding trends predicted by cell sorting-based CHAP-SEQ analysis, in which these variants exhibited increased inferred Fmean values in the Seq576 background.

Together, the plate-reader and *in vitro* refolding results provide support for the biological relevance of the CHAP-SEQ-derived folding metric across independent experimental contexts.

**Pairwise comparison of residue-level $log_2$ratioF reveals condition-specific folding patterns**

To compare folding effects across chaperones on a residue-by-residue basis, we computed $log_2$ratioF values for each condition. Spearman correlation analysis revealed moderate to strong concordance among the protein-based chaperones but little correlation between Seq576 and the protein-based chaperones (Fig. 1E). Full pairwise scatter plots are shown in Supplementary Fig. S10A. Pearson correlation analysis similarly showed moderate to strong linear concordance among the protein-based chaperones ($r = 0.45–0.77$), whereas correlations between protein-based chaperones and Seq576 were near zero ($r = −0.058$ to 0.097; Fig. S10B). These complementary analyses support a distinct residue-level pattern for Seq576.

**Amino acid and structural determinants of chaperone-dependent folding**

To determine whether chaperone-dependent folding effects exhibit residue-type specificity, we examined the distribution of WT-oriented $log_2$ratioF values across amino acid categories and structural exposure classes (Fig. S11). Heatmap analysis revealed that protein-based chaperones showed a strong preference for buried hydrophobic and hydrophobic aromatic residues within β-barrel interior. In contrast, Seq576 showed no significant overall preference for amino acid class or buried versus surface-exposed residues, although specific category-exposure combinations exhibited condition-specific shifts in mean $log_2$ratioF values. Within loop regions, the relative Seq576 versus protein-based chaperone response varied across amino acid classes (Kruskal–Wallis $H = 11.43$, $p = 0.0435$; permutation $p = 0.0502$). Charged residues showed the largest shifts toward Seq576 dependence, and a follow-up comparison showed greater relative Seq576 dependence for charged loop residues than for non-charged loop residues (Mann–Whitney $p = 0.0021$; permutation $p = 0.0019$). Together, these analyses indicate that protein-based chaperones preferentially assist canonical core-stabilizing residue classes, whereas Seq576 exhibits a more context-dependent residue-level preference profile shaped by the local structural environment.

**Seq576-specific β12 hotspot and its association with structural dynamics**

To determine whether this apparent clustering reflected statistically significant sequence contiguity, we performed sliding-window enrichment and longest-run permutation analyses (Table S7). Using a lower 15% threshold, Seq576 displayed the longest six-residue consecutive run spanning residues 197–202), (permutation p-value = 0.002, 9 FDR-significant windows, and a contiguous enriched region spanning residues 193–210. In contrast, GroEL, DnaK, and Spy conditions showed only short runs (length = 3), no FDR-significant windows, and non-significant permutation p-values (~0.49), consistent with random expectation.

Residue-level RMSD analysis further revealed structural variability within the same β12 region (Fig. S14). Global correlation analysis showed modest relationships between RMSD values and folding phenotypes across conditions (Table S8A). However, across the tested thresholds, residues with high RMSD significantly overlapped with residues showing low Seq576 $log_2$ratioF values, whereas no comparable enrichment was observed for protein-based chaperones (Table S8B). At the representative 10% threshold, the overlap was significant by both Fisher's exact test ($p = 0.019$) and permutation testing ($p = 0.018$). Sliding-window analysis likewise showed that local RMSD was negatively correlated with mean Seq576 log2ratioF (Spearman $\rho = −0.386$, $p = 5.06 \times 10^{-9}$) and positively correlated with the local fraction of bottom-10% Seq576 log2ratioF residues ($\rho = 0.326$, $p = 1.07 \times 10^{-6}$; Table S8C). Together, these analyses associate the Seq576-specific 193–210 hotspot with elevated structural variability in the C-terminal β12 region.

**Follow-up characterization of R157A**

R157A exhibited strongly reduced fluorescence relative to WT TagRFP675 across the chaperone conditions tested (Fig. S15A, B). Although GroEL and DnaK increased R157A fluorescence to some extent, fluorescence remained substantially reduced compared with WT. Examination of the underlying CHAP-seq sorting distributions showed that R157A was consistently represented in the Low-fluorescence bin but was frequently absent or below the predefined detection threshold in the corresponding High-fluorescence bin under protein-chaperone conditions (Fig. S15C). Across GroEL, DnaK, and Spy, R157A was detected in all Low-bin samples (10/10) but in only 3 of 10 High-bin samples. High-bin detection was lower under protein-chaperone conditions than in Empty controls (3/10 versus 6/6; two-sided Fisher's exact test, $p = 0.011$). In particular, R157A was not detected in any of the three GroEL High-bin replicates.

Because R157A fluorescence was strongly reduced and therefore concentrated predominantly within the Low-fluorescence range, the two-bin CHAP-seq sorting design provided limited resolution for quantitatively distinguishing chaperone-dependent effects on this variant. Thus, these follow-up measurements are consistent with a strong folding defect associated with R157A but were not used for further quantitative interpretation of residue-specific chaperone dependence.

**Structural distribution and statistical identification of high-variability residues**

Residue-level variability (sd_log2) was first examined using a distribution-based stratification approach. Within each condition, residues were partitioned into five categories (QD1-QD5) based on the mean (μ) and standard deviation (σ) of the sd_log2 distribution, with QD5 defined as $SD \geq \mu + 2\sigma$. Structural mapping showed that QD5 residues were observed in both β-sheet and loop regions; however, they were more frequently located at structural peripheries, including upper loop regions, lower terminal loops, and β-strand edges, rather than within the buried β-sheet core (Fig. S16).

To determine whether elevated sd_log2 values represented disproportionately high variability relative to effect magnitude, we next modeled the relationship between $|log_2 ratioF|$ and sd_log2 using LOESS regression. Residues within the top 5% of positive SD residuals were classified as LOESS high-variability candidates. Independently, Mahalanobis distance analysis identified multivariate outliers in the joint distribution of ($|log_2 ratioF|$, sd_log2). The intersection of these two criteria yielded a limited subset of statistically robust high-variability residues in each condition (Fig. S17).

Structural mapping of combined outliers revealed spatially dispersed patterns rather than consistent clustering. Overlap across conditions was limited, and no shared high-SD hotspot was observed across all chaperone conditions.

**Robustness of baseline-foldability-dependent amplification**

To determine whether folding effects depend on baseline folding levels, residue-level $log_2$ratioF values were plotted against baseline fluorescence measured under the Empty condition (Fig. S18). Protein-based chaperones (GroEL, DnaK, and Spy) exhibited positive correlations between $log_2$ratioF and baseline folding levels, indicating greater folding gains for residues with higher baseline folding. In contrast, Seq576 displayed a weak negative relationship between $log_2$ratioF and baseline folding level. To determine whether the residual-based framework reflects the same folding signal captured by $log_2$ratioF, we further compared centered residual values with $log_2$ratioF across variants (Fig. S19). Consistently, strong positive correlations were observed for the protein-based chaperones GroEL, DnaK, and Spy, but not for Seq576.

To exclude sequencing-depth effects, we next examined the relationship between pooled sequencing read depth (AD_sum_pooled) and deviations from the Empty baseline regression. Across all conditions, centered residuals exhibited no systematic correlation with pooled sequencing read depth (AD_sum_pooled), indicating that the observed dispersion was unlikely to arise from sequencing-coverage differences (Fig. S20). We next examined whether deviations from the Empty baseline model depend on the baseline folding level of each variant. Although statistically significant positive correlations were observed across conditions ($p < 1 \times 10^{-10}$), the slopes were small, indicating that baseline folding explains only a minor fraction of the residual variance (Fig. S21). These results suggest that baseline foldability alone does not account for the observed residual variation.

In Fig. 5B, quintile-based redistribution analysis revealed a marked increase between Q3 ($40^{th}$–$60^{th}$ percentile) and Q4 ($60^{th}$–$80^{th}$ percentile) for GroEL (+0.123), DnaK (+0.127), and Spy (+0.145), compared with Seq576 (+0.051), corresponding to an approximately 2.5 to 3-fold stronger amplification (Table S9). Increasing the percentile resolution from quintiles to deciles confirmed that this amplification was concentrated within the $60^{th}$–$80^{th}$ percentile range for protein-based chaperones, whereas Seq576 displayed a more gradual redistribution pattern without a distinct mid-high spike (Fig. S22). Exclusion of 10 variants that remained in the extreme baseline bins (Q1 or Q5) with minimal redistribution did not appreciably alter the redistribution patterns. Together, these sensitivity analyses support a baseline-foldability-dependent amplification pattern specific to the protein-based chaperones.

**Predicted stability does not explain folding rescue**

To test whether the observed chaperone-dependent folding effects could be explained by mutation-induced stability changes, ThermoMPNN-predicted ΔΔG values were compared with experimentally inferred folding phenotypes (Fig. S23). Across residues, predicted ΔΔG values showed little correlation with $log_2$ratioF measurements. These results indicate that the predicted thermostability alone does explain the folding effects observed in the CHAP-SEQ assay.

**Codon usage does not explain residue-level chaperone effects**

Local codon usage showed no detectable relationship with residue-level rescue for any chaperone condition (Fig. S24A–D). Spearman correlations between local codon frequency and $log_2$ratioF were close to zero and not statistically significant. Likewise, native alanine positions encoded by GCC and GCT showed no systematic differences in rescue values (Fig. S24E). Together, these analyses indicate that the observed residue-level rescue patterns are unlikely to arise from local codon usage effects.

**Weak condition-specific trends along the protein sequence**

To examine whether chaperone effects vary along the polypeptide sequence, we analyzed the relationship between residue position and chaperone-dependent folding effects ($log_2$ratioF) (Fig. S25). GroEL and DnaK showed no significant positional relationship (GroEL: ρ = -0.03, p = 0.70; DnaK: ρ = 0.07, p = 0.32), whereas Spy showed a weak positive correlation (ρ = 0.18, p = 0.005) and Seq576 a weak negative correlation ($\rho$ = -0.16, $p$ = 0.017). LOWESS smoothing showed gradual positional trends consistent with these correlations. Overall, sequence position showed only weak associations with chaperone-dependent folding effects.

### SI Materials and Methods

#### Post-sorting library processing, quality control, and sequencing

Sorted low- and high-folding populations were plated on LB agar containing ampicillin and chloramphenicol and incubated overnight at 37 °C. Cells were scraped, pooled, and stored as glycerol stocks (15 v/v% glycerol) at -80 °C until use.

For plasmid preparation of the sorted libraries, frozen stocks were plated onto five LB agar plates per condition to ensure sufficient cell numbers and library coverage. Plasmid DNA was purified from pooled cells using standard midi-prep procedures (Promega), yielding DNA of sufficient purity and quantity for downstream next-generation sequencing (NGS). Purified plasmid libraries were re-transformed into fresh *E. coli* MC4100(DE3) cells by electroporation, and re-transformed populations were expanded on at least three independent plates per condition.

To verify accurate separation of low- and high-folding populations and to exclude confounding effects such as chromosomal mutations, re-transformed libraries were subjected to additional fluorescence-based quality control (QC) analyses. Cell pellet color, bulk fluorescence intensity, and FACS profiles were compared across the original mutation library, sorted populations, and re-transformed cells. Only samples showing consistent folding-dependent behavior across all QC metrics were advanced to sequencing. Samples failing QC in Round 2 are indicated as red squares in the corresponding figure (Fig. S3F, S3U, S3Z, and S3AD). All QC-passing samples were analyzed by NGS with three biological replicates per condition, except Spy (n=4).

#### NGS and sequencing quality assessment

Selected high- and low-fluorescence populations were submitted to Azenta Life Sciences for next-generation sequencing. Round 1 libraries were sequenced on an Illumina MiSeq instrument, whereas Round 2 libraries were sequenced using an Illumina platform; both rounds used 2 × 150-bp paired-end sequencing. Vendor-provided quality-control metrics included read count, sequence yield, mean quality score, and the percentage of bases with a Phred quality score of at least 30 (Q30). Sequencing-quality summaries were compiled for the libraries retained for downstream CHAP-SEQ analysis, including three biological replicates per condition for Empty, GroEL, Seq576, and DnaK, and four biological replicates for Spy, with high- and low-fluorescence bins sequenced separately. Aggregate quality metrics were calculated across the listed samples using sequence yield as the weighting factor.

#### NGS processing and strict variant filtering for alanine scanning

Raw next-generation sequencing (NGS) reads were processed to quantify variant frequencies for alanine (GCG) scanning libraries. For each variant, read depth (DP) and variant-supporting read counts (AD) were extracted, where DP denotes the total number of reads covering a given codon position and AD denotes the number of reads supporting the alanine (GCG) variant. Variant allele frequency (*freq*) was calculated as *freq = AD / DP*.

To ensure robust and unambiguous quantification of alanine substitutions, a strict codon-level filtering strategy was applied. For each variant, codon evaluation was restricted to the codon positions required to encode the alanine (GCG) substitution (*need_idx*), which corresponds to all mutated positions in the alanine scanning library. Variant validity was determined according to the following decision logic. If any codon position within *need_idx* lacked positional coverage, the variant was considered unobserved, and both AD and DP were set to zero (flag: *missing_position*). If coverage was present but evidence for the complete GCG codon was incomplete across the required positions, AD and DP were likewise set to zero (flag: *no_gcg_evidence*). Because alanine substitutions in this library are uniquely encoded by the GCG codon, only variants with complete and consistent GCG evidence across all required positions were considered valid; variants with partial or inconsistent codon evidence were treated as unobserved (See Fig. S4).

For variants passing all validity criteria, the raw alanine read count ($AD^{raw}$) was calculated as the sum of AD across all positions in *need_idx*. For frequency-based analyses, AD and DP were computed as the mean values across *need_idx*, thereby normalizing for the number of nucleotide substitutions required to convert the original codon to the alanine (GCG) (n = 1–3, depending on the original codon sequence). This approach ensured consistent treatment of variants with differing codon requirements while preserving $AD^{raw}$ as an inspection metric.

The output of this strict filtering pipeline was a curated variant table containing $AD^{raw}$, AD, DP, variant frequency, and explicit flags indicating the reason for exclusion or acceptance of each variant. This filtered dataset was used for all downstream analyses.

**Inference of mean fluorescence (Fmean) from sorting fractions**

To infer mean fluorescence values from sequencing-based sorting data, we modeled single-variant cellular fluorescence as a log-normal distribution, consistent with established flow cytometry measurements of clonal populations (1). Under this model, the probability that a cell expressing variant *i* exceeds the fluorescence gate boundary $F_g$ is determined by the mean fluorescence $F_i$ and the log-normal width parameter σ.

Following the formulation introduced in a previous study (Eq. 2, (2)), the probability that fluorescence exceeds the gate threshold is given by:

$$P(F \geq F_g \mid F_i, \sigma) = \frac{1}{2} - \frac{1}{2}\mathrm{erf}\left(\frac{\ln F_g - \ln F_i + \frac{1}{2}\sigma^2}{\sigma\sqrt{2}}\right) \quad \text{(Eq. 1)}$$

where fluorescence values are assumed to follow a unimodal log-normal distribution and denote the Gaussian error function.

**1) Derivation of pHigh from NGS-derived bin counts.**

For each variant *i* in biological replicate *r*, effective read counts in the high- and low-fluorescence bins were obtained after strict codon-level filtering, yielding $AD^{\mathrm{eff}}_{H,i,r}$ and $AD^{\mathrm{eff}}_{L,i,r}$, with corresponding effective depths $DP^{\mathrm{eff}}_{H,i,r}$ and $DP^{\mathrm{eff}}_{L,i,r}$. Bin frequencies were defined as:

$$freq_{H,i,r} = \frac{AD^{\mathrm{eff}}_{H,i,r}}{DP^{\mathrm{eff}}_{H,i,r}}, \quad freq_{L,i,r} = \frac{AD^{\mathrm{eff}}_{L,i,r}}{DP^{\mathrm{eff}}_{L,i,r}} \quad \text{(Eq. 2)}$$

The raw pHigh (high-bin probability) was computed as:

$$\mathrm{pHigh}^{\mathrm{raw}}_{i,r} = \frac{freq_{H,i,r}}{freq_{H,i,r} + freq_{L,i,r}} \quad \text{(Eq. 3)}$$

Variants with no detectable bin signal (*i.e.*, $\boldsymbol{freq_{H,i,r}} = \boldsymbol{freq_{L,i,r}} = \mathbf{0}$) yield an undefined ratio under Eq. (3) and were subsequently handled through Quantile-based numerical stabilization prior to inversion, as described below

**2) Quantile-based numerical stabilization of bin probabilities.**

To prevent numerical divergence during inversion of the log-normal gating model, we implemented a round-level, distribution-adaptive ε stabilization. For each round, a small constant e was defined as:

$$\mathrm{e} = \frac{1}{2} \cdot Q_q(freq_H + freq_L) \quad \text{(Eq. 4)}$$

where $Q_q$ denotes the q-th percentile (q = 1.25, 2.5, 5, 7.5, or 10) of non-zero values of ($freq_H$ + $freq_L$) across all variants within that round.

To evaluate the impact of ε magnitude on inference robustness, we systematically compared stabilized and raw pHigh values across quantile thresholds. Concordance was assessed using Spearman and Pearson correlations, as well as mean and maximum absolute differences. In addition, replicate-level correlations were calculated for each threshold to evaluate improvements in biological reproducibility. Based on the balance between minimizing deviation from raw probabilities and maximizing replicate consistency, the $Q_5$ was selected as optimal stabilization thresholds for downstream analyses.

The stabilized probability was then computed as:

$$\mathrm{pHigh}^{\mathrm{stabilized}}_{i,r} = \frac{freq_{H,i,r} + 1\mathrm{e}}{freq_{H,i,r} + freq_{L,i,r} + 2\mathrm{e}} \quad \text{(Eq. 5)}$$

thereby ensuring finite values for numerical inversion under the log-normal gating model.

**3) Estimation of σ from FACS statistics.**

The log-normal width parameter σ was estimated directly from fluorescence distribution statistics obtained by flow cytometry. σ was calculated analytically from the measured full width at half maximum (FWHM) and the mode of each fluorescence histogram (Fig. S5).

$$\sigma = \ln\left(\frac{\frac{\mathrm{FWHM}}{\mathrm{mode}(X)}+\sqrt{\left(\frac{\mathrm{FWHM}}{\mathrm{mode}(X)}\right)^2+4}}{\sqrt{2\ln 2}}\right) \quad \text{(Eq. 6)}$$

Fluorescence distributions were measured under identical instrument settings using representative reference variants, including a low-bin mutant (L110A), a high-bin mutant (S28A), and WT TagRFP675 co-expressed with chaperones. Sixteen reference variants were measured in triplicate (48 total fluorescence distributions). σ was calculated independently for each replicate distribution and subsequently averaged per variant. Variant-level σ estimates had a median of 0.23 (mean = 0.29, SD = 0.30). The median σ (0.23) was used as the global width parameter for downstream inference.

**4) Inference of mean fluorescence (Fmean).**

Rearranging the expression for $P(F \geq F_g \mid F_i, \sigma)$ yields the explicit solution for the mean fluorescence:

$$F_{mean,i,r} = F_g \exp\left(\frac{1}{2}\sigma^2 - \sigma\sqrt{2}\,\mathrm{erf}^{-1}\left(1-2\mathrm{pHigh}_{i,r}^{stabilized}\right)\right) \quad \text{(Eq. 7)}$$

This equation was used to infer replicate-level mean fluorescence values for each variant under each experimental condition. Note that absolute fluorescence values can vary across experiments; downstream analyses therefore rely on ratio-based metrics ($\log_2$ratioF), and median scaling is applied when comparing Fmean values across rounds, as described below.

**Statistical validation**

Using the selected $Q_5$ stabilization threshold, we next assessed robustness and reproducibility of the inference pipeline.

Depth ratios ($DP_H/DP_L$) were calculated for each variant within each replicate and summarized by round. The fraction of variants with zero effective depth (DP=0) was reported for each round-condition combination. Robustness of pHigh calculation was assessed by comparing frequency-based and count-based estimates within each round.

To assess quantitative consistency of the inferred folding metric, Fmean values were compared with their corresponding pHigh values within each replicate. Linear regression was used as a descriptive measure of concordance, and Pearson's $R^2$ values were calculated.

Replicate-level reproducibility was evaluated using inferred Fmean values for all variants retained after strict filtering and stabilization. For each condition and round, all pairwise replicate comparisons were performed, and Pearson correlation coefficients were calculated to assess linear agreement.

**Orthogonal validation of inferred Fmean values**

To orthogonally validate CHAP-SEQ-inferred Fmean values, eighteen individual alanine substitution variants (H10A, N11A, N20A, S28A, G37A, S66A, Y96A, D98A, A104A, T108A, L110A, Q111A, C114A, K120A, S128A, N158A, R197A, and L232A) were generated in the pBAD/HisD-TagRFP675 plasmid by site-directed mutagenesis (QuikChange II, Agilent). These variants were selected from low-bin and high-bin populations identified during CHAP-SEQ selection. Co-expression with chaperones, induction conditions, and fluorescence measurements were performed as described above. Measurements were carried out in biological triplicate (Spy in quadruplicate). Bulk fluorescence values were compared to inferred Fmean values for the corresponding variants and conditions, and correlations were assessed using Spearman rank correlation coefficients.

***In vitro* unfolding and refolding assays**

Expression and purification of TagRFP675 were performed as previously described (3). Briefly, wild-type TagRFP675, TagRFP675(Q111A) and TagRFP675(L232A) were expressed in E. coli BL21(DE3), induced with 0.2% *L*-arabinose, and purified using Bio-Scale Mini Nuvia IMAC Ni-Charged column (Bio-Rad). Protein purity was confirmed by SDS-PAGE, and concentration was determined by both gel quantification and spectrophotometrically.

The G4-forming single-strand DNA oligonucleotide Seq576 (5′-TGTCGGGCGGGGAGGGGGGG-3′) was prepared. Oligonucleotides were annealed in 10 mM potassium phosphate buffer (pH 7.5) by heating to 95 °C followed by slow cooling. A non-G4-forming single-stranded DNA control (Seq42, 5′-AACGAAAGAACATAATCTCG-3′) was prepared under identical buffer conditions.

Chemical unfolding and refolding assays were conducted as previously described (Huang et al., 2025). TagRFP675 was denatured in 6 M guanidine hydrochloride and diluted into refolding buffer at 25 °C. Final concentration of TagRFP675 was 0.5 μM and residual Gu-HCl concentrations were <20 mM. Seq576 was pre-equilibrated in the cuvette prior to protein injection at 1:2 molar ratios (protein:DNA). Fluorescence recovery was monitored at excitation/emission 598/675 nm using an Agilent Cary Eclipse spectrophotometer. Control experiments were performed with Seq42 under identical conditions. All experiments were performed at least in duplicate.

**Calculation of chaperone-dependent folding effect (log$_2$ratioF)**

To quantify the magnitude of folding enhancement relative to baseline expression, we computed the fluorescence ratio for each residue:

$$ratioF_{chap} = \frac{Fmean_{chap}}{Fmean_{Empty}} \quad \text{(Eq. 8)}$$

and expressed the effect size on a symmetric scale:

$$log2ratioF_{chap} = \log_2\left(\frac{Fmean_{chap}}{Fmean_{Empty}}\right) \quad \text{(Eq. 9)}$$

This metric quantifies the relative folding response with chaperone for each mutant compared to its baseline folding level and enables direct comparison of condition-dependent effects across residues. Positive log$_2$ratioF values indicate increased folding relative to the Empty control, whereas values near zero indicate minimal deviation from baseline.

For analyses performed within each experimental round, log$_2$ratioF values were computed directly from the inferred Fmean values without additional scaling. For cross-round comparisons (including global correlation analyses), Round 2 Fmean values were median-scaled to match the Empty baseline of Round 1 prior to ratio calculation. Specifically, a scaling factor was defined as the ratio of the median Fmean$_{\text{Empty}}$ values between Round 1 and Round 2:

$$\alpha = \frac{\mathrm{median}(F_{\text{Empty}}^{\text{Round 1}})}{\mathrm{median}(F_{\text{Empty}}^{\text{Round 2}})} \quad \text{(Eq. 10)}$$

and all Fmean values in Round 2 were multiplied by this factor prior to ratio calculation:

$$F_{\text{mean}}^{\text{Round 2, scaled}} = \alpha \cdot F_{\text{mean}}^{\text{Round 2}} \quad \text{(Eq. 11)}$$

This alignment was applied exclusively for cross-round comparisons; all within-round analyses were performed using unscaled Fmean values.

**Structural projection of residue-level effects**

Residue-level log$_2$ratioF values were projected onto the crystal structure of TagRFP675 (PDB ID: 4KGF). Only chain A was retained for visualization to avoid redundancy from symmetric chains. Residues outside the analyzed range (2–233) were excluded.

To preserve directional information while maintaining visual contrast, values were scaled independently within each condition using a piecewise linear transformation centered at the mean, such that the minimum, mean, and maximum values were mapped to -1, 0, and +1, respectively. Values below and above the mean were scaled separately to preserve directional differences. Thus, color intensity reflects relative deviation from the mean within each condition rather than absolute magnitude comparisons across conditions.

The resulting modified PDB files were visualized in ChimeraX (4) using continuous color gradients to represent effect magnitude, while sphere overlays were used to encode propagated variability (sd_log2) or statistically defined outlier categories where applicable. This approach enabled structural coloring proportional to residue-level folding effects, allowing qualitative comparison of spatial patterns between chaperone conditions.

**Condition-normalized identification of chaperone-specific and concordant outliers**

To compare residue-level folding-effect patterns across chaperone conditions with different log$_2$ratioF distributions, log$_2$ratioF values were standardized separately within each condition by z-score normalization. Residues with negative z-scores were classified according to effect-score (ES) categories: ES3 ($-2 < z \leq -1$), ES4 ($-3 < z \leq -2$), and ES5 ($z \leq -3$). These categories identify residues showing relatively low $\log_2$ratioF values within each condition.

Overlap of ES3–ES5 residues was then compared across conditions to identify chaperone-specific and concordant patterns. Seq576-specific residues were defined as ES3–ES5 residues identified for Seq576 but not for any of the three protein-based chaperones. Protein-based-chaperone-specific residues were defined as ES3–ES5 residues identified in at least one protein-based chaperone condition but not in Seq576. Among the protein-based chaperones, GroEL-, DnaK-, and Spy-specific residues were defined according to condition-specific ES3–ES5 membership, whereas concordant residues were defined as residues classified as ES3–ES5 in all three protein-based chaperone conditions.

For structural visualization, ES categories were encoded by sphere size, with ES5 residues displayed as the largest spheres, ES4 residues as intermediate-sized spheres, and ES3 residues as the smallest spheres. Residues were mapped onto the TagRFP675 structure (PDB ID: 4KGF), with sphere size indicating ES category.

Because R157 was the only residue classified as an ES3–ES5 outlier across all three protein-based chaperones, R157A was selected for follow-up characterization. R157A fluorescence was measured under Empty and chaperone conditions using the same reporter-fluorescence assay described above and compared with WT TagRFP675. In addition, R157A variant frequencies were extracted from the processed CHAP-seq datasets separately for the Low- and High-fluorescence bins of each replicate. Samples in which the expected alanine-encoding GCG sequence at residue 157 did not meet the predefined detection criteria were classified as *missing_position* and assigned a variant frequency of zero, consistent with the CHAP-seq processing pipeline. Replicate frequencies are shown as individual values with mean ± SD. For a secondary analysis of High-bin detection, R157A was classified as detected or below threshold in each sample, and detection frequencies were compared between Empty and pooled protein-based-chaperone conditions (GroEL, DnaK, and Spy) using a two-sided Fisher's exact test.

**Replicate normalization and uncertainty estimation**

Prior to residue-level averaging, replicate-specific intensity drift was corrected within each condition. For each replicate, fluorescence values were scaled by a factor equal to the ratio of the condition-level median fluorescence to the replicate median fluorescence. This normalization preserved condition-dependent differences while correcting for replicate-to-replicate variation.

Residue-level mean fluorescence (Fmean) and standard deviation (Fsd) were then calculated across normalized replicates.

Uncertainty in $\log_2$ratioF was estimated by analytical error propagation, yielding the propagated standard deviation sd_log2 for each residue.

For $x = \text{Fmean}_{\text{Chap}}$, $y = \text{Fmean}_{\text{Empty}}$, the propagated uncertainty of the log2 ratio was calculated as:

$$sd\_log2 = \sqrt{\left(\frac{Fsd_{chap}}{x\ln 2}\right)^2 + \left(\frac{Fsd_{Empty}}{y\ln 2}\right)^2} \qquad \text{(Eq. 12)}$$

where $\text{Fsd}_{\text{Chap}}$ and $\text{Fsd}_{\text{Empty}}$ represent the replicate-level standard deviations of the respective conditions.

**Structural visualization of residue-level variability**

To visualize the spatial distribution of residue-level variability in $\log_2$ratioF, two complementary representations were generated.

**1) SD distribution-based structural mapping.**

Residue-level standard deviations of $\log_2$ratioF (sd_log2) were multiplied by a constant scaling factor for visualization and partitioned into five groups (QD1–QD5) based on mean (μ) and standard deviation (σ) of the sd_log2 distribution within each condition.

Residues were assigned as follows:

- QD1: $SD < \mu - \sigma$
- QD2: $\mu - \sigma \leq SD < \mu$

- QD3: $\mu \leq SD < \mu + \sigma$
- QD4: $\mu + \sigma \leq SD < \mu + 2\sigma$
- QD5: $SD \geq \mu + 2\sigma$

These categories were encoded as sphere size, enabling simultaneous visualization of folding effect magnitude and replicate variability across the protein structure.

**2) Statistical identification of high-variability outliers.**

To identify residues exhibiting unusually high variability relative to their effect size, we modeled the relationship between $|\log_2 \text{ratioF}|$ and sd_log2 using LOESS regression (5) (smoothing fraction = 0.3).

For each residue, an SD residual was defined as:

$$sd_{residual} = sd_{log2} - sd_{expected,LOESS} \quad \text{(Eq. 13)}$$

Residues within the top 5% of SD residuals were classified as LOESS high-variability candidates.

Independently, Mahalanobis distances (6) were computed from the bivariate distribution of ($|\log_2 \text{ratioF}|$ , sd_log2) using the empirical mean vector and covariance matrix. Residues exceeding the $\chi^2$ threshold (df = 2, p = 0.975) were classified as multivariate outliers.

For structural visualization, residues were categorized into three levels:

- Combined outliers (LOESS ∩ Mahalanobis): displayed with largest sphere size
- LOESS-only outliers: intermediate size
- Mahalanobis-only outliers: smaller size

This hierarchical encoding distinguished globally elevated variability from statistically anomalous variability relative to effect magnitude.

**Sliding-window enrichment and permutation-based hotspot analysis**

To determine whether residues exhibiting strong chaperone-dependent effects cluster along the primary sequence, sliding-window enrichment analysis was performed on residue-level $\log_2$ratioF values. For each condition, strong-effect residues were defined across multiple percentile thresholds corresponding to the lower 5%, 10%, 15%, 20%, and 25% of the $\log_2$ratioF distribution to assess the robustness of sequence clustering (Table S7). For each threshold, sliding-window enrichment was evaluated using a window size of 10 residues, and statistical significance was assessed using a binomial survival test based on the global frequency of strong residues, followed by Benjamini-Hochberg correction (FDR < 0.05). In parallel, the longest consecutive run of strong residues was compared to a null distribution generated from 10,000 random permutations of residue labels. All analyses were performed independently for each condition. The 15% threshold was selected as the representative cutoff.

**Structure prediction using AlphaFold3**

Protein-nucleic acid complex structures were predicted using AlphaFold3 (7). The amino acid sequence of TagRFP675 and the DNA sequence corresponding to Seq576 were used as inputs. Default parameters were applied unless otherwise specified. Predicted structures were ranked based on model confidence metrics, including predicted local distance difference test (pLDDT) and interface predicted TM-score (ipTM). The top-ranked model was selected for visualization and further analysis.

**Ensemble refinement and root-mean-square deviation (RMSD) analysis of structural dynamics**

Visual inspection of the 4KGF crystal structure and maps suggested considerable unmodeled structural heterogeneity. Therefore, re-refinement was performed in Phenix using both single-model and ensemble refinement for comparison and otherwise identical parameters (8, 9). Single-model refinement produced Rwork/Rfree of 0.22/0.27, and ensemble refinement produced Rwork/RFree of 0.21/0.26, both with bond RMSD of 0.007 Å, but with ensemble refinement having an improved angle RMSD of 0.97° as opposed to 1.0° for the single model refinement. Together with improved fit to map by visual inspection, these metrics show that the multi-model refinement is more appropriate for this structure.

Residue-level structural dynamics were calculated in ChimeraX (4) using RMSD values calculated from the ensemble refinement. To evaluate relationships between structural dynamics and folding phenotypes, Spearman and Pearson correlations were calculated between residue RMSD values and experimentally inferred folding metrics, including baseline folding levels (Fmean) and chaperone-dependent folding effects ($log_2$ratioF).

To determine whether structurally dynamic residues overlap with residues showing strong chaperone-dependent folding effects, enrichment analyses were performed comparing residues within the top 5%, 10%, 15%, 20%, and 25% RMSD levels to residues within the corresponding bottom percentiles of $log_2$ratioF for each condition. Overlap significance was evaluated using Fisher's exact test and permutation testing. The 10% threshold was selected as the representative cutoff for subsequent analysis.

Spatial correspondence between structural variability and chaperone-dependent folding effects was further examined using a sliding-window analysis with a window size of 10 residues. For each window, mean RMSD and mean $log_2$ratioF were calculated, together with the fraction of residues belonging to top 10% of RMSD values and the fraction belonging to the bottom 10% of $log_2$ratioF values. Spearman correlations were calculated between mean RMSD and mean $log_2$ratioF and, separately, between frac_RMSD_top10 and frac_log2_bottom10 for each condition.

Structural visualization was performed by mapping RMSD values onto the wild-type TagRFP675 protein structure. Residues with high RMSD values were visualized using ensemble representations to illustrate local conformational variability.

**Residue-level category analysis**

To investigate whether chaperone-dependent folding effects exhibit systematic preferences with respect to amino acid properties and structural context, residue-level $log_2$ratioF values were analyzed. Each residue was annotated according to its physicochemical amino acid category and structural exposure class. Amino acid categories were defined as hydrophobic (A, V, L, I, M), hydrophobic aromatic (F, W, Y), polar neutral (S, T, N, Q, C), charged basic (K, R, H), charged acidic (D, E), and glycine/proline (G, P). Structural exposure classes were defined based on solvent accessibility and β-barrel topology as buried (In), surface-exposed (Out), loop, both (positions exhibiting mixed characteristics), and glycine-specific classification where applicable.

For each chaperone condition, the distribution of $log_2$ratioF values across categories was visualized using violin plots with overlaid boxplots indicating medians and interquartile ranges. Median values were color-encoded using a continuous green gradient, with darker tones corresponding to lower $log_2$ratioF values. Statistical comparisons between categories were performed using Welch's t-test to account for unequal variances. Resulting p-values were adjusted for multiple testing using the Benjamini-Hochberg procedure to control the false discovery rate. All statistical analyses were conducted independently for each chaperone condition.

To assess whether amino acid identity and structural exposure jointly influence chaperone-dependent folding effects, residues were further grouped according to the combination of amino acid category and exposure class. For each combination, the mean $log_2$ratioF value was calculated and visualized as a heatmap. Color intensity represents the mean $log_2$ratioF value within each group, with darker colors corresponding to lower $log_2$ratioF values. The number of residues included in each category combination is indicated within each cell.

To further quantify amino-acid-class dependence within loop residues, $log_2$ratioF values were standardized separately within each chaperone condition among residues classified as loops. For residues with complete measurements across all four conditions, the mean standardized response of the three protein-based chaperones (GroEL, DnaK, and Spy) was calculated and subtracted from the standardized Seq576 response. Thus, negative values indicate relatively greater Seq576 dependence, whereas positive values indicate relatively greater protein-chaperone dependence. Differences in this Seq576-versus-protein-chaperone response among the six amino acid classes were evaluated using a Kruskal–Wallis test and a permutation test in which amino-acid-class labels were randomly permuted among loop residues (100,000 permutations). As a follow-up to the pattern observed in the amino-acid-class/structural-context heatmap, charged residues, comprising the basic and acidic classes, were compared with the remaining non-charged loop residues using a two-sided Mann–Whitney U test and a two-sided permutation test of the difference in group means (100,000 permutations).

**Comparison of folding outcomes across chaperone conditions**

To compare folding outcomes across conditions, mean fluorescence values (Fmean) were analyzed at the residue level. For each variant, Fmean measured under each chaperone condition (GroEL, DnaK, Spy, and Seq576) was compared with the corresponding baseline fluorescence measured under the Empty control. Scatter plots were generated by plotting Fmean under each chaperone condition against Fmean measured under the corresponding Empty control. Variants corresponding to residues 2-233 were included in the analysis.

To quantify the scaling relationship between baseline folding and folding outcomes under each condition, regression slopes were estimated using Deming regression, which accounts for measurement error in both variables. Deming regression was implemented assuming equal error variance between variables ($\lambda = 1$). To estimate uncertainty in the slope parameter, bootstrap resampling was performed (1000 resamples), and the standard error of the slope was calculated from the bootstrap distribution. Pearson correlation coefficients were also computed to quantify the strength of the linear relationship between baseline folding and folding outcomes.

For visualization, axis ranges were standardized across panels to enable direct comparison of slopes and dispersion between chaperone systems.

**Baseline-foldability dependence of residue-level folding effects**

To examine whether chaperone-dependent folding effects depend on baseline folding levels, residue-level $\log_2$ratioF values were analyzed as a function of baseline fluorescence measured under the Empty condition ($\text{Fmean}_{\text{Empty}}$). For each condition, $\log_2$ratioF values were plotted against $\text{Fmean}_{\text{Empty}}$ using scatter plots with linear regression lines for visualization. Linear regression slopes, Pearson correlation coefficients, and associated p-values were calculated using least squares regression. To further estimate the scaling relationship between baseline folding levels and $\log_2$ratioF values while accounting for measurement uncertainty in both variables, Deming regression was also performed assuming equal error variance ($\lambda = 1$). Deming regression slopes and intercepts were computed for each condition.

For visualization, y-axis ranges were standardized across panels to allow comparison of slope magnitudes while preventing extreme compression in conditions with narrow dynamic ranges.

**Consistency analysis between residual and $\log_2$ratioF**

To evaluate whether the residual-based metric captures the same folding signal as the conventional $\log_2$ratioF metric, we examined the relationship between centered residual values and $\log_2$ratioF for each chaperone condition. Centered residuals were computed relative to the Empty baseline regression and mean-centered within each condition as described above. For each variant, centered $\text{residual}_{\text{chap}}$ was plotted against $\log_2\text{ratioF}_{\text{chap}}$, and the relationship was evaluated using linear regression and Pearson correlation analysis.

**Depth artifact control analysis**

To assess whether sequencing depth inflates residual dispersion, we examined the relationship between pooled read depth (AD_sum_pooled) and the centered residuals calculated relative to the Empty baseline model as described above. Because the magnitude of deviation from the baseline was the quantity of interest, absolute centered residuals were plotted against AD_sum_pooled for each chaperone condition. Linear regression and Pearson correlation analyses were performed independently for each condition to determine whether residual dispersion systematically varied with sequencing depth.

**Baseline-dependent redistribution analysis**

To examine whether deviations from the baseline folding model depend on the initial folding state of each variant, we analyzed the relationship between baseline folding levels and Empty-referenced residuals. Centered residual values were calculated relative to the Empty baseline regression and mean-centered within each condition as described above. For each chaperone condition, the baseline folding level of each variant was defined as the mean fluorescence measured under the Empty condition ($\text{Fmean}_{\text{Empty}}$). Centered residual values for each variant were then plotted against $Fmean_{Empty}$. Linear regression and Pearson correlation coefficients were computed to quantify the relationship between baseline folding level and residual deviation.

To reduce replicate-level noise and ensure consistent variant representation across conditions, residual values were averaged at the residue level prior to analysis. All analyses were performed independently for each experimental round.

**Rank-dependent redistribution analysis**

Variants were ranked according to Fmean under the corresponding Empty condition and partitioned into equal-frequency baseline bins using quantile-based discretization (Python pandas qcut, with bin-edge adjustment for duplicate values). For the primary analysis, five equal-frequency bins (quintiles; Q1-Q5) were generated. Within each biological replicate and chaperone condition, variants were independently ranked according to Fmean, and upward redistribution was defined as a positive rank shift relative to the corresponding Empty baseline. For each Empty baseline quintile, the fraction of variants exhibiting upward redistribution was calculated at the replicate level and subsequently averaged across biological replicates.

The increase in upward redistribution from Q3 ($40^{th}$–$60^{th}$ percentile) to Q4 ($60^{th}$–$80^{th}$ percentile) was used to quantify mid-high percentile amplification. To assess robustness to percentile resolution, the analysis was repeated using ten equal-frequency bins (deciles).

As an additional sensitivity analysis,10 variants that consistently occupied the extreme Empty baseline bins (Q1 or Q5) and exhibited minimal rank redistribution across chaperon conditions were excluded and the redistribution analysis was repeated.

Statistical analyses were performed using replicate-level upward-redistribution fractions. For the quintile analysis, a planned linear contrast was used to compare the mean upward-redistribution fraction of the three protein-based chaperones (GroEL, DnaK, and Spy) with that of Seq576 at Q4, corresponding to the $60^{th}$–$80^{th}$ percentile baseline-foldability range. Effect estimates, 95% confidence intervals, t statistics, and p-values were obtained from the fitted linear model.

To evaluate redistribution patterns at higher percentile resolution, decile-based data were analyzed using a linear model including condition, quantile, and their interaction. The overall condition × quantile interaction was evaluated by ANOVA. Targeted planned contrasts were subsequently used to compare the combined protein-based-chaperone response with Seq576 within the Q7–Q8 range. P-values from the targeted comparisons were adjusted for multiple testing using the Holm method.

**ThermoMPNN stability prediction**

Predicted stability effects of alanine substitutions were estimated using ThermoMPNN (32). The crystal structure of the reporter protein (PDB ID: 4KGF) was used as structural input. For each residue, alanine substitution was evaluated and the predicted stability change (ΔΔG_pred) was calculated using the ThermoMPNN inference pipeline. Predicted mutations were filtered to include only single alanine substitutions corresponding to the experimental alanine-scanning library, and residue numbering was adjusted to match PDB positions. The resulting ΔΔG_pred values were compared with experimentally inferred chaperone-dependent folding effects ($log_2ratioF$).

**Codon usage analysis**

To assess whether codon usage contributes to residue-level chaperone-dependent folding effects, we performed two analyses. First, local codon usage frequency was calculated for each residue as the average codon usage frequency within a ±3 residue window and compared with $log_2ratioF$ for each chaperone condition using Spearman correlation. Next, native alanine positions originally encoded by GCC or GCT were examined to determine whether synonymous codon identity was associated with scaled $log_2ratioF$ values. Ten alanine positions were included and scaled $log_2ratioF$ values were compared between GCC- and GCT-encoded groups using a two-sided Mann-Whitney U test.

**Positional analysis of chaperone-dependent folding effects**

To examine whether chaperone-dependent folding effects varied along the TagRFP675 sequence, residue position was compared with $log_2ratioF$ for each condition using Spearman rank correlation. LOWESS smoothing (fraction = 0.18) was applied for visualization of broad positional trends. Analyses were performed independently for GroEL, DnaK, Spy, and Seq576.

**Data Availability**

Raw sequencing data generated in this study will be deposited in the NCBI Sequence Read Archive (SRA) and made publicly available upon publication. Processed sequencing and residue-level folding datasets are provided in Dataset S1. Structural visualization and coordinate datasets provided in Dataset S2 are publicly available through Zenodo (https://doi.org/10.5281/zenodo.20274818).

### SI Figures

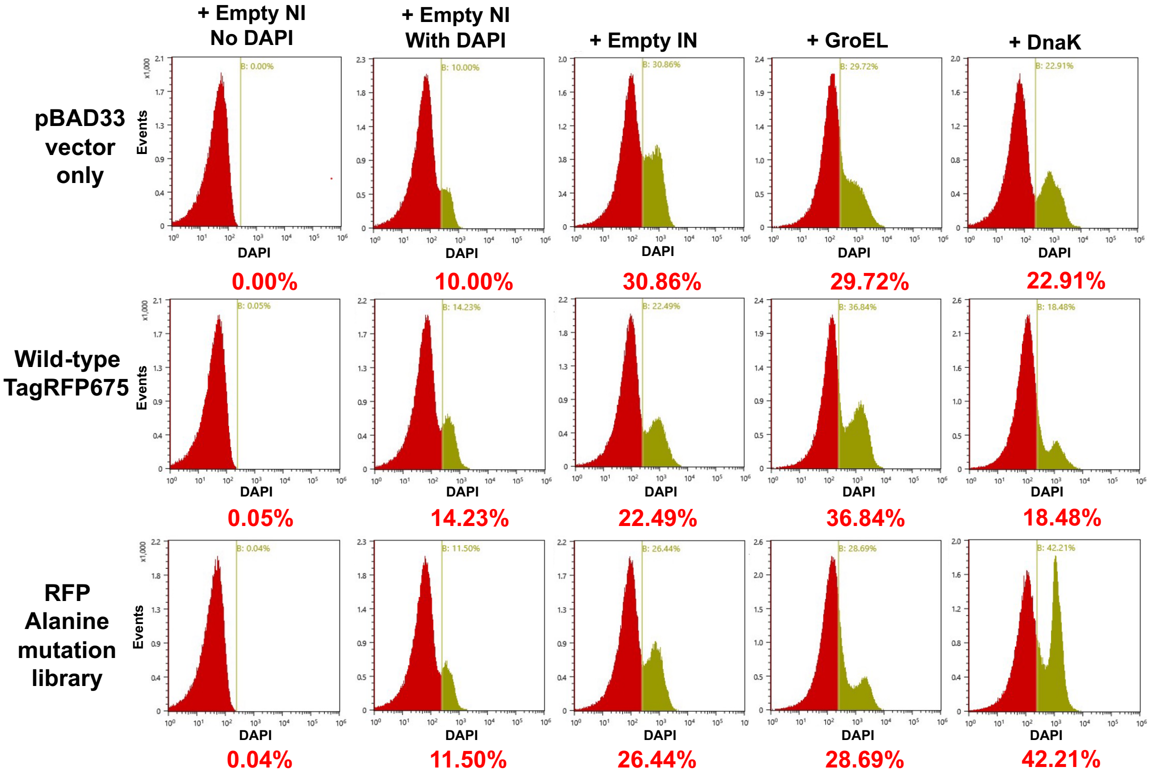


**Fig. S1. Cell sorting robustness and viability assessed by DAPI staining.**
Representative DAPI fluorescence histograms for cells subjected to FACS under multiple experimental conditions, including pBAD33 vector only controls, WT TagRFP675 expressed with chaperones, and RFP alanine mutation libraries expressed with chaperones. Olive-colored populations indicate cells with high DAPI fluorescence, which were excluded from all sorting gates. DAPI-high fractions were defined relative to the empty vector, non-induced control (top, left) and ranged from approximately 20–40% across all conditions. No systemic condition-specific increase in the DAPI-high populations was observed, indicating comparable cell viability during sorting regardless of chaperone expression or library complexity.

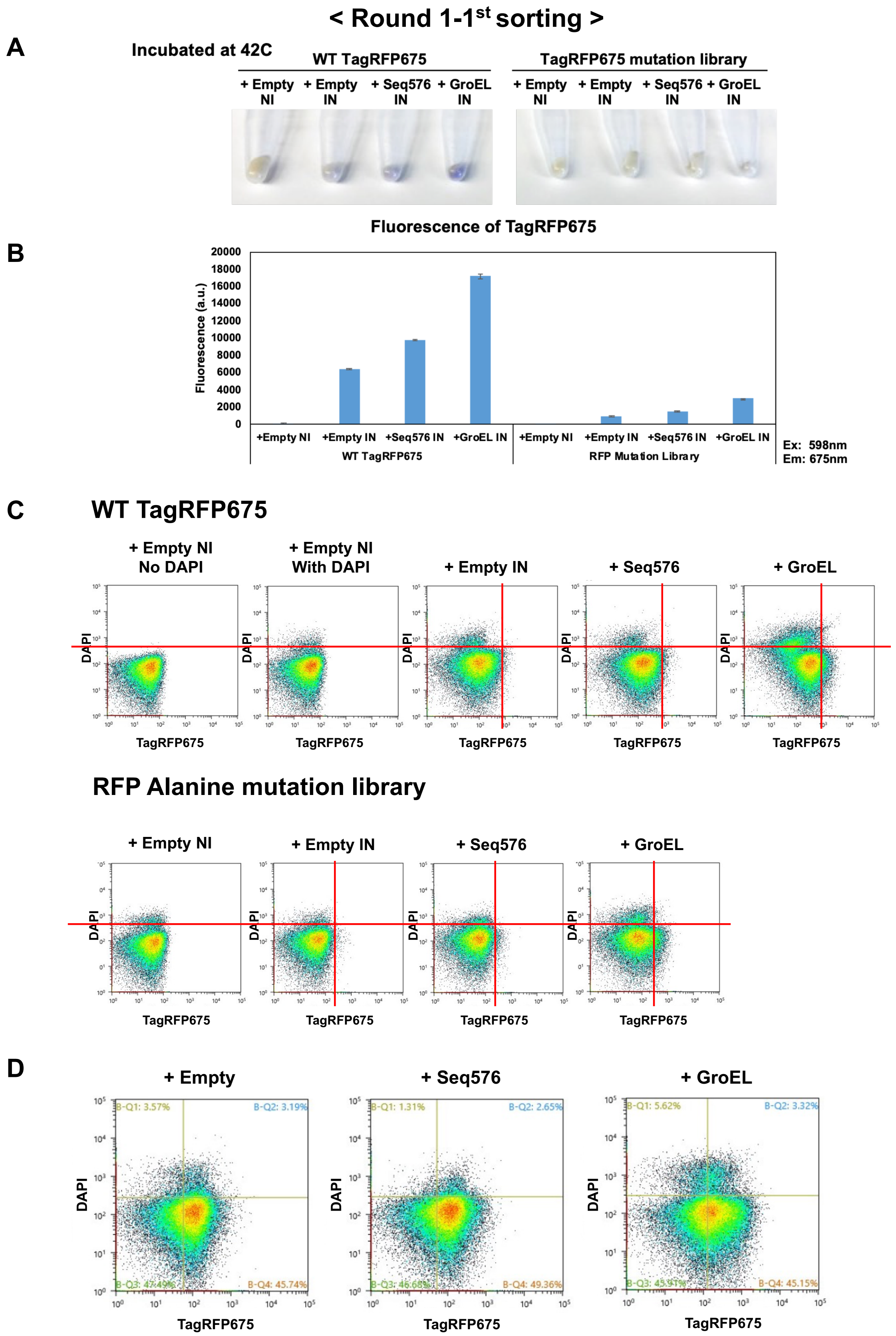


**(Fig. S2. Continued)**

### < Round 1-1st validation and Round 1-2nd sorting >

E

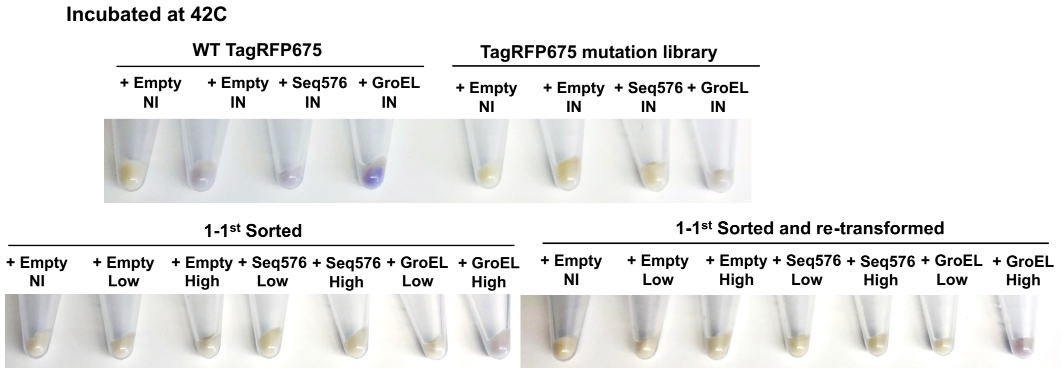


F

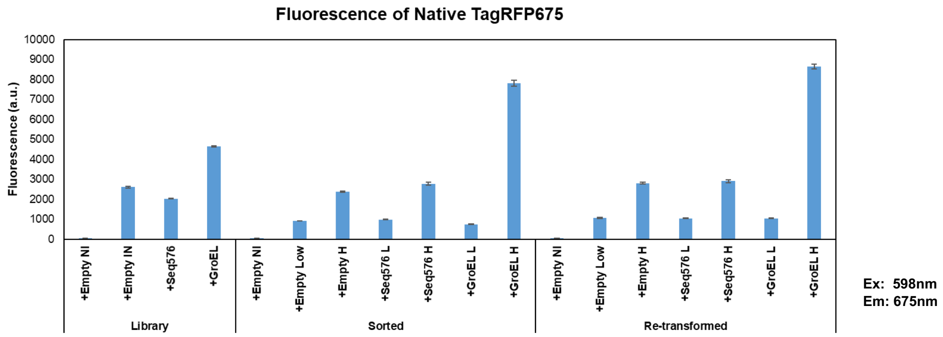


G

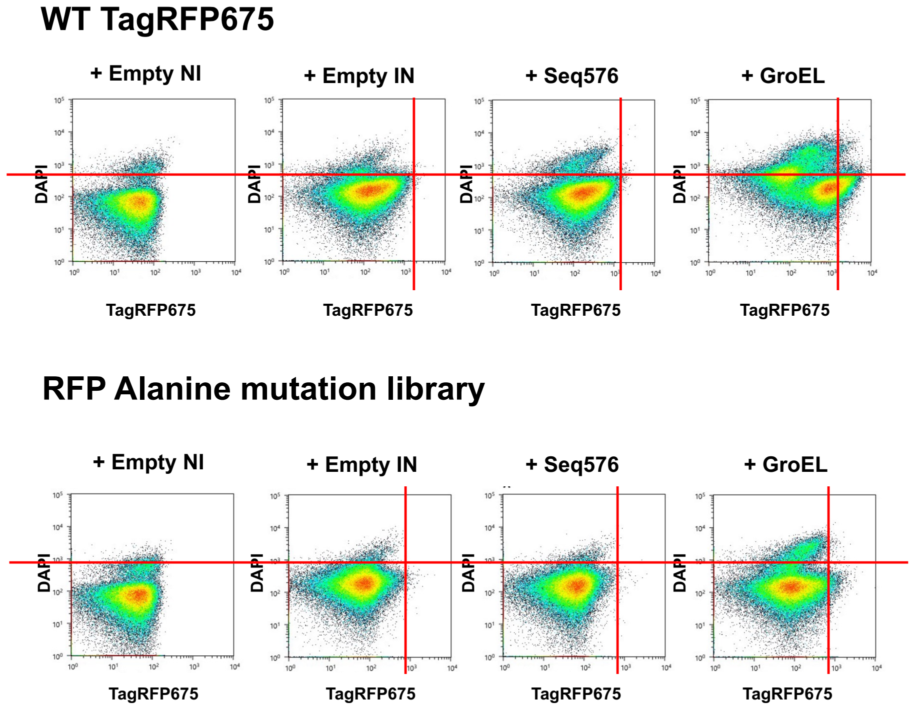


**(Fig. S2. Continued)**

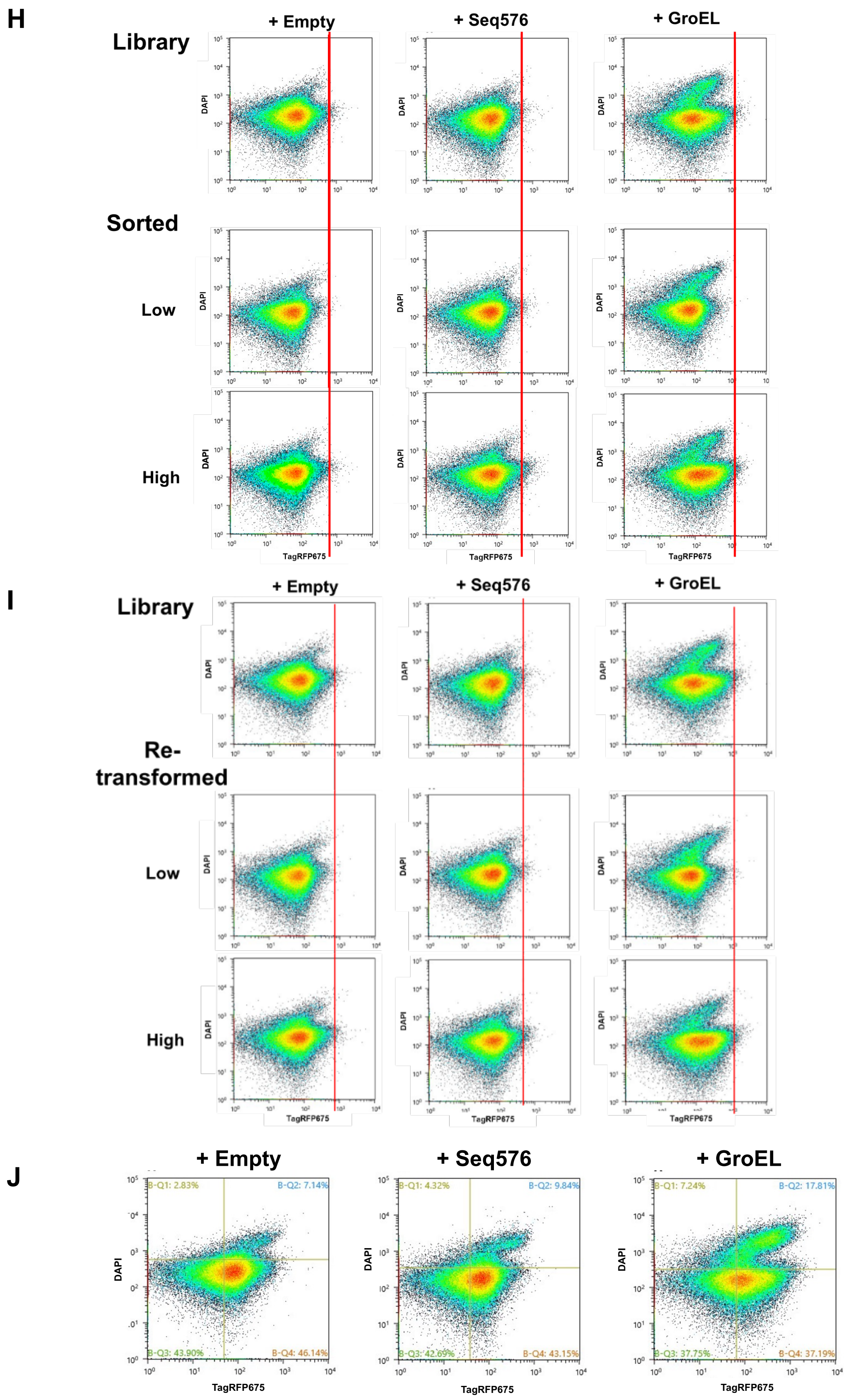


**(Fig. S2. Continued)**

### < Round 1-2nd validation and Round 1-3rd sorting >

K

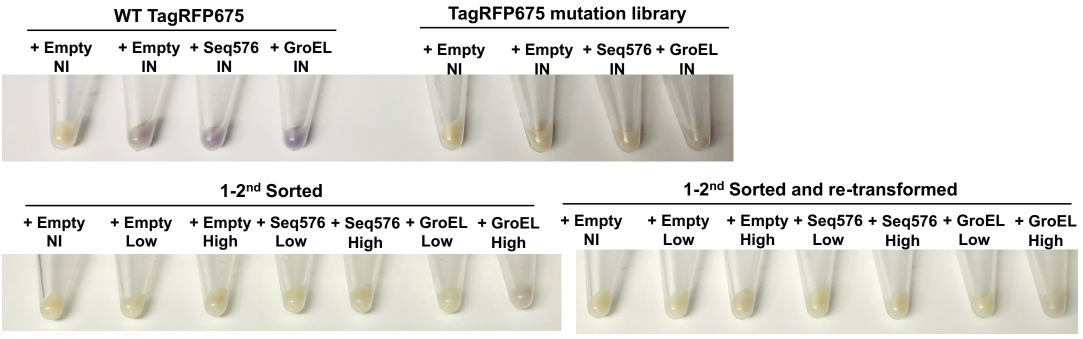


L

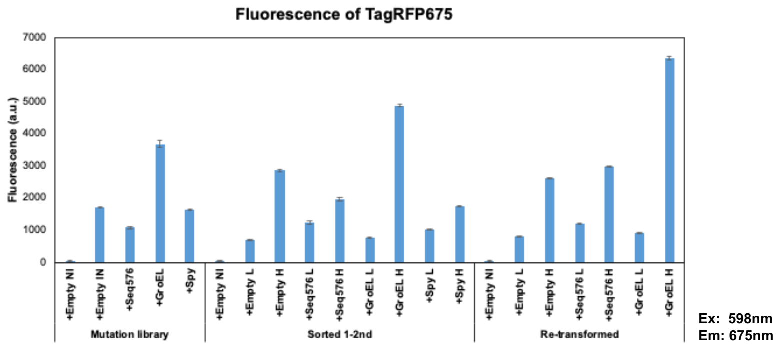


M

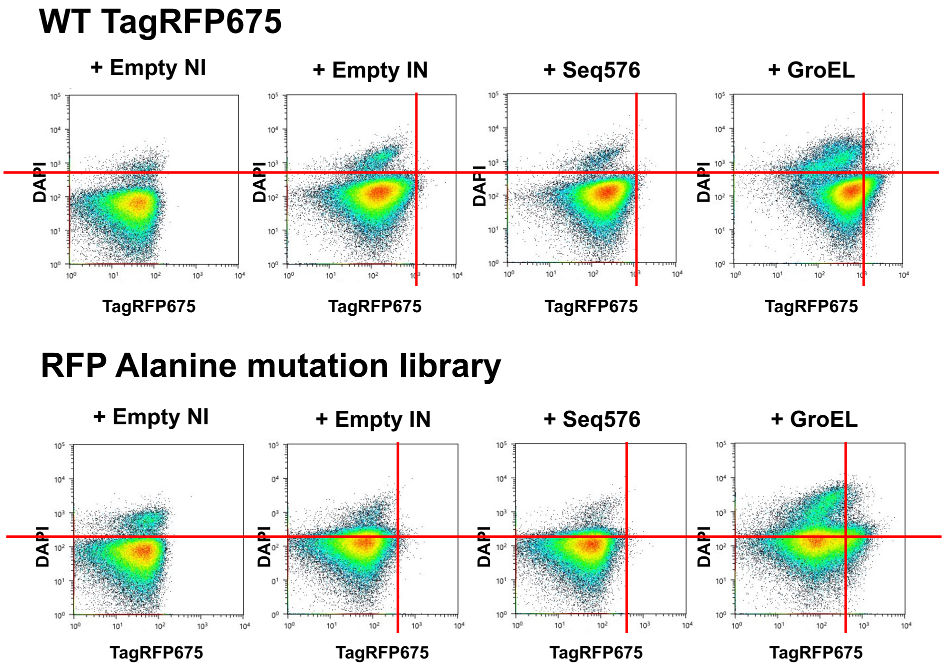


**(Fig. S2. Continued)**

N

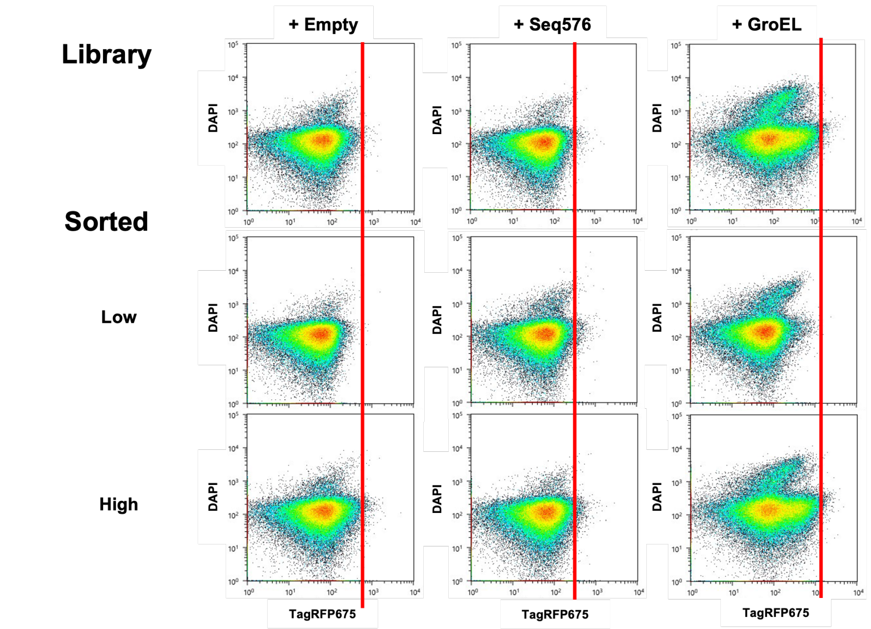


O

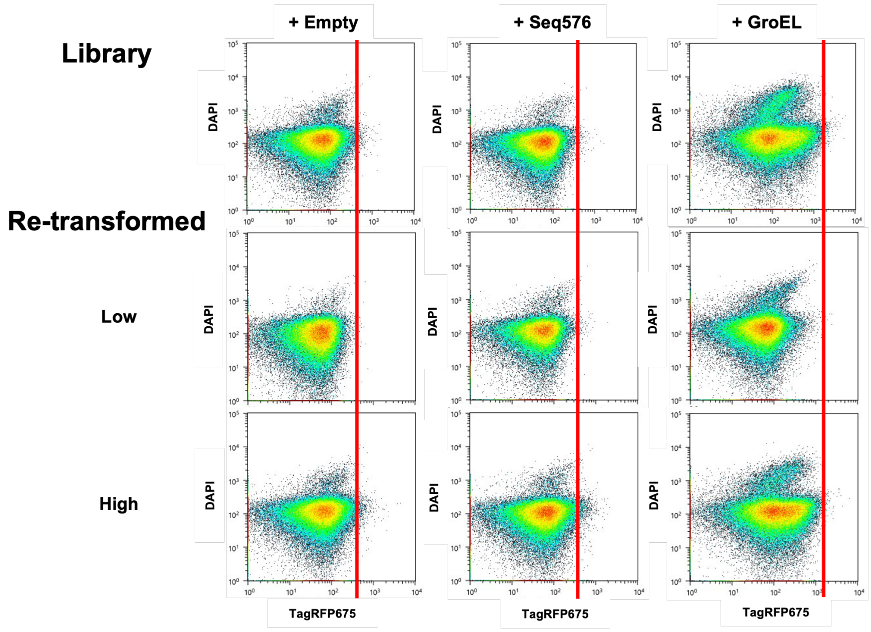


P

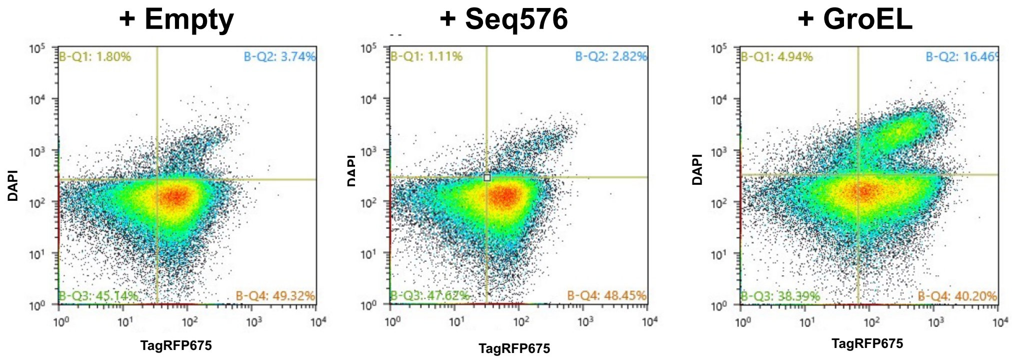


(Fig. S2. Continued)

### < Round 1-3$^{rd}$ validation >

**Q**

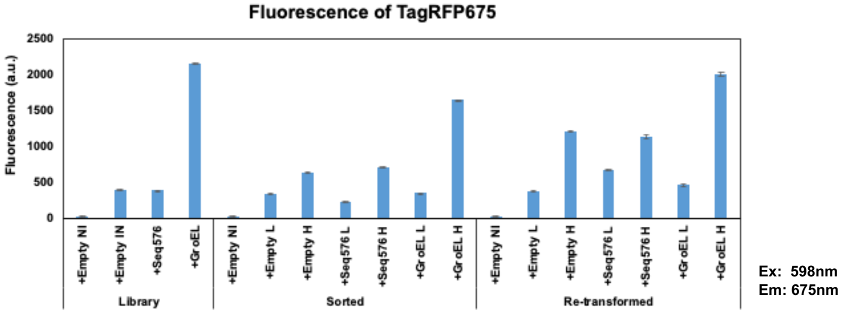


**R**

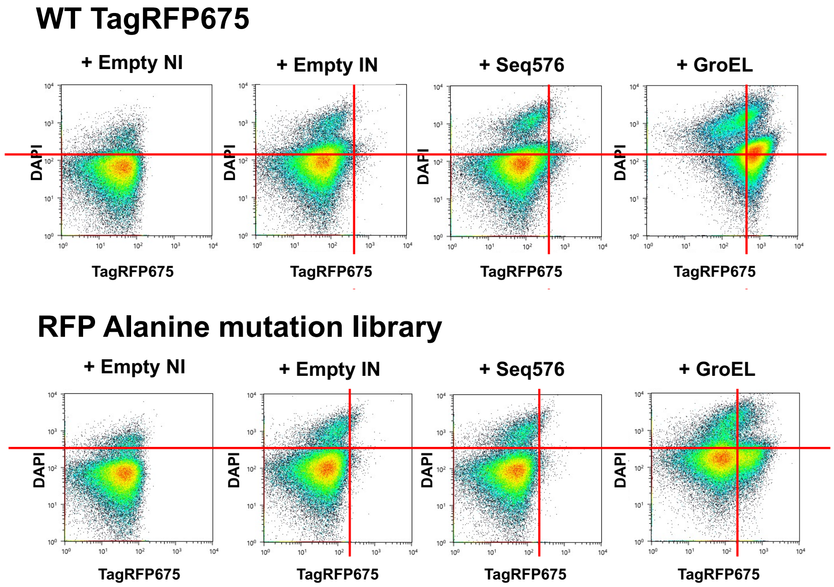


**S**

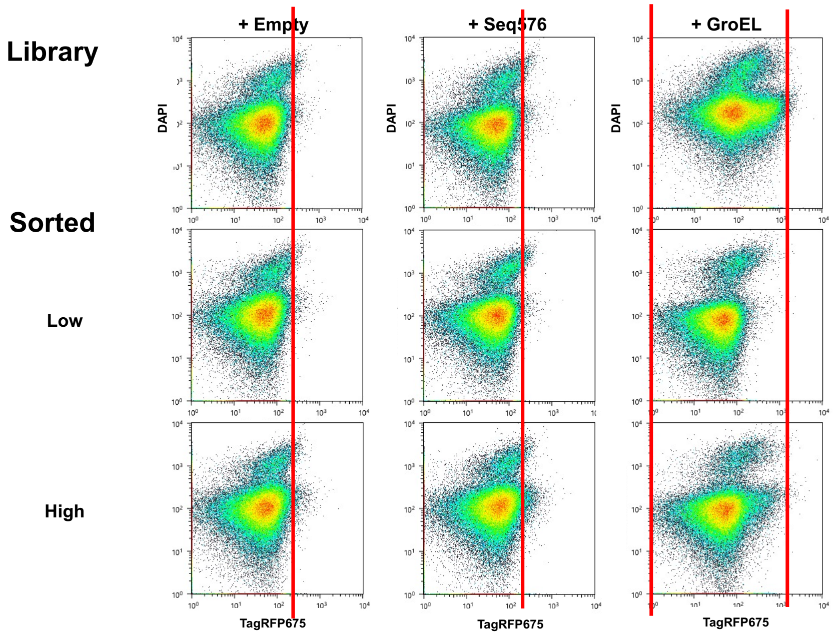


**(Fig. S2. Continued)**

T

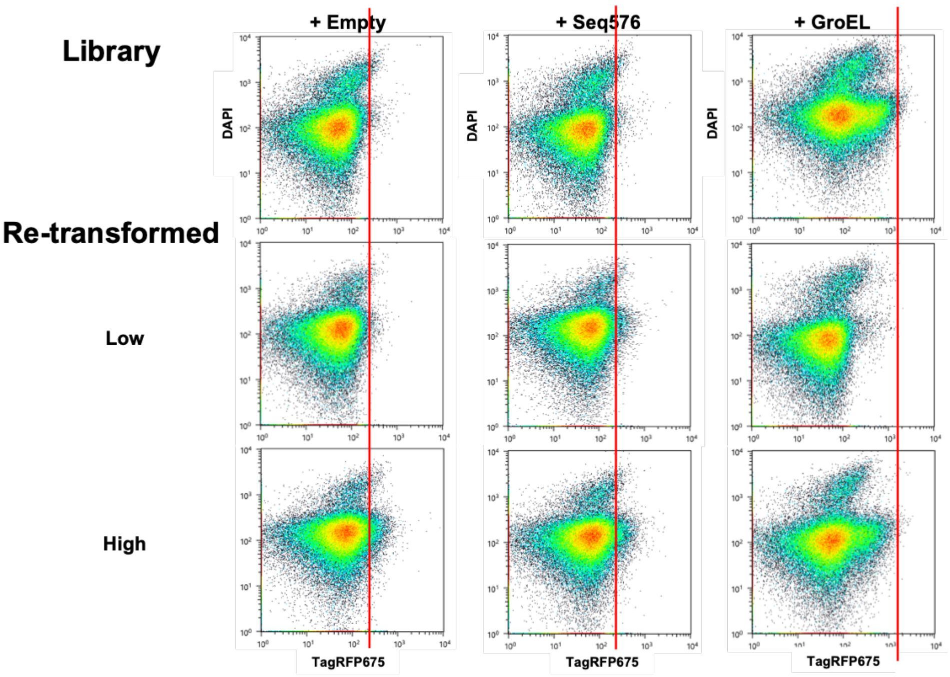


**Fig. S2. Iterative FACS sorting and validation of folding-dependent populations during Round 1.** **(A–D) Round 1-1st sorting.** (**A**) Representative cell pellet color comparison for WT TagRFP675 and alanine mutation libraries expressed with chaperones (Empty, GroEL, Seq576). (**B**) Bulk fluorescence measurements of WT and mutation library populations prior to sorting. (**C**) FACS fluorescence profiles of WT (top) and mutation libraries (bottom). (**D**) Initial FACS-based separation of mutation libraries (1-1st sorting) into low- and high-folding populations using an approximately 50:50 gating strategy.

**(E–J) Validation of Round 1-1st sorting and Round 1-2nd sorting.** (**E**) Cell pellet color comparison among WT, mutation library, sorted populations, and re-transformed (sorted populations after plasmid extraction and re-transformation into fresh cells). (**F**) Bulk fluorescence measurements demonstrating substantially higher fluorescence in high-folding populations compared to low-folding populations for both sorted and re-transformed libraries. Low-folding populations consistently exhibited fluorescence levels lower than those of the mutation libraries under the Empty condition. (**G**) FACS fluorescence profiles of WT (top) and mutation libraries (bottom). (**H**) Comparison of FACS profiles between the original mutation library and sorted low- and high-folding populations. (**I**) Comparison of FACS profiles between the original mutation library and re-transformed low- and high-folding populations. In both cases, high-folding populations show pronounced rightward shifts toward higher fluorescence intensities. (**J**) Second FACS sorting (Round 1-2nd sorting) based on validated fluorescence separation.

**(K–P) Validation of Round 1-2nd sorting and Round 1-3rd sorting.** (**K**) Cell pellet color comparison for WT, mutation library, sorted populations, and re-transformed populations. (**L**) Bulk fluorescence measurements confirming sustained enrichment of high-folding populations relative to low-folding populations. (**M**) FACS fluorescence profiles of WT (top) and mutation libraries (bottom). (**N**) FACS comparison between the original mutation library and sorted low- and high-folding populations. (**O**) FACS comparison between the original mutation library and re-transformed low- and high-folding populations, showing consistent population shifts toward higher fluorescence in high-folding samples. (**P**) Third FACS sorting (Round 1-3rd sorting).

**(Q–T) Final validation following Round 1-3rd sorting.** (**Q**) Bulk fluorescence measurements of low- and high-folding populations. (**R**) FACS fluorescence profiles of WT (top) and mutation libraries (bottom). (**S**) Comparison of FACS profiles between the original mutation library and sorted populations. (**T**) Comparison of FACS profiles between the original mutation library and re-transformed populations.

### < Round 2-1st sorting >

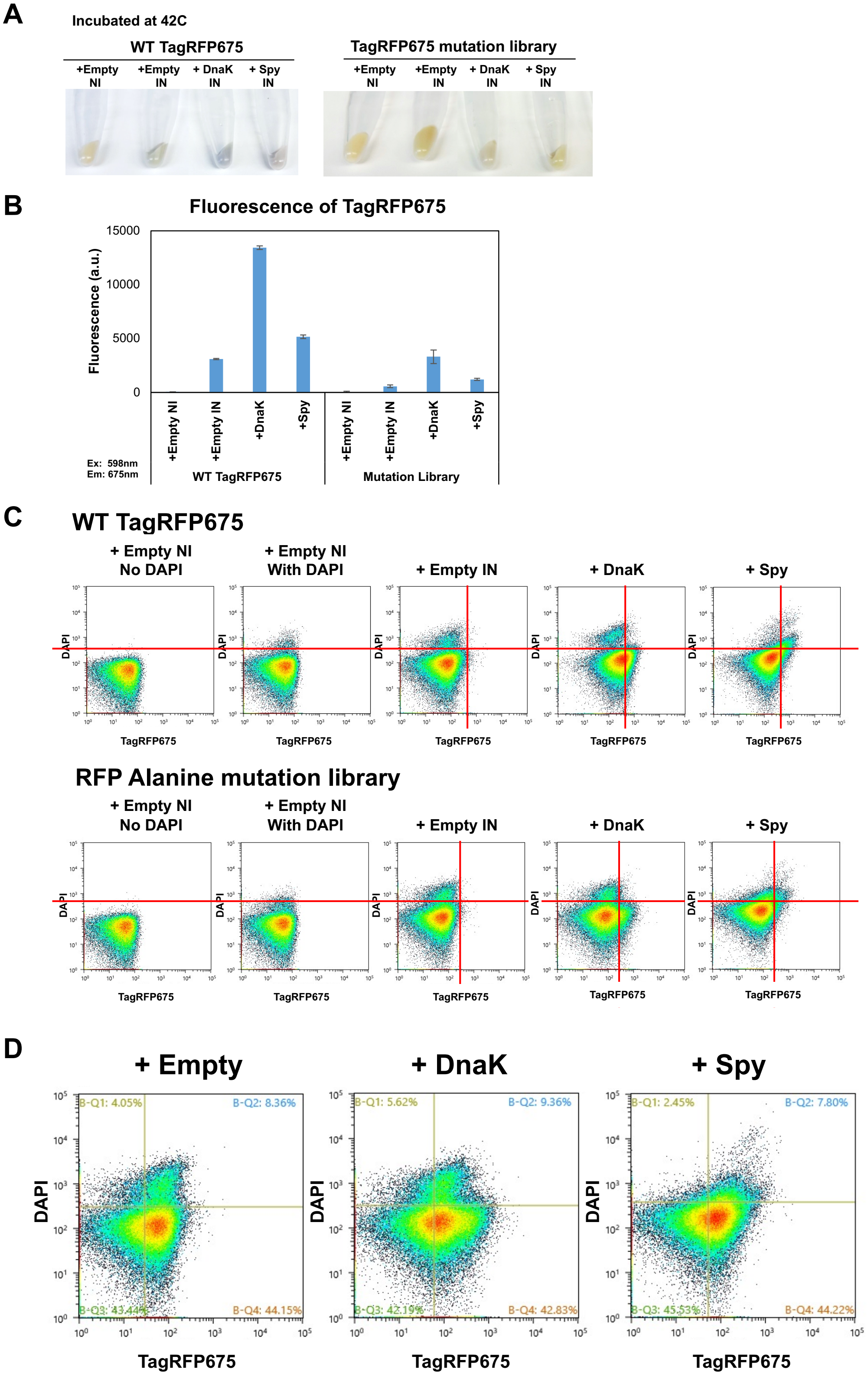


**(Fig. S3. Continued)**

< Round 2-1st validation and Round 2-2nd sorting >

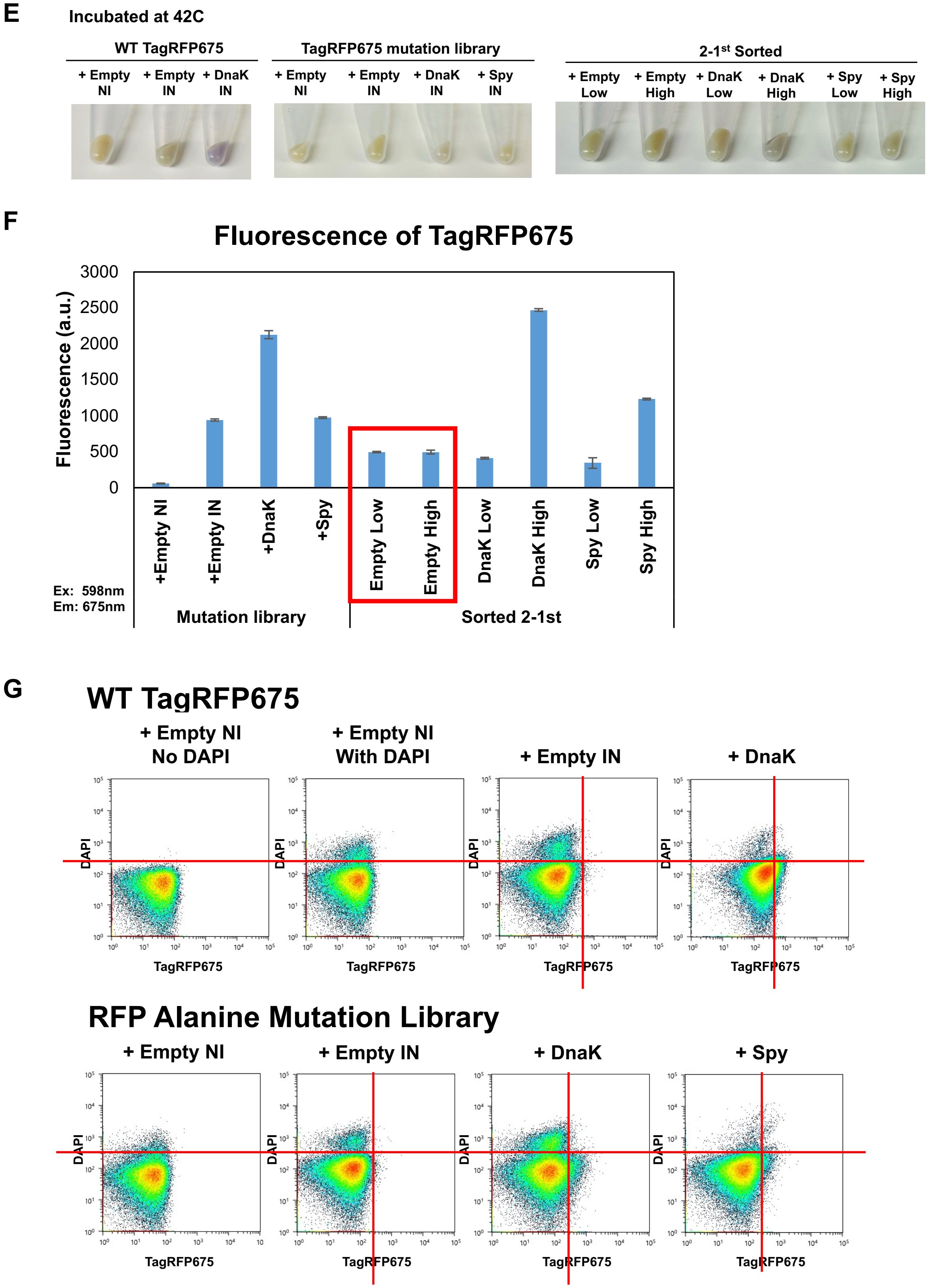


(Fig. S3. Continued)

H

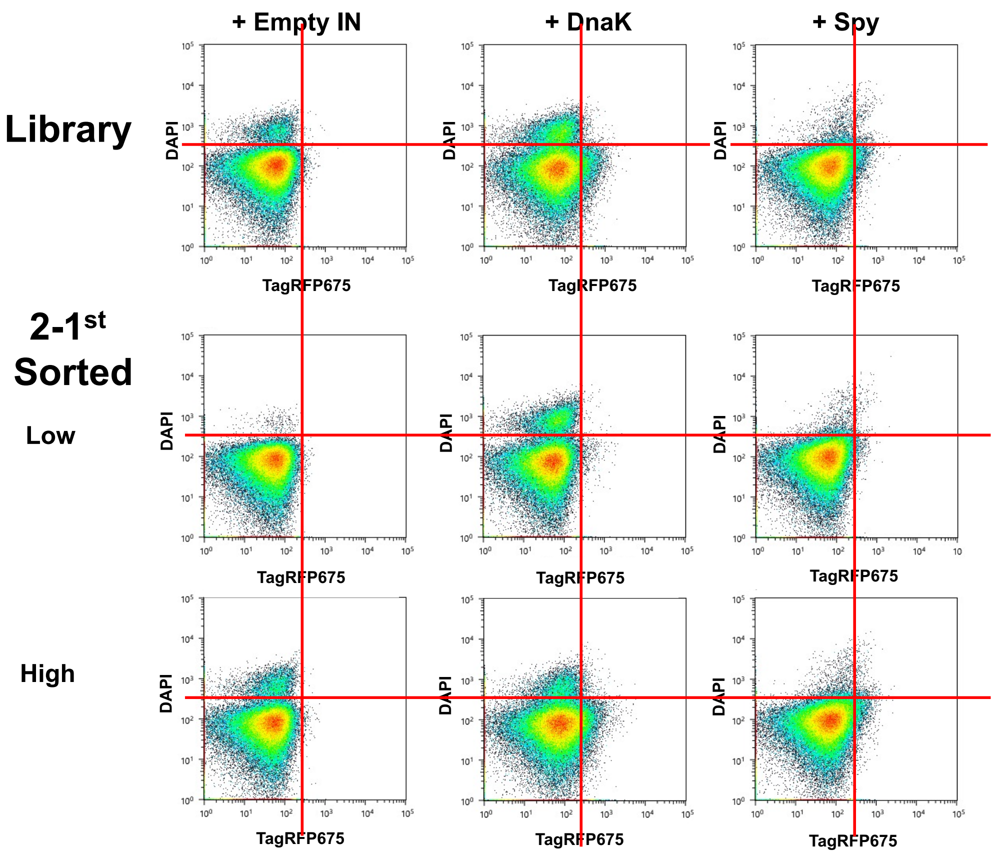


I

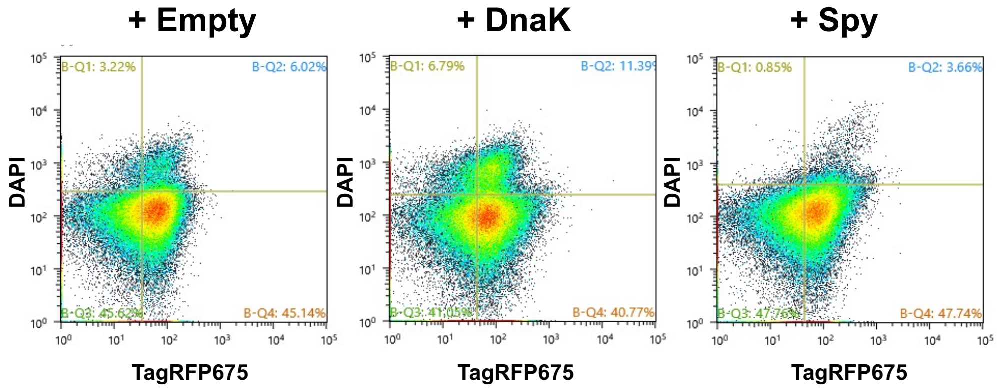


**(Fig. S3. Continued)**

### < Round 2-2nd validation and Round 2-3rd sorting >

J

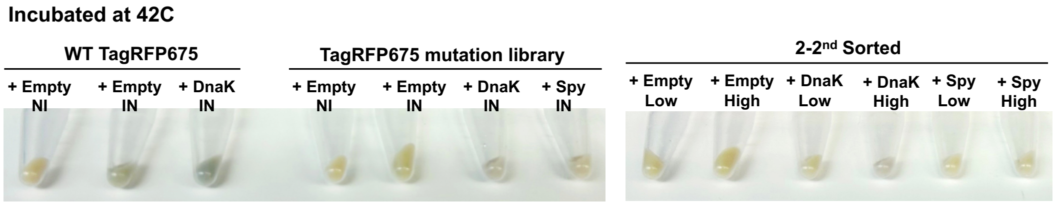


K

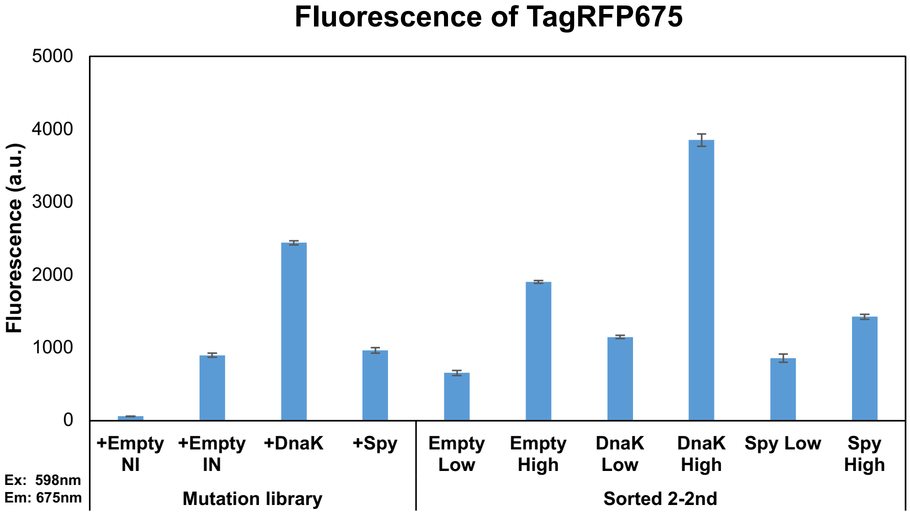


L

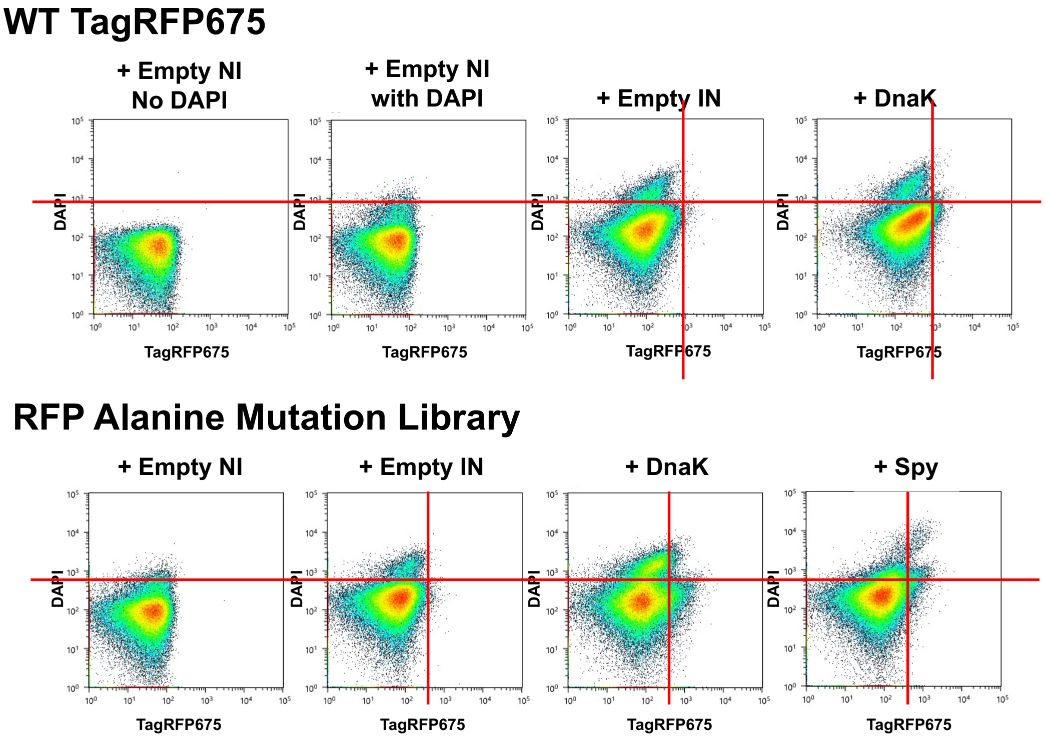


(Fig. S3. Continued)

M

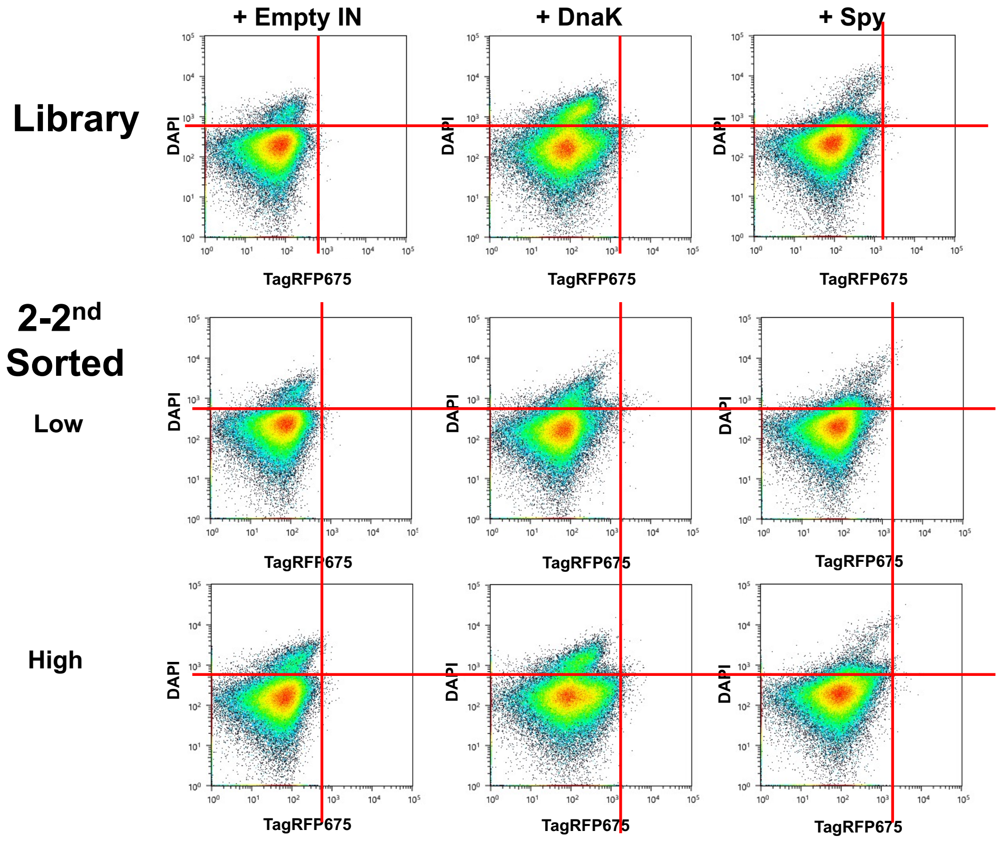


N

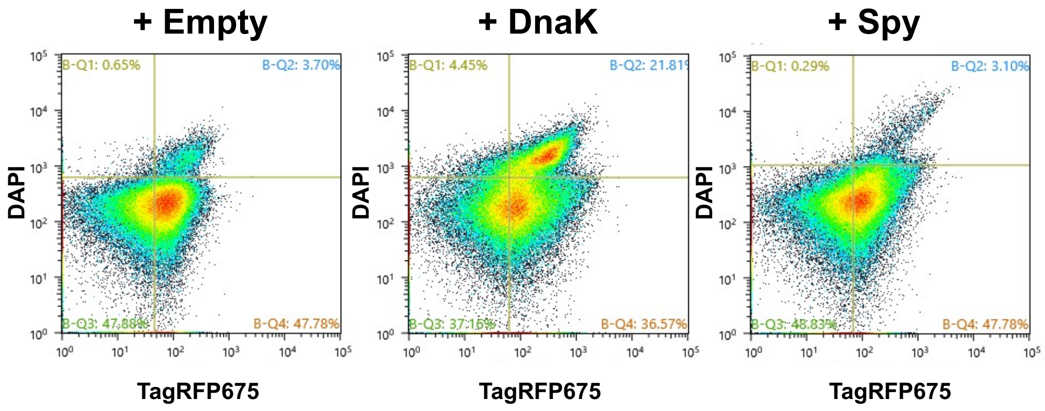


(Fig. S3. Continued)

### < Round 2-3rd validation and Round 2-4th and 2-5th sorting >

**O**

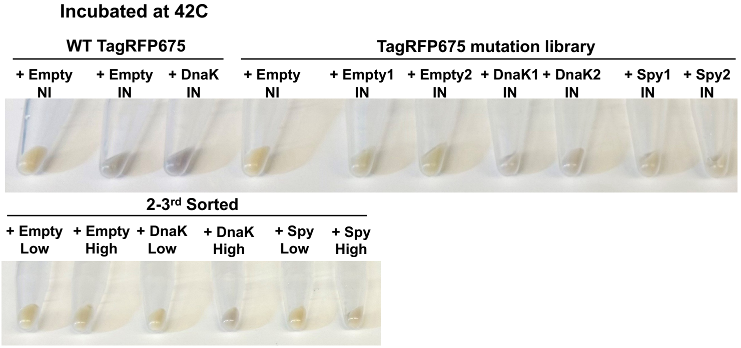


**P**

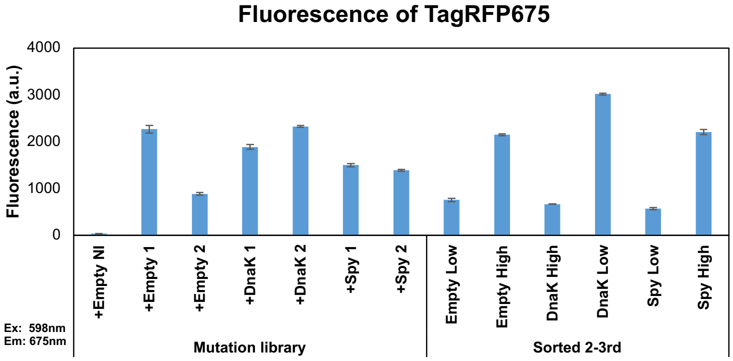


**Q**

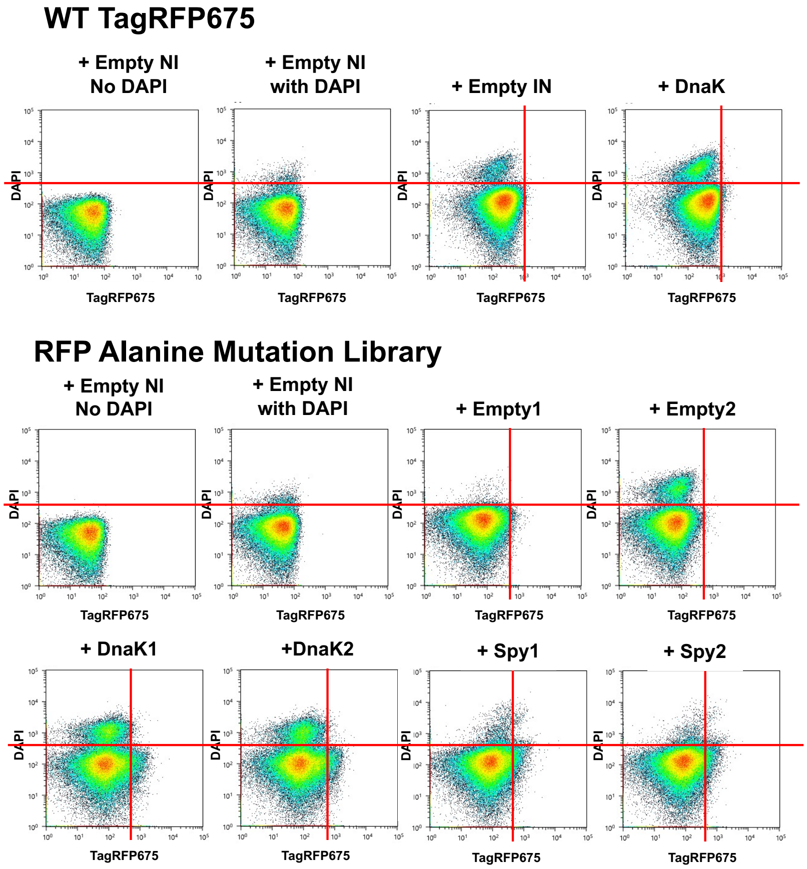


**(Fig. S3. Continued)**

R

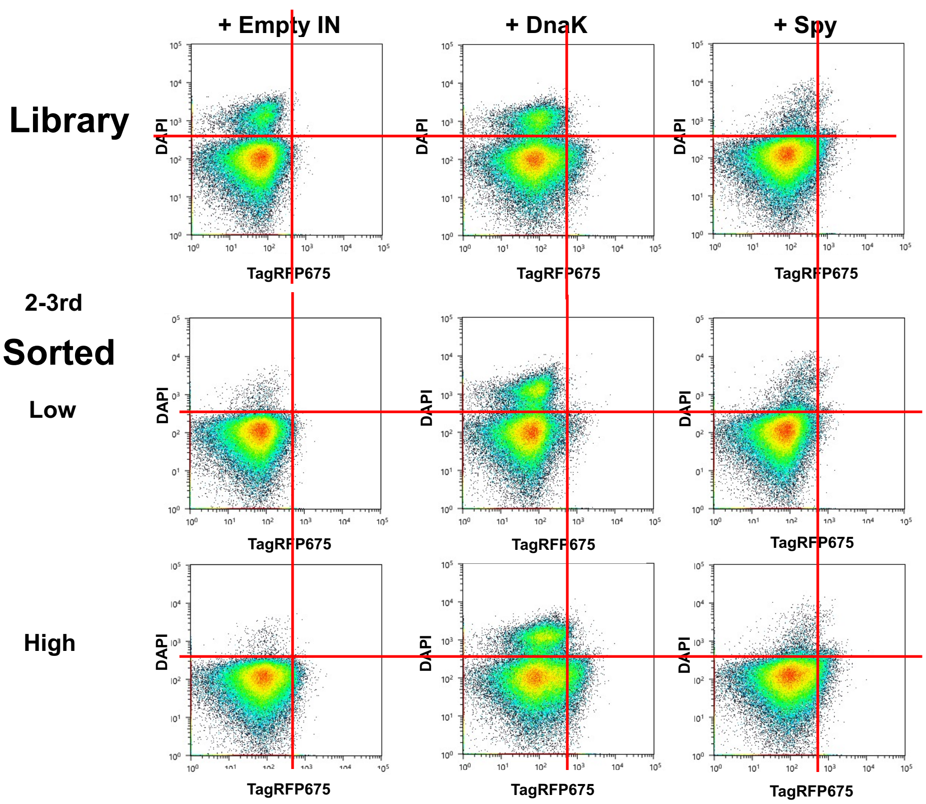


S

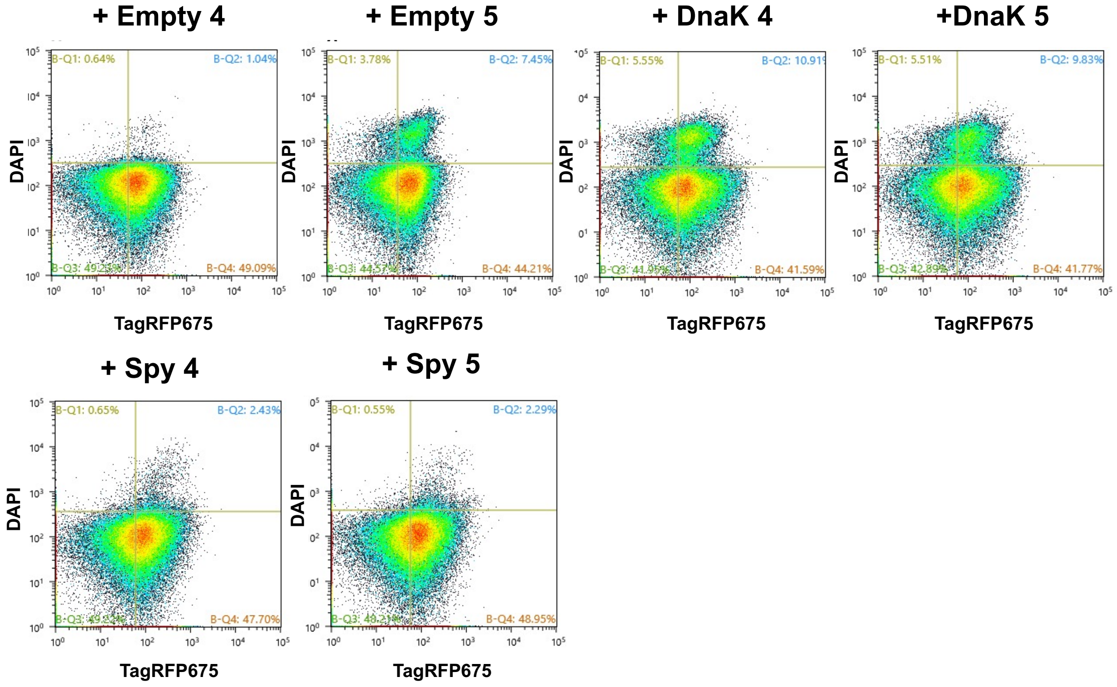


**(Fig. S3. Continued)**

### < Round 2-4$^{th}$ and 2-5$^{th}$ validation >

**T**

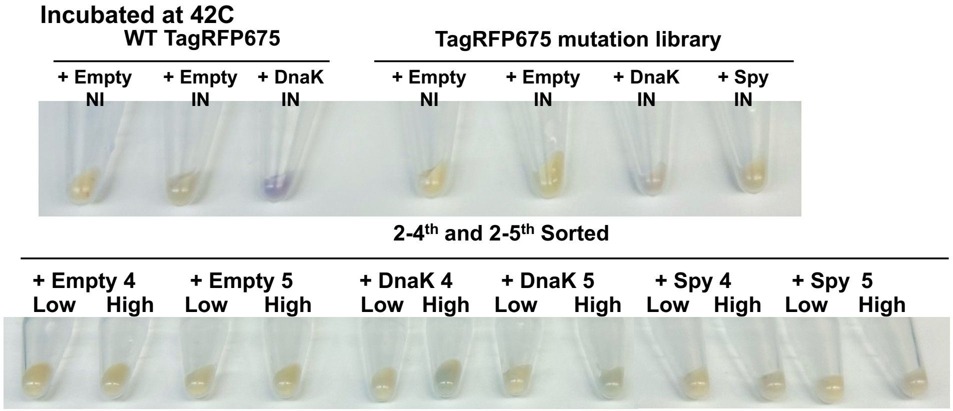


**U**

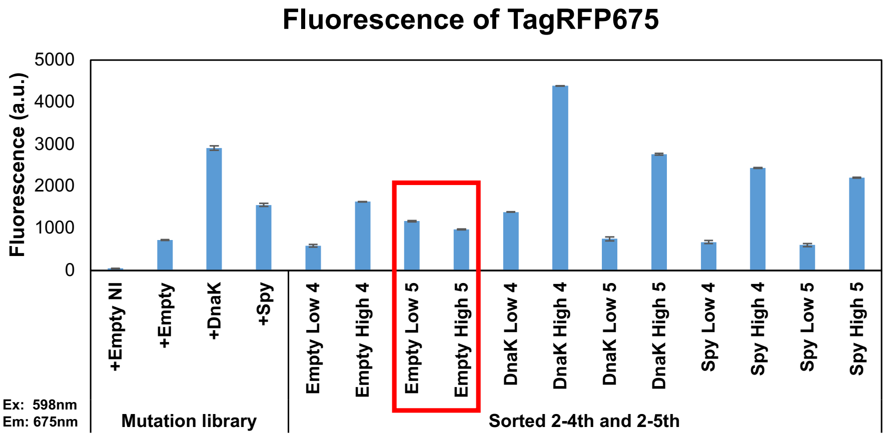


**V**

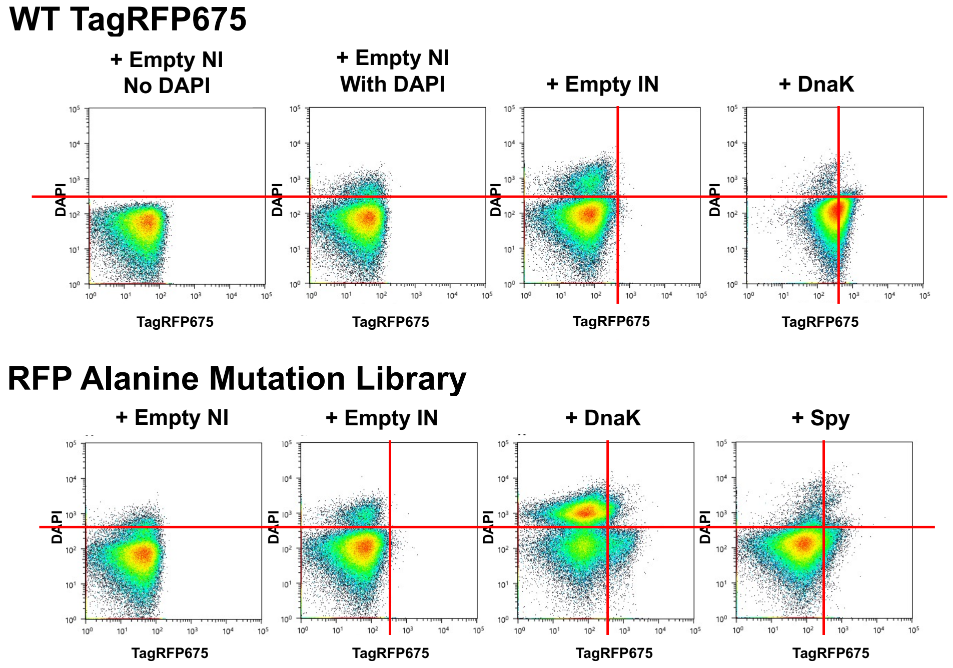


**(Fig. S3. Continued)**

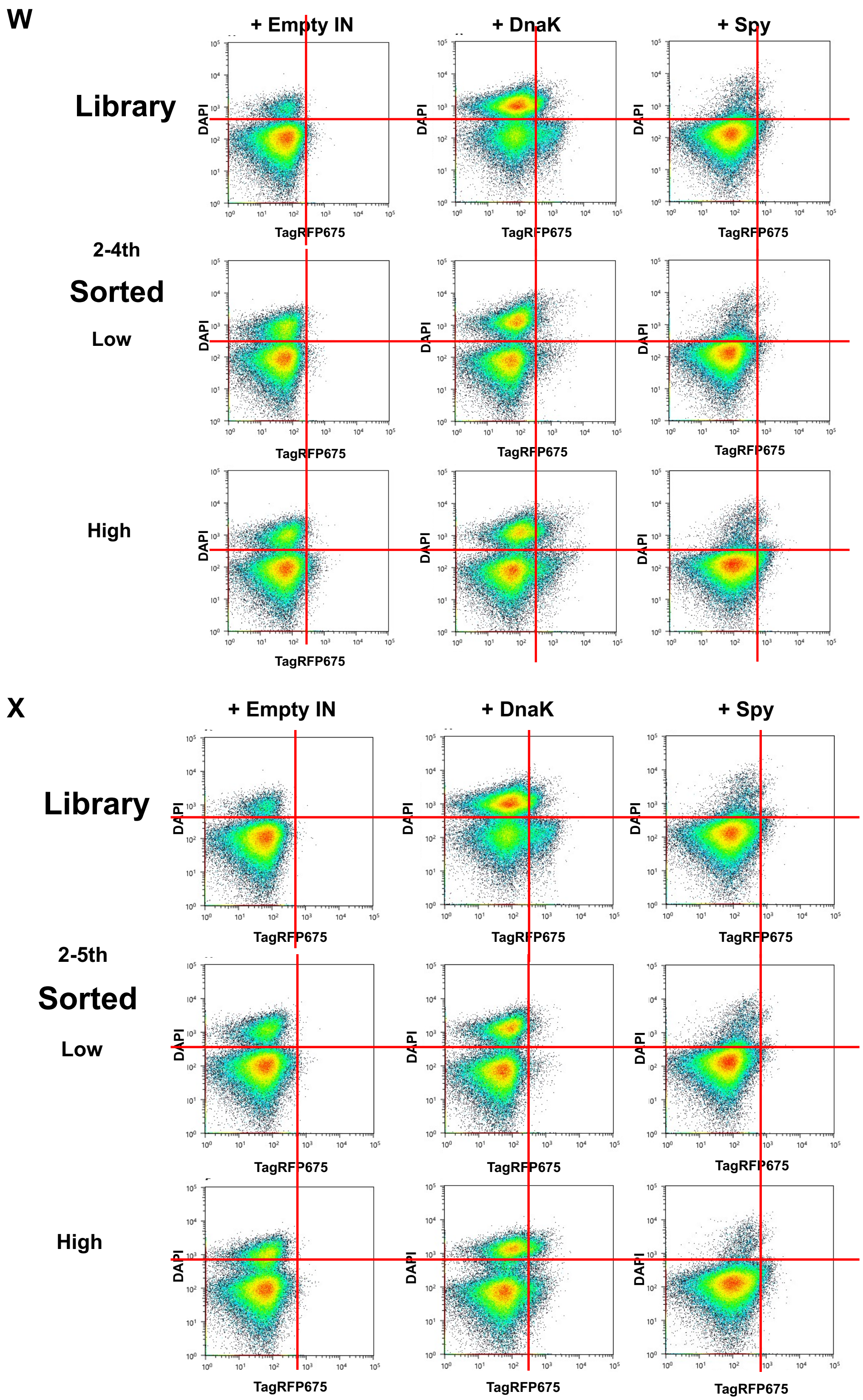


**(Fig. S3. Continued)**

### < Round 2 validation of re-transformed populations 1 >

**Y**

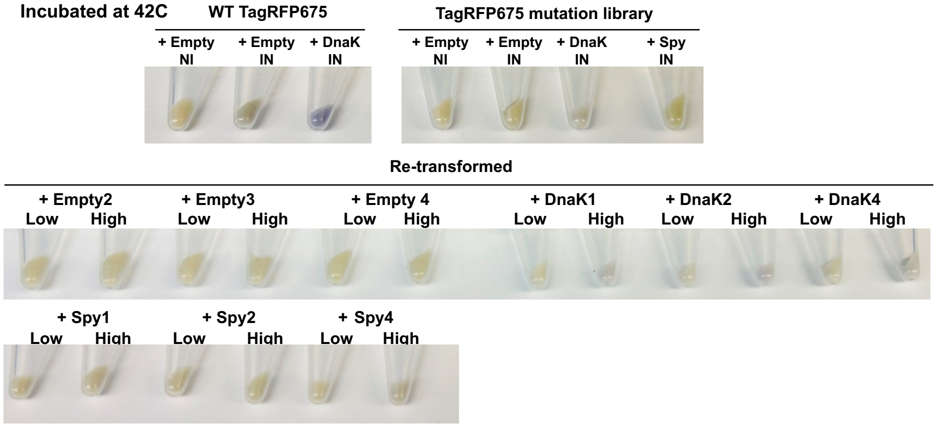


**Z**

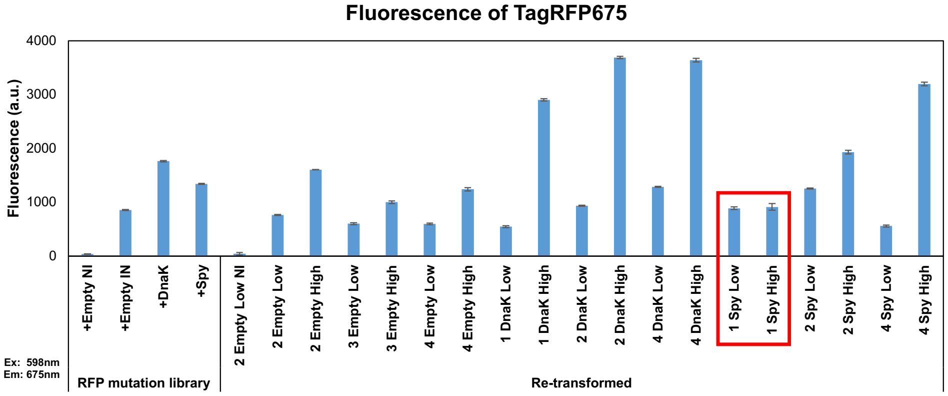


**AA**

#### WT TagRFP675

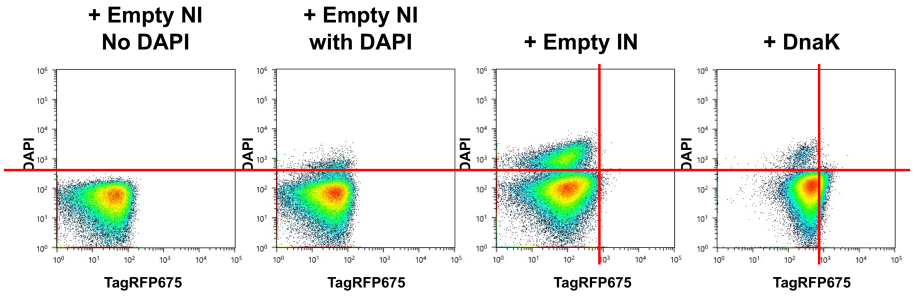


#### Mutation Library

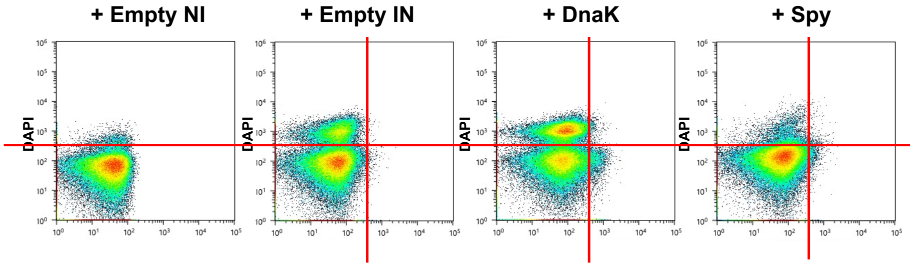


**(Fig. S3. Continued)**

AB

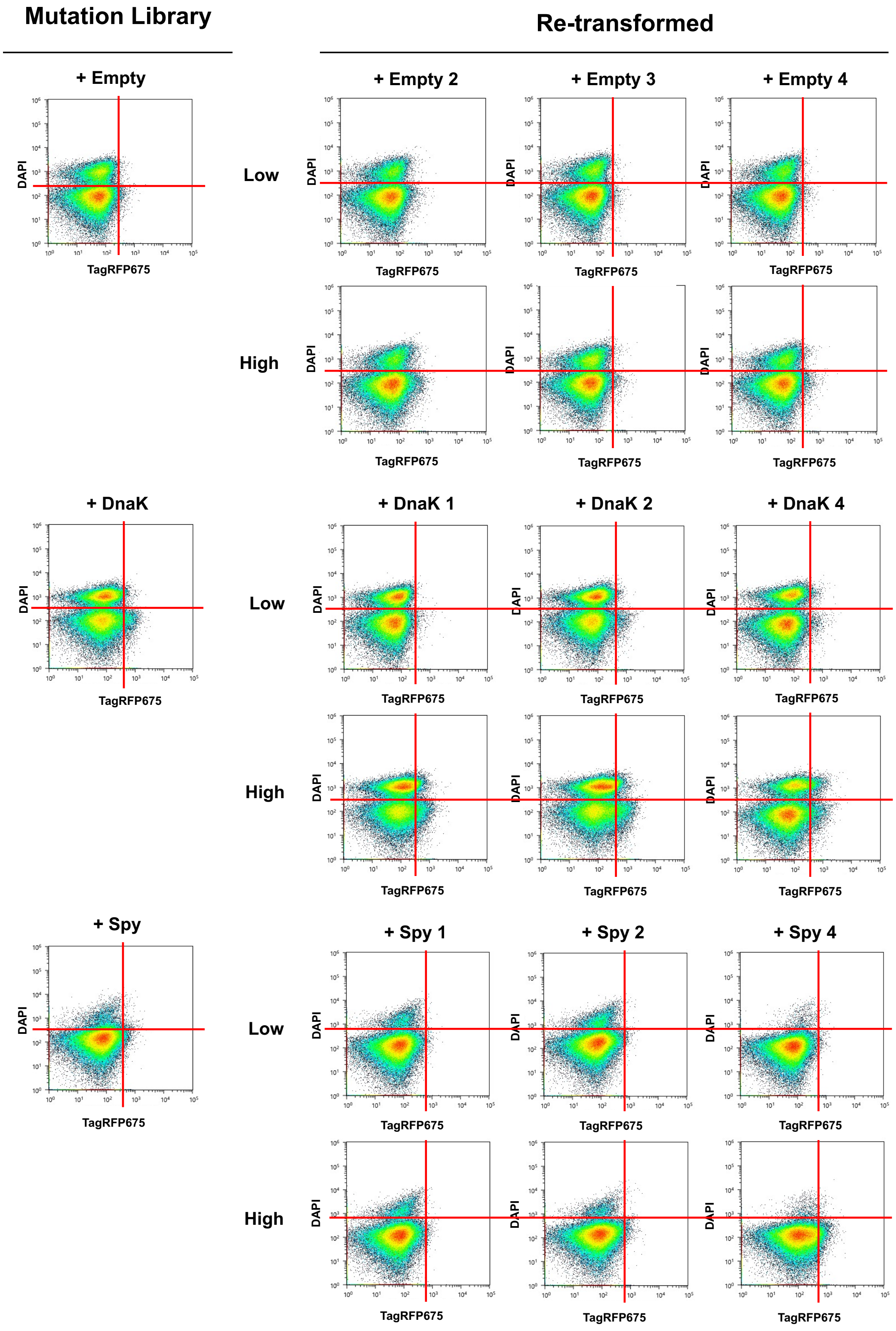


**(Fig. S3. Continued)**

< Round 2 validation of re-transformed populations 2 >

AC

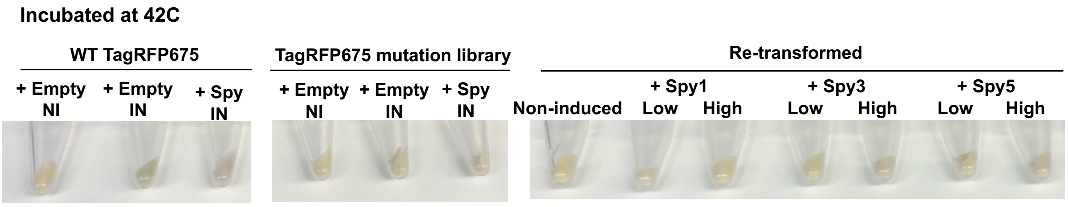


AD

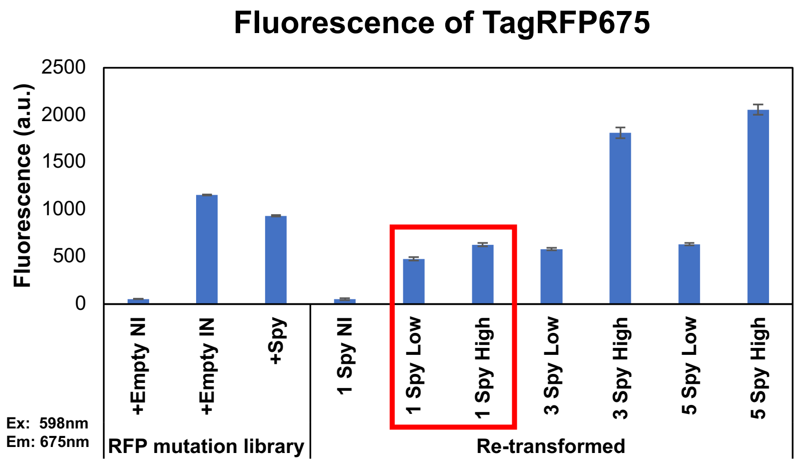


AE

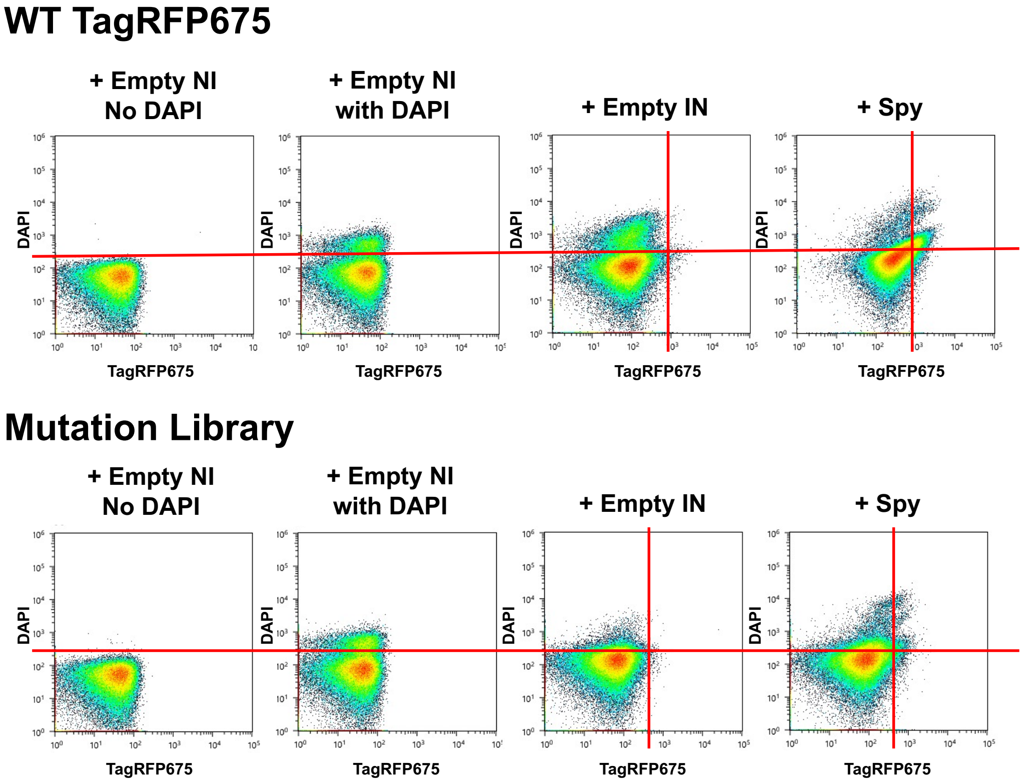


(Fig. S3. Continued)

**AF**

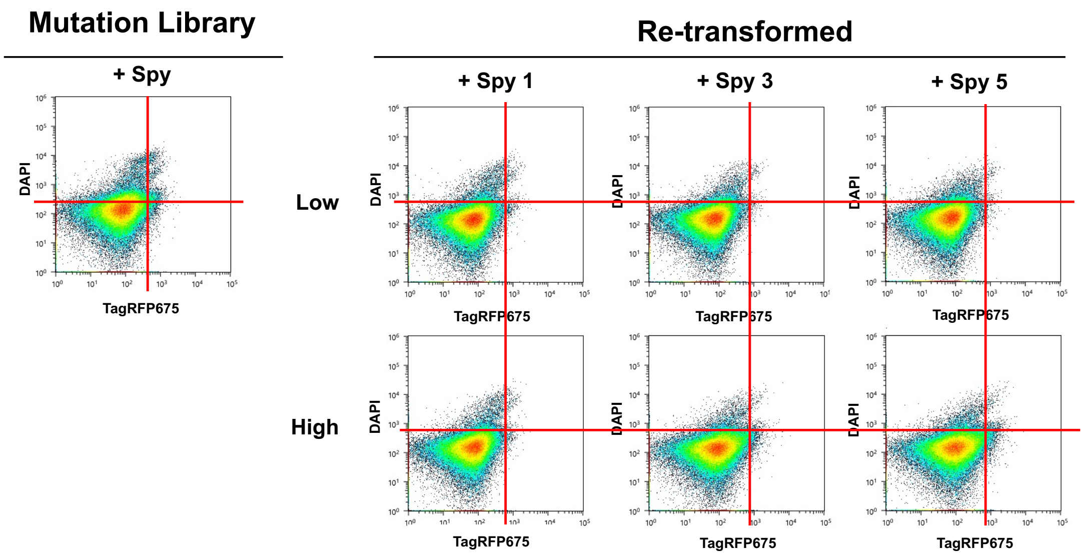


**Fig. S3. Iterative FACS sorting, validation, and quality-controlled replicate selection during Round 2.** **(A–D) Round 2-1$^{st}$ sorting.** (**A**) Cell pellet color comparison of WT and alanine mutation libraries prior to sorting. (**B**) Bulk fluorescence measurements of WT and mutation libraries. (**C**) FACS fluorescence profiles of WT (top) and mutation libraries (bottom). (**D**) First FACS-based separation of mutation libraries into low- and high-folding populations (Round 2-1$^{st}$ sorting).

**(E–I) Validation of Round 2-1$^{st}$ sorting and Round 2-2$^{nd}$ sorting.** (**E**) Cell pellet color comparison following the first sorting. (F) Bulk fluorescence measurements of sorted populations. (**G**) FACS fluorescence profiles of WT (top) and mutation libraries (bottom). (**H**) Comparison of FACS profiles between the original mutation library and sorted low- and high-folding populations. (**I**) Second FACS sorting (Round 2-2$^{nd}$ sorting).

**(J–N) Validation of Round 2-2$^{nd}$ sorting and Round 2-3$^{rd}$ sorting.** (**J**) Cell pellet color comparison following the second sorting. (**K**) Bulk fluorescence measurements of sorted populations. (**L**) FACS fluorescence profiles of WT (top) and mutation libraries (bottom). (**M**) Comparison of FACS profiles between the original mutation library and sorted low- and high-folding populations. (**N**) Third FACS sorting (Round 2-3$^{rd}$ sorting).

**(O–S) Validation of Round 2-3$^{rd}$ sorting and Round 2-4$^{th}$/2-5$^{th}$ sorting.** (**O**) Cell pellet color comparison following the third sorting. (**P**) Bulk fluorescence measurements of sorted populations. (**Q**) FACS fluorescence profiles of WT (top) and mutation libraries (bottom). (**R**) Comparison of FACS profiles between the original mutation library and sorted low- and high-folding populations. (**S**) Fourth and fifth FACS sorting (Round 2-4$^{th}$ and 2-5$^{th}$ sorting).

**(T–X) Validation of Round 2-4$^{th}$ and Round 2-5$^{th}$ sorting.** (**T**) Cell pellet color comparison following late-stage sorting iterations. (**U**) Bulk fluorescence measurements of sorted populations. (**V**) FACS fluorescence profiles of WT (top) and mutation libraries (bottom). (**W**) Comparison of FACS profiles between the original mutation library and sorted populations from the fourth iteration. (**X**) Comparison of FACS profiles between the original mutation library and sorted populations from the fifth iteration.

**(Y–AB) Validation of re-transformed populations (batch 1).** (**Y**) Cell pellet color comparison following re-transformation. (**Z**) Bulk fluorescence measurements of re-transformed populations. (**AA**) FACS fluorescence profiles of WT (top) and mutation libraries (bottom). (**AB**) Comparison of FACS profiles between the original mutation library and re-transformed low- and high-folding populations.

**(AC–AF) Validation of re-transformed populations (batch 2).** (**AC**) Cell pellet color comparison following re-transformation. (**AD**) Bulk fluorescence measurements of re-transformed populations. (**AE**) FACS fluorescence profiles of WT (top) and mutation libraries (bottom). (**AF**) Comparison of FACS profiles between the original mutation library and re-transformed low- and high-folding populations.

Bulk fluorescence measurements corresponding to validation-failed samples are indicated by red squares. Only samples passing all validation criteria were advanced to next-generation sequencing; final sample selection is summarized in Table S2.

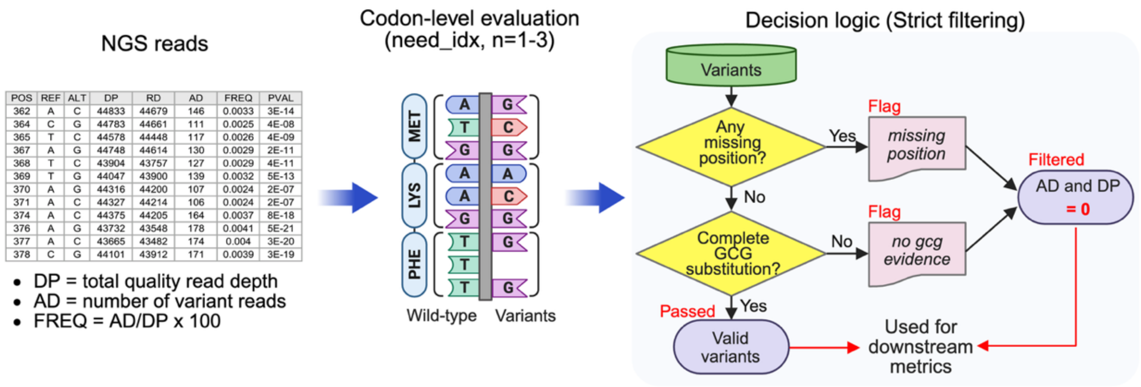


**Fig. S4. NGS strict filtering and variant calling pipeline.**
Overview of raw NGS data processing for alanine (GCG) scanning libraries. Codon-level evaluation was restricted to positions where the wild-type codon differed from the target codon (*need_idx*). Decision logic for variant validity, including handling of missing positional coverage and incomplete GCG evidence, is illustrated. The pipeline outputs a filtered variant table with explicit flags for downstream analysis.

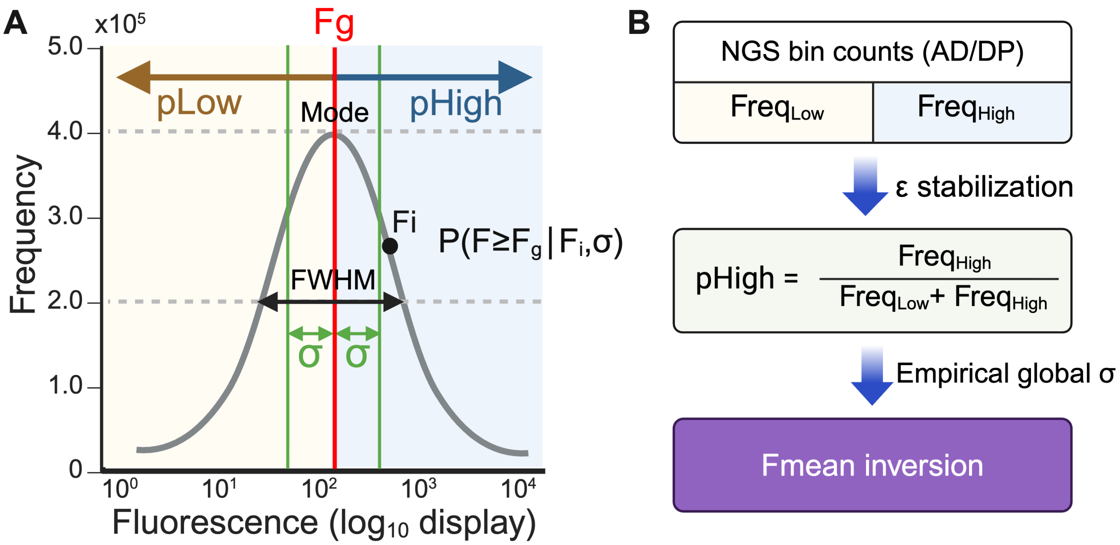


**Fig. S5. Inference of mean fluorescence from sequencing-based bin counts under a log-normal gating model.**

**(A)** Schematic representation of the log-normal fluorescence distribution assumed for individual variants. The distribution is characterized by a mode and full width at half maximum (FWHM), from which the log-space width parameter σ was derived. The fluorescence gate threshold $F_g$ partitions the distribution into two regions corresponding to cells below and above the gate, defining the probabilities $P_{Low} = P(F < F_g)$ and $P_{High} = P(F \geq F_g)$. The shaded regions represent these probabilities under the log-normal model.

**(B)** Computational workflow for estimating the mean fluorescence of variant *i* ($F_{mean,\ i}$) from sequencing-based bin counts. NGS-derived bin frequencies were numerically stabilized using ε, the fraction above the gate was computed, and $F_{mean,\ i}$ was obtained by inverting the log-normal gating relationship (Eq. 7) using a global empirically estimated σ.

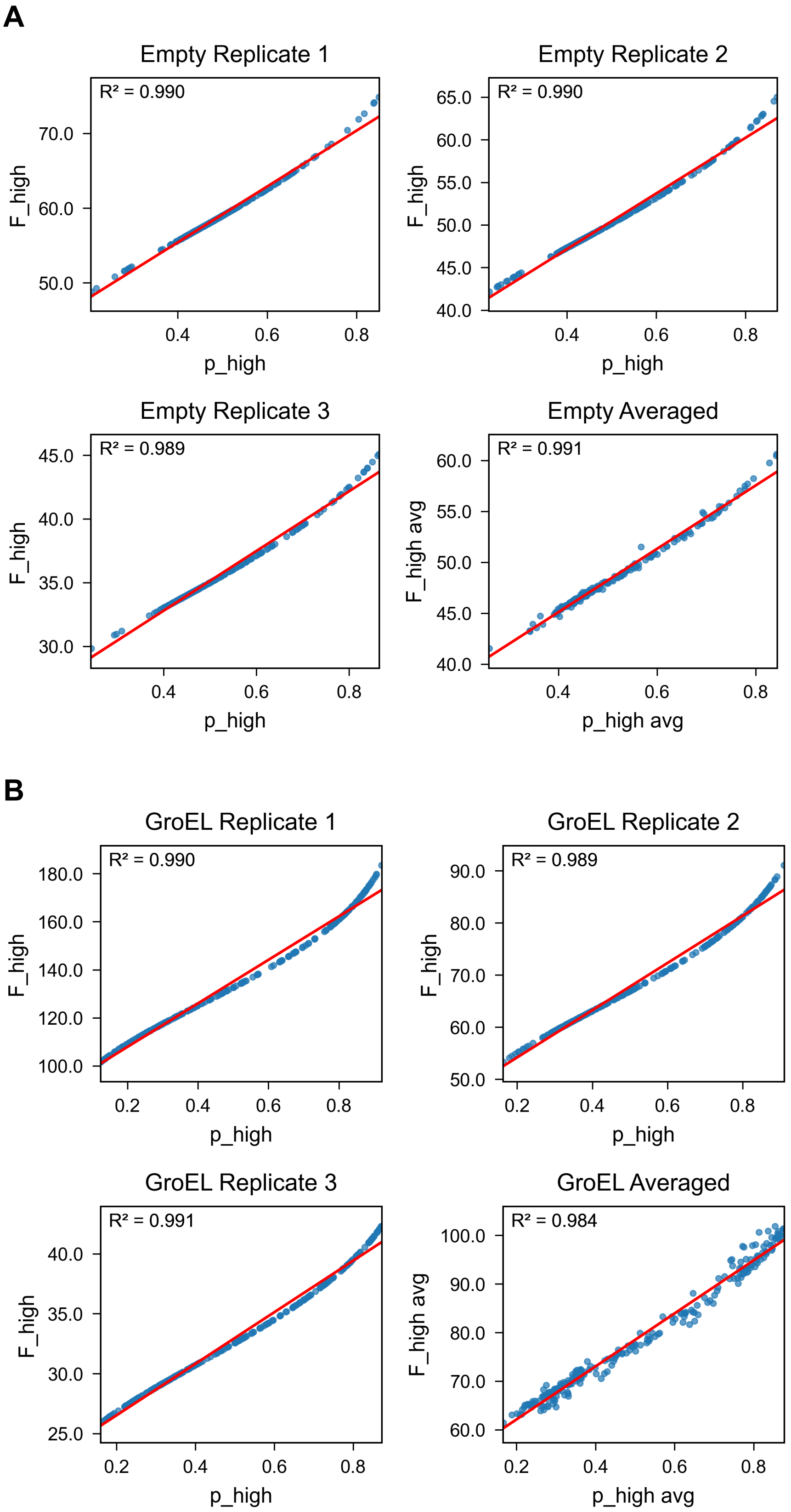


**(Fig. S6. Continued)**

**C**

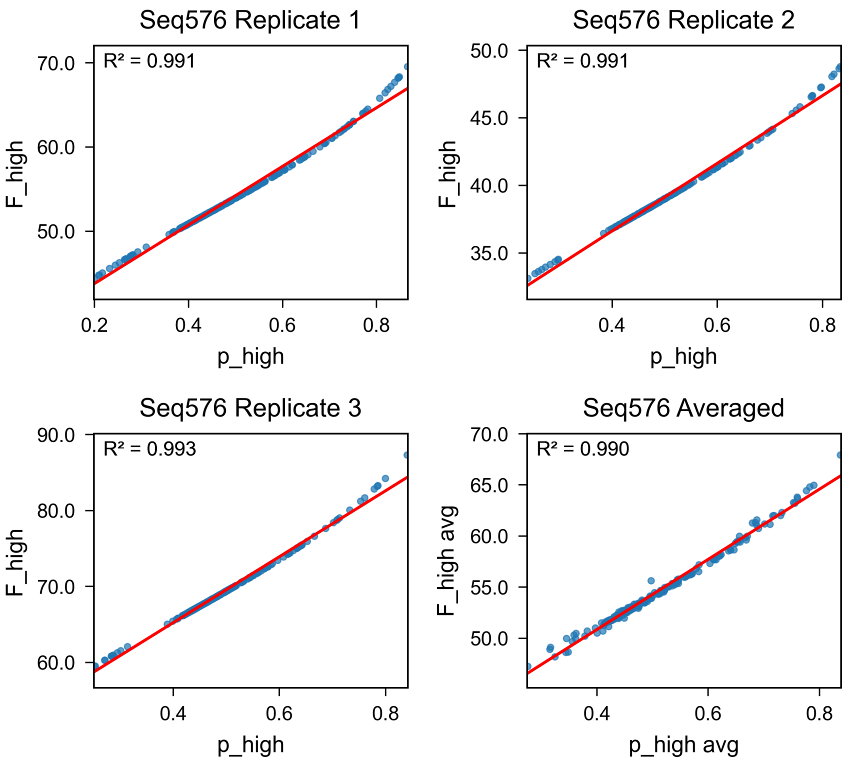


**D**

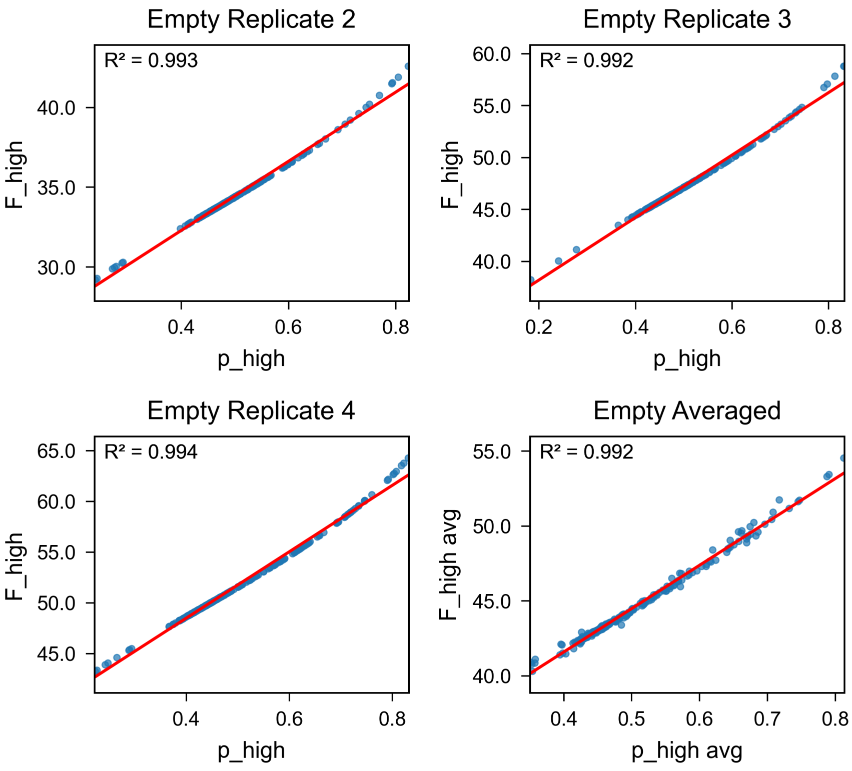


**(Fig. S6. Continued)**

**E**

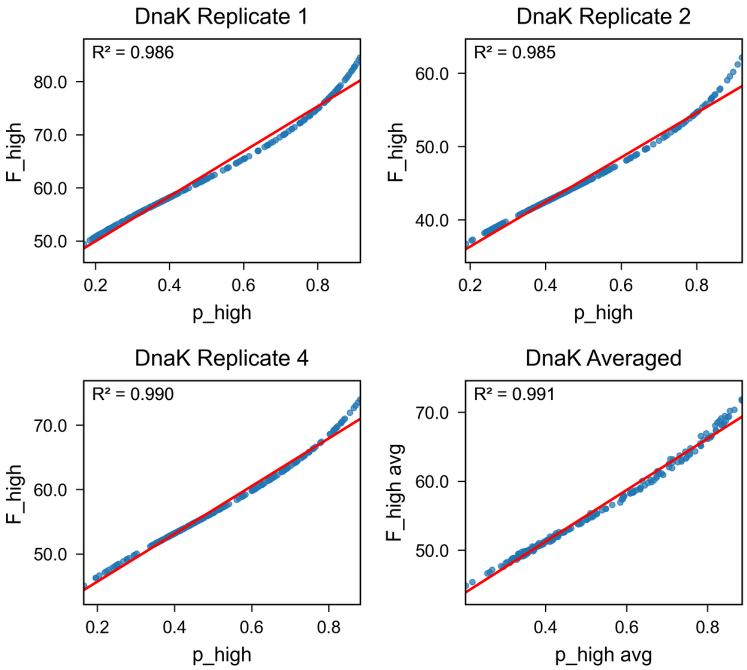


**F**

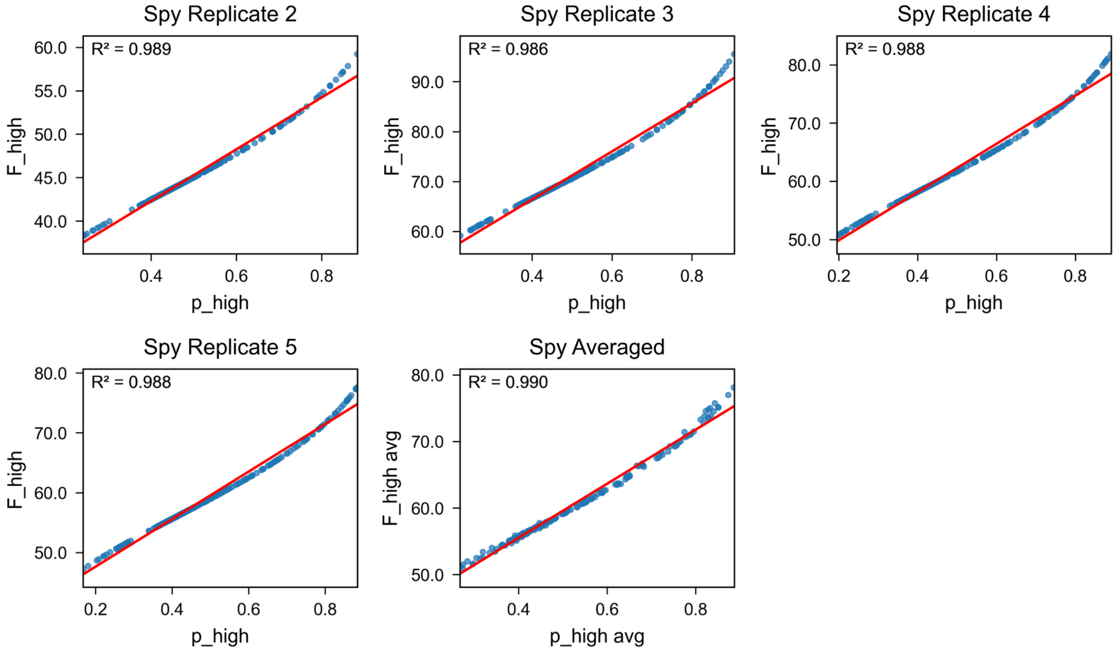


**Fig. S6. Relationship between pHigh and Fmean across conditions.**
Scatter plots of inferred Fmean values as a function of pHigh for each round and condition: (**A**) Round 1 Empty, (**B**) Round 1 GroEL, (**C**) Round 1 Seq576, (**D**) Round 2 Empty, (**E**) Round 2 DnaK, and (**F**) Round 2 Spy. Linear regression fits (red lines) are overlaid, and corresponding $R^2$ values are indicated. Across all conditions, Fmean increased monotonically with pHigh, as expected from the analytical inversion of the log-normal gating model. Deviations from linearity at extreme pHigh values are consistent with the nonlinear form of the inversion and boundary effects introduced by probability stabilization.

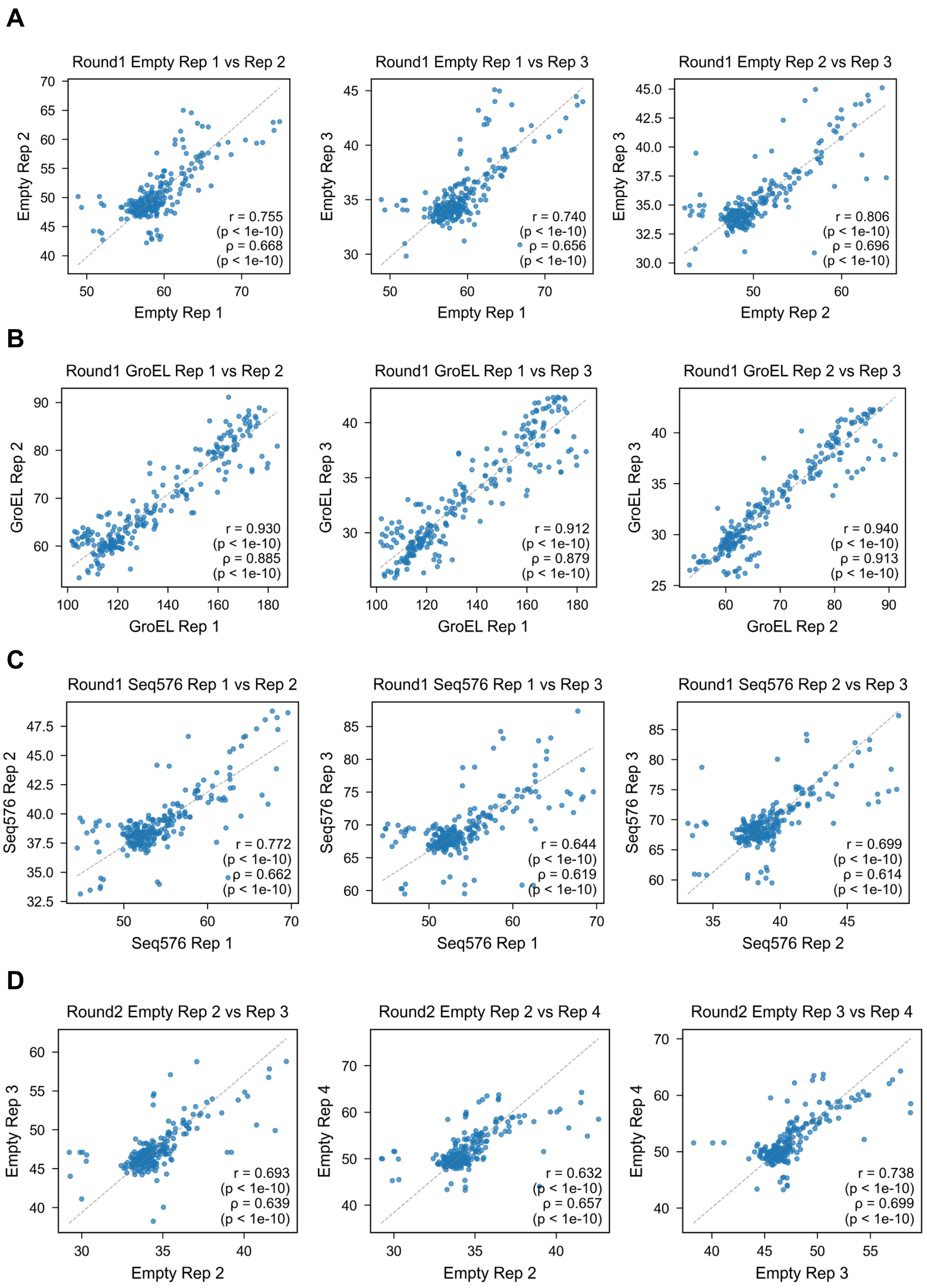


**(Fig. S7. Continued)**

**E**

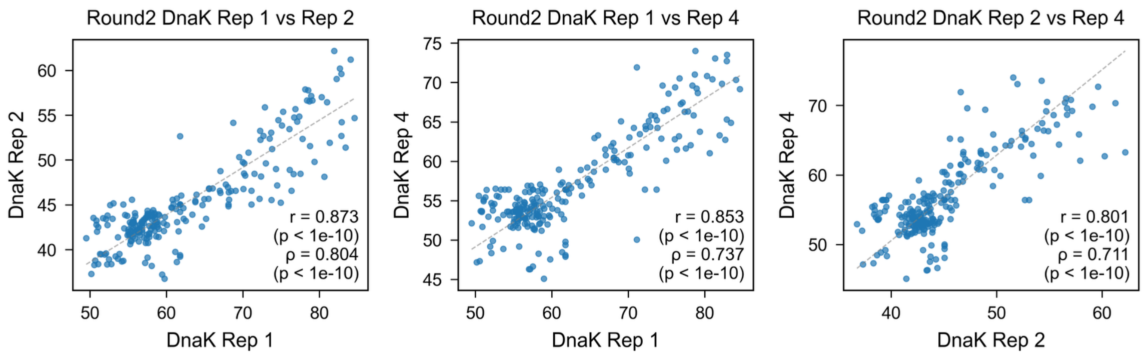


**F**

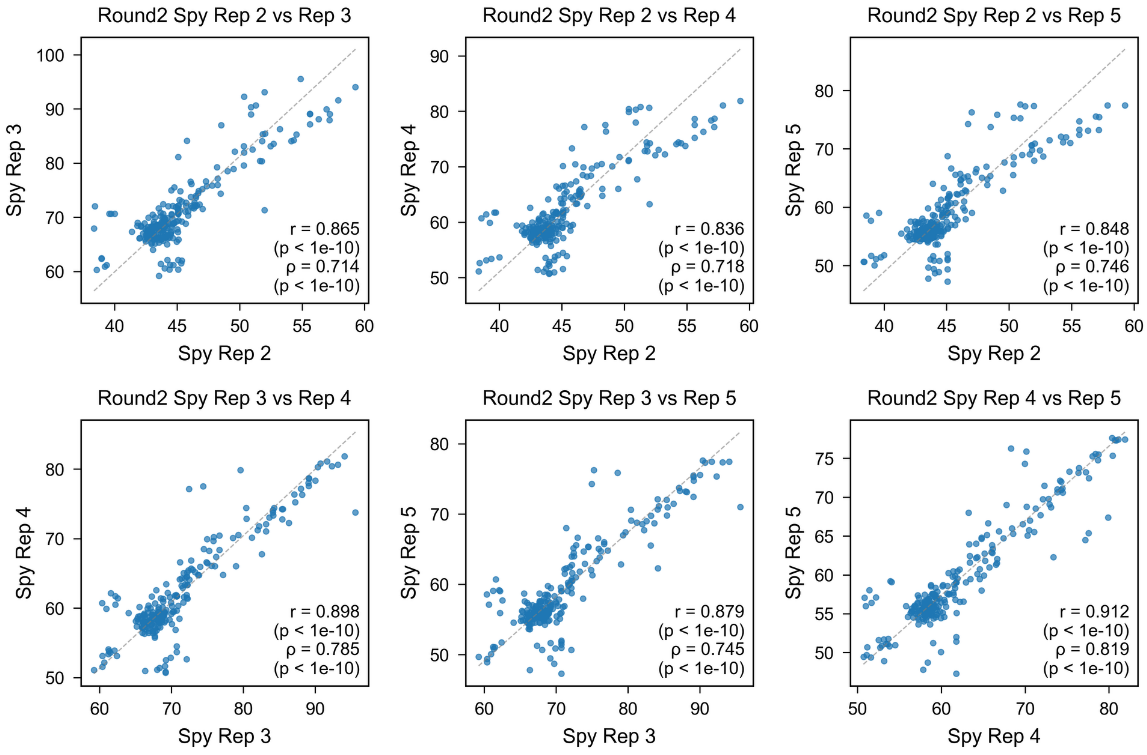


**Fig. S7. Replicate reproducibility across all conditions.**
Pairwise comparisons of inferred Fmean values between biological replicates for each round and condition: (**A**) Round 1 Empty, (**B**) Round 1 GroEL, (**C**) Round 1 Seq576, (**D**) Round 2 Empty, (**E**) Round 2 DnaK, and (**F**) Round 2 Spy. Pearson (*r*) and Spearman (*ρ*) correlation coefficients with corresponding p-values are indicated in each panel. The dashed line represents the identity line (slope = 1). Detailed regression statistics, including slope and residual dispersion, are summarized in Table S6.

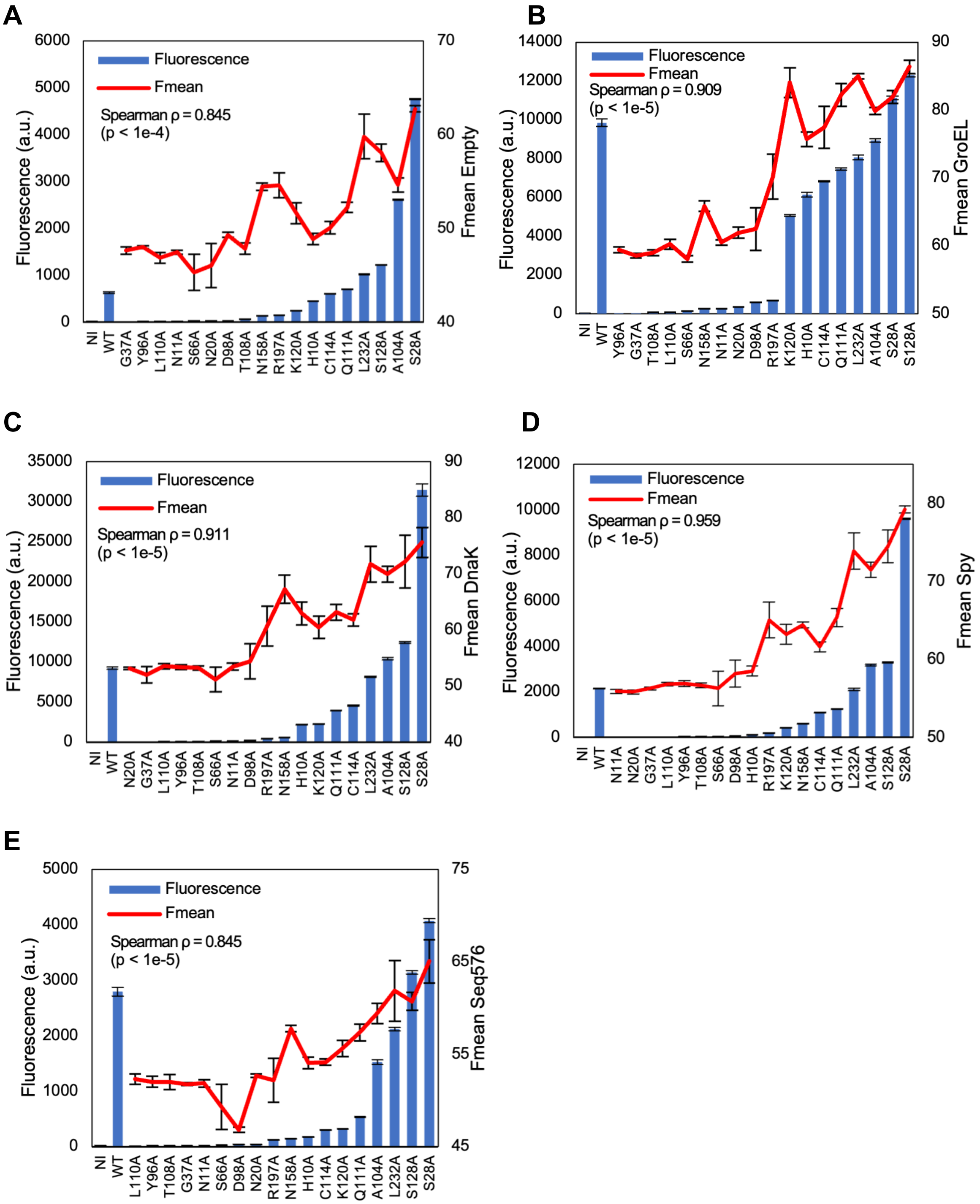


**Fig. S8. Orthogonal validation of CHAP-SEQ-inferred mean fluorescence.**
Eighteen individual alanine substitution variants selected from low- and high-bin populations were generated by site-directed mutagenesis and co-expressed with the indicated chaperones. Plate reader fluorescence values (blue bars; mean ± SEM) were compared to CHAP-SEQ-inferred Fmean values (red lines). Spearman rank correlation coefficients (ρ) and associated p-values are indicated for each condition (A, Empty; B, GroEL; C, DnaK; D, Spy; E, Seq576). Measurements were performed in biological triplicate (Spy in quadruplicate). NI and WT indicate non-induced and wild type, respectively.

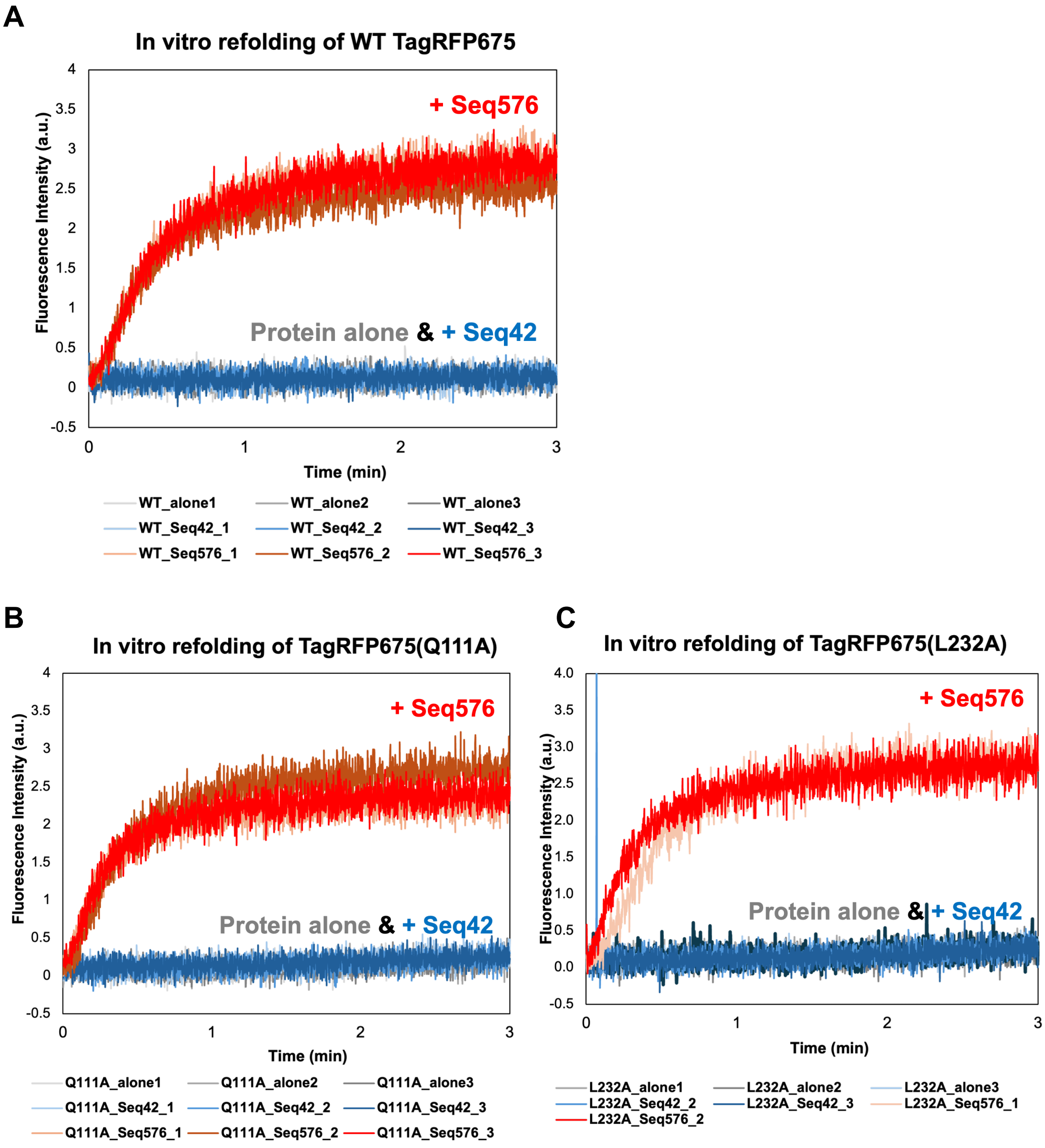


**Fig. S9. Seq576 enhances *in vitro* refolding of TagRFP675 variants.**
Chemically denatured wild-type TagRFP675 (**A**), TagRFP675(Q111A) (**B**), and TagRFP675(L232A) (**C**) (final concentration 0.5 μM) were diluted into refolding buffer in the presence of G4 DNA (Seq576) at a 1:2 molar ratio (protein:DNA) or a non-G4 control (Seq42) under identical conditions. Native fluorescence recovery (598/675 nm) was monitored over time. Seq576 enhanced fluorescence recovery relative to control conditions, consistent with enhanced recovery of the native fluorescent state.

A

B

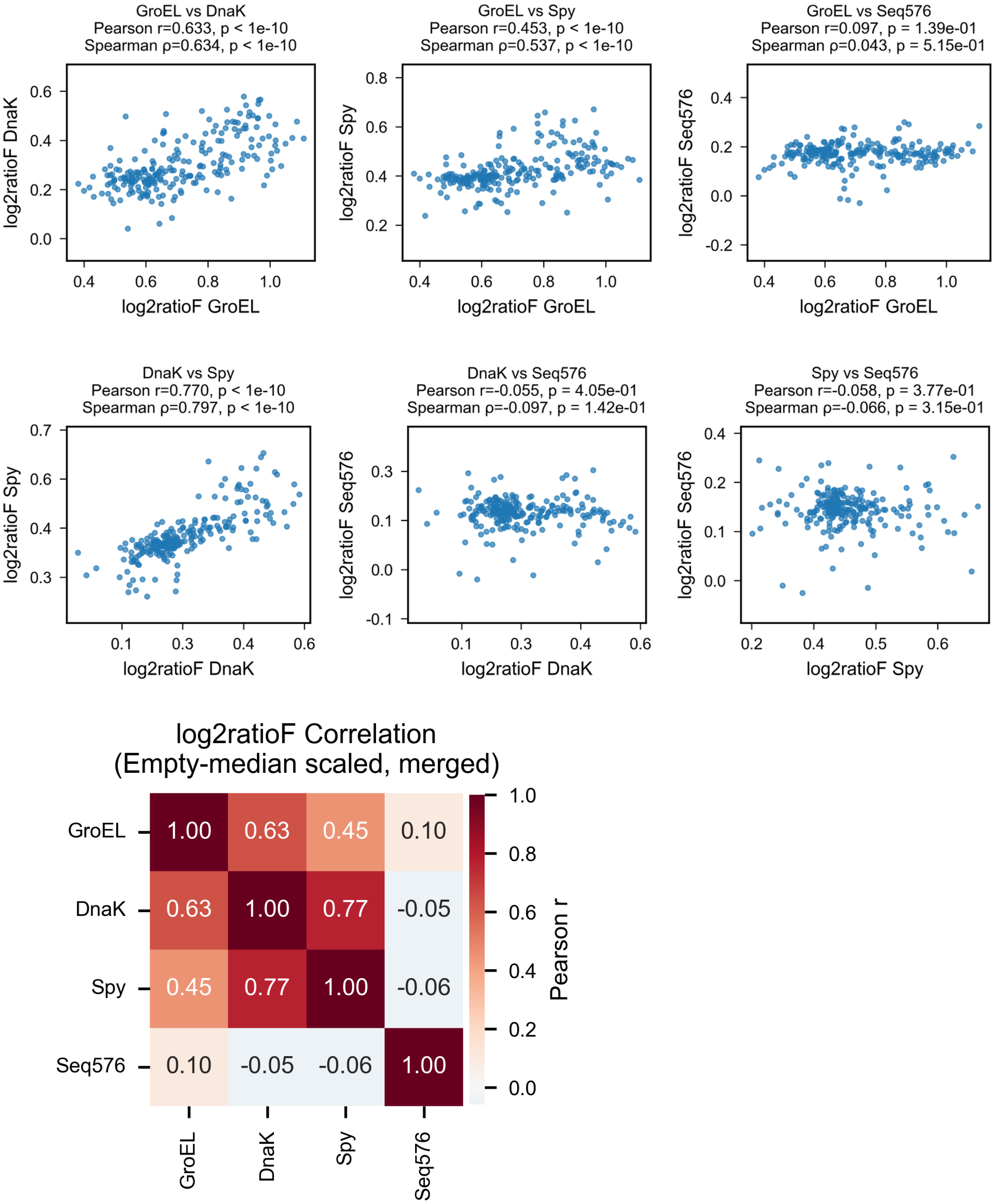


**Fig. S10. Pairwise residue-level $log_2$ratioF correlations.**
(**A**) All pairwise scatter plots corresponding to the correlation matrix shown in Fig. 1E. Each panel shows residue-level $log_2$ratioF values between two conditions, with Pearson and Spearman correlation coefficients indicated. For cross-round comparisons, Round 2 Fmean values were median-scaled to the Round 1 Empty baseline prior to ratio calculation. (**B**) Pearson correlation matrix of residue-level $log_2$ratioF values across chaperone conditions.

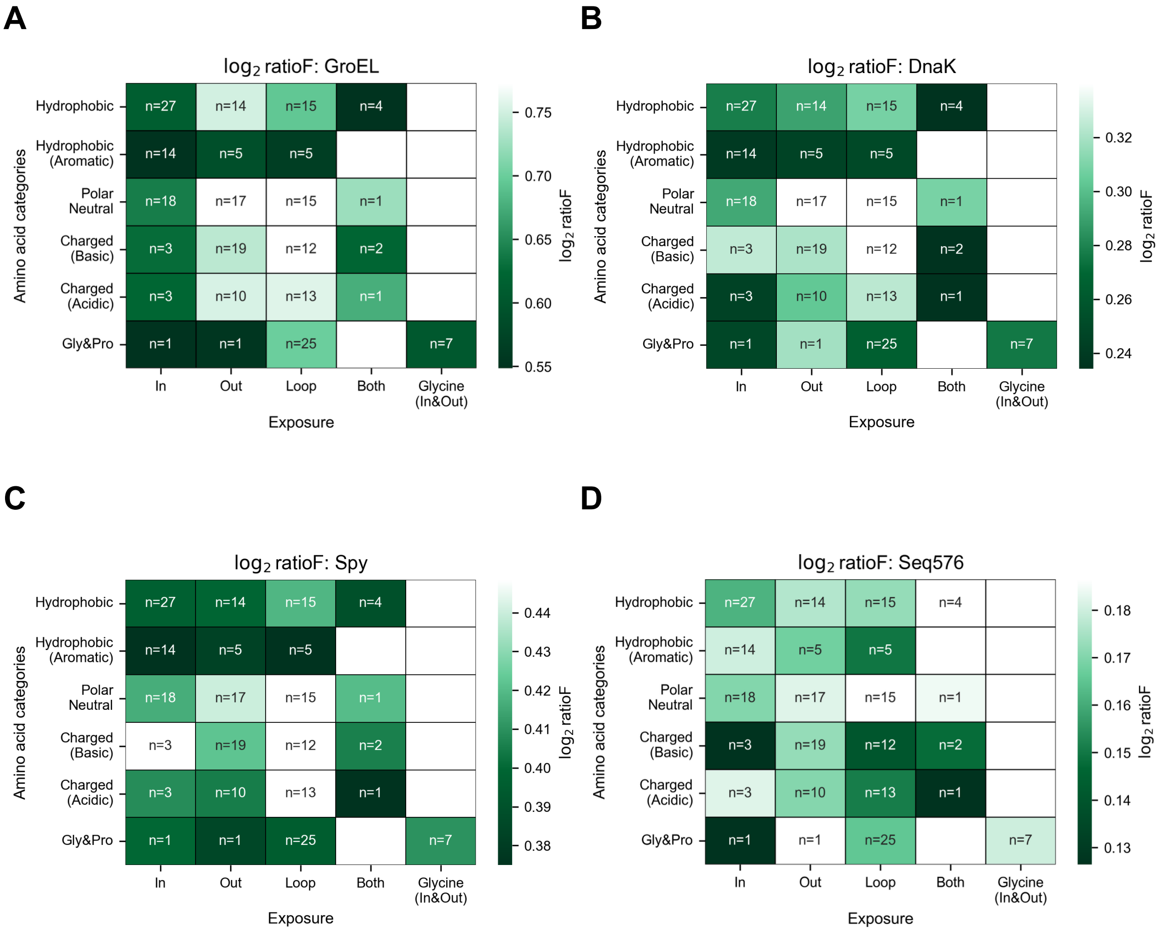


**Fig. S11. Combinatorial amino acid and exposure heatmap.**
Heatmap representation of $log_2$ratioF values grouped by the combination of amino acid category and structural exposure class under (**A**) GroEL, (**B**) DnaK, (**C**) Spy, and (**D**) Seq576 conditions. Residues were stratified according to physicochemical class (hydrophobic, hydrophobic aromatic, polar neutral, charged basic, charged acidic, glycine/proline) and exposure category (buried, surface-exposed, loop, both, glycine-specific where applicable). Each cell represents the mean $log_2$ratioF value of residues belonging to the corresponding category combination. Color intensity reflects the magnitude of the mean effect, with darker shades indicating residues with lower $log_2$ratioF values and therefore greater dependence on chaperone-assisted folding. The number of residues (n) in each category combination is indicated within each cell. Empty cells represent category combinations not present in the structure. A loop-specific follow-up analysis showed that charged residues exhibited greater relative Seq576 dependence than non-charged residues (Mann–Whitney $p = 0.0021$; permutation $p = 0.0019$; see Supplementary Methods and Results).

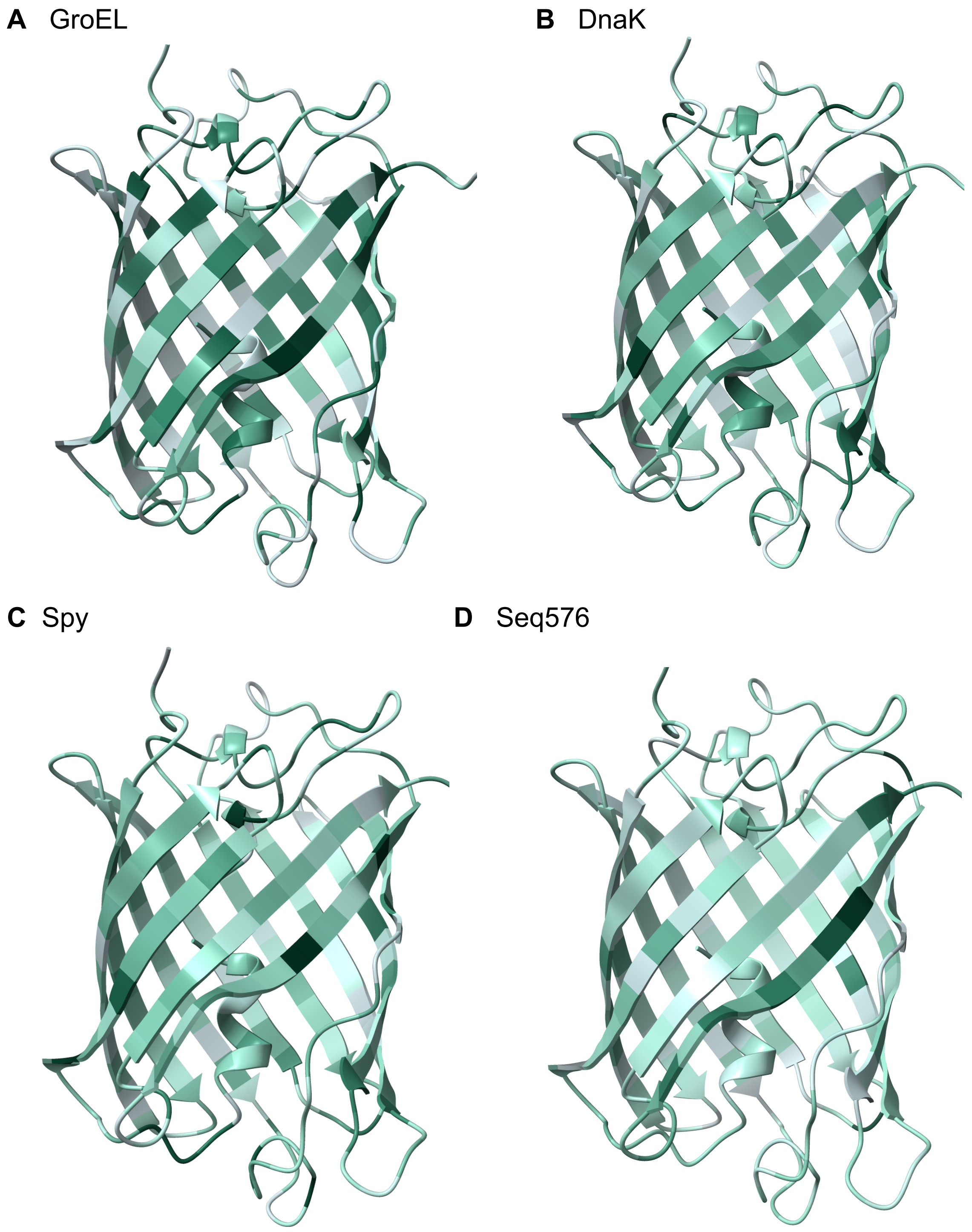


**Fig. S12. Comparative structural mapping across chaperones.**
(**A**–**D**) Projection of residue-level $log_2$ratioF values onto the TagRFP675 structure under GroEL (**A**), DnaK (**B**), Spy (**C**), and Seq576 (**D**) conditions. Color scaling was performed independently within each condition to visualize relative residue-level patterns and therefore does not represent absolute cross-condition differences in $log_2$ratioF magnitude. Rotating structural visualizations corresponding to GroEL, DnaK, Spy, and Seq576 conditions are provided in Movies S1–S4, respectively.

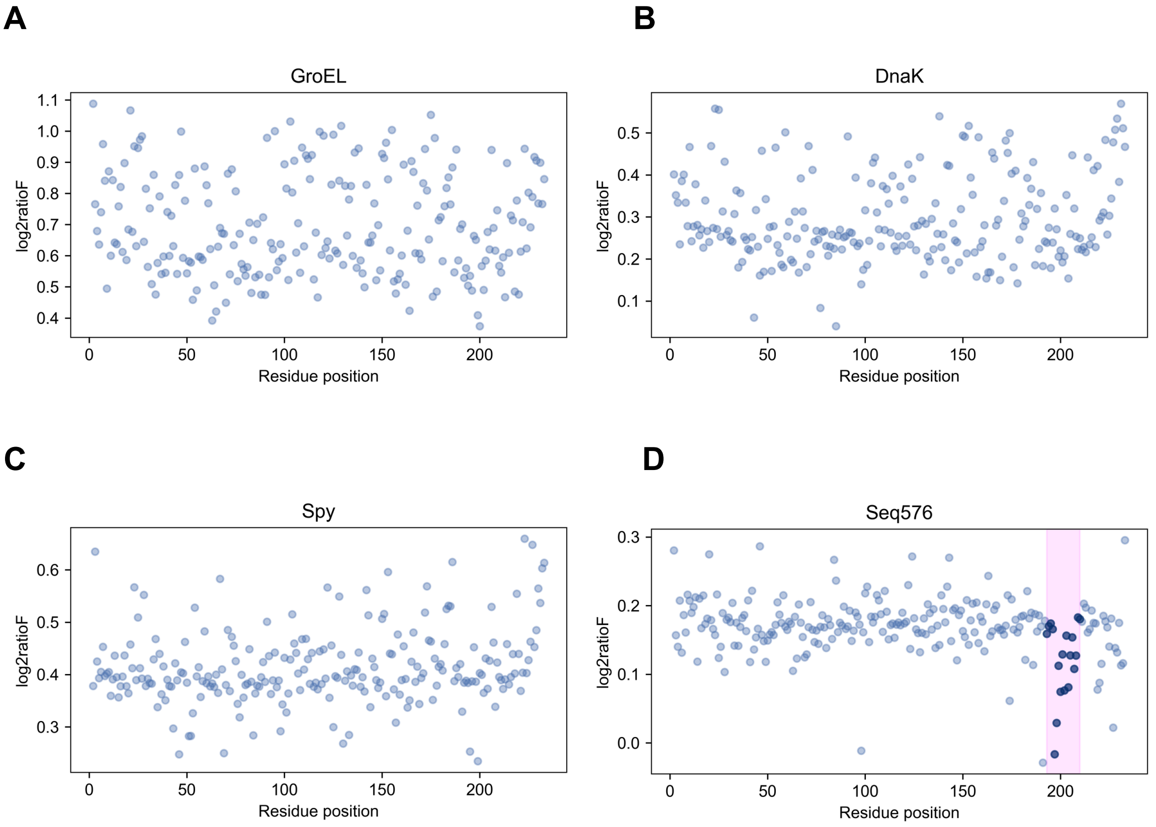


**Fig. S13. Residue-level distribution of log₂ratioF across chaperone conditions.**
Scatter plots showing $log_2$ratioF values as a function of residue position for GroEL, DnaK, Spy, and Seq576. Each dot represents the $log_2$ratioF of an individual residue. These plots correspond to the residue-level relationships summarized in the correlation matrix (Fig. 3E) and provide a one-dimensional view of folding responses across the sequence. Among the conditions, only Seq576 exhibits a significant contiguous sequence clustering, consistent with the statistically significant contiguous region identified by sliding-window and permutation analyses (Fig. 3E–F). The region spanning residues 193–210, including the core cluster at residues 197–202, is highlighted in magenta.

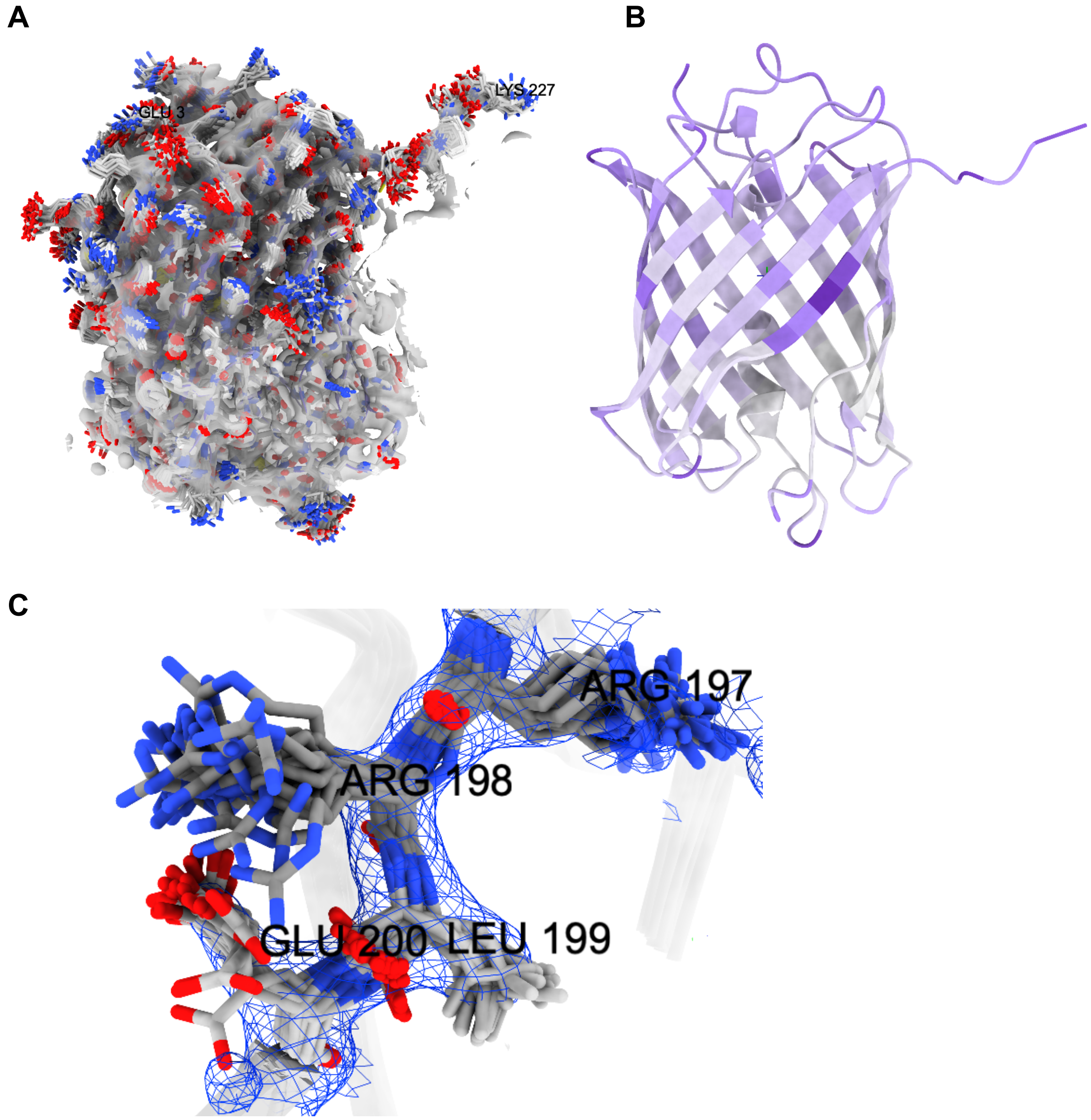


**Fig. S14. Structural dynamics of TagRFP675 and localization of the Seq576-specific hotspot.** (**A**) Structural ensemble representation of the wild-type TagRFP675 protein used for RMSD calculation. (**B**) Residue-level RMSD values mapped onto the TagRFP675 structure using the B-factor field. Residues with higher RMSD values are shown in darker purple, indicating greater positional variability across the structural ensemble. (**C**) Close-up view of the 193–210 region identified by sliding-window and permutation analyses as important for Seq576 function. Ensemble representations illustrate conformational variability of residues within this region, including residues 197–200, which exhibit highly elevated RMSD values. Rotating structural visualizations of (**A**) and (**B**) are provided in Movies S5–S6, respectively.

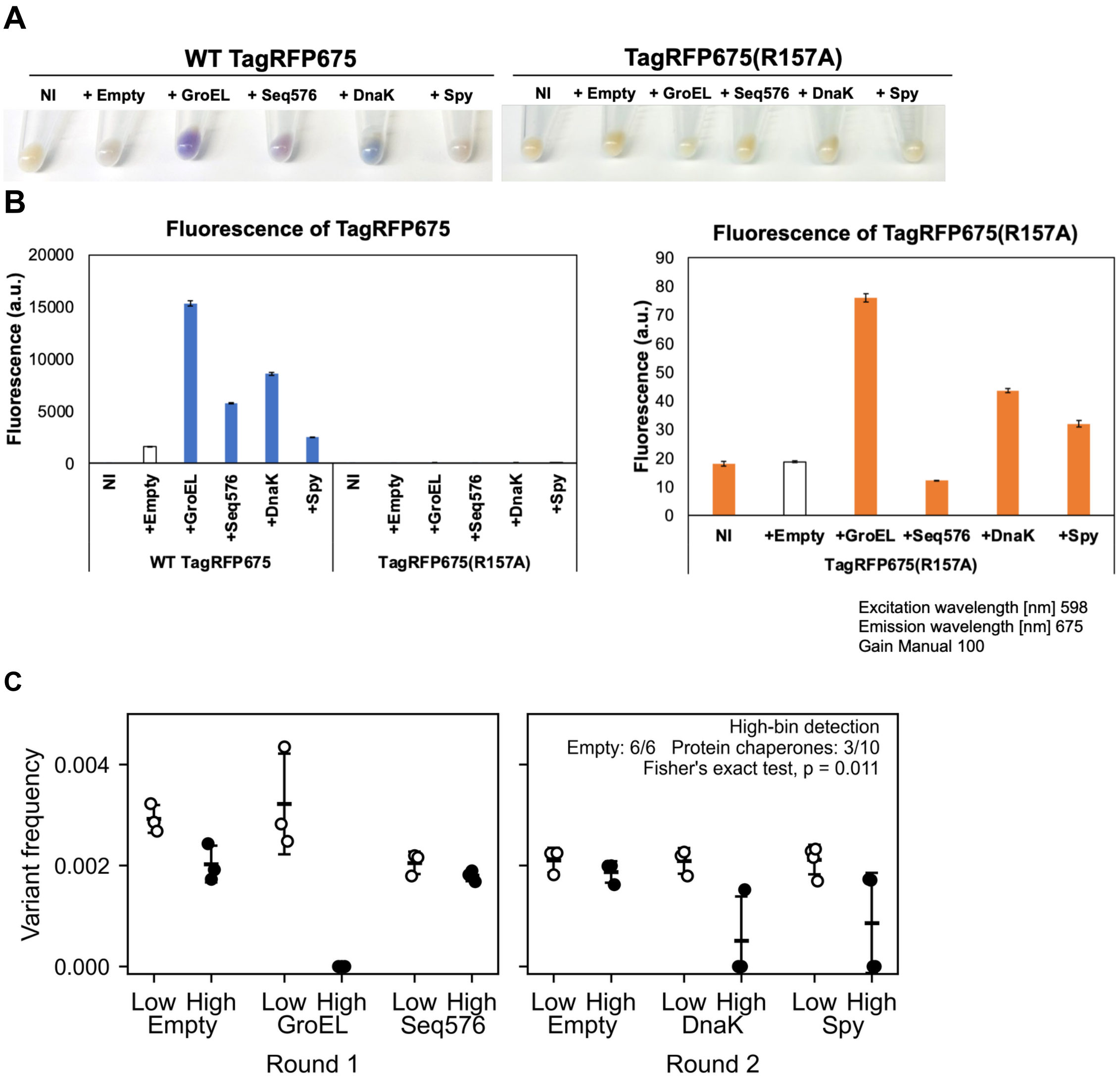


**Fig. S15. Follow-up characterization of R157A.**

(**A**) Cell pellet color comparison of WT TagRFP675 and TagRFP675(R157A) under the indicated Empty and chaperone conditions. (**B**) Bulk fluorescence measurements of WT TagRFP675 and TagRFP675(R157A) under the indicated conditions. R157A exhibited strongly reduced fluorescence relative to WT (left), although GroEL and DnaK increased R157A fluorescence to some extent (right). (**C**) Distribution of R157A variant frequencies between the Low- and High-fluorescence bins in CHAP-seq Round 1 and Round 2. Open circles indicate Low-bin frequencies and filled circles indicate High-bin frequencies; horizontal bars and error bars indicate mean ± SD. Samples in which R157A was below the predefined detection threshold were assigned a variant frequency of zero. R157A was detected in 3/10 High-bin samples across the protein-chaperone conditions (GroEL, DnaK, and Spy), compared with 6/6 Empty High-bin samples (two-sided Fisher's exact test, $p = 0.011$).

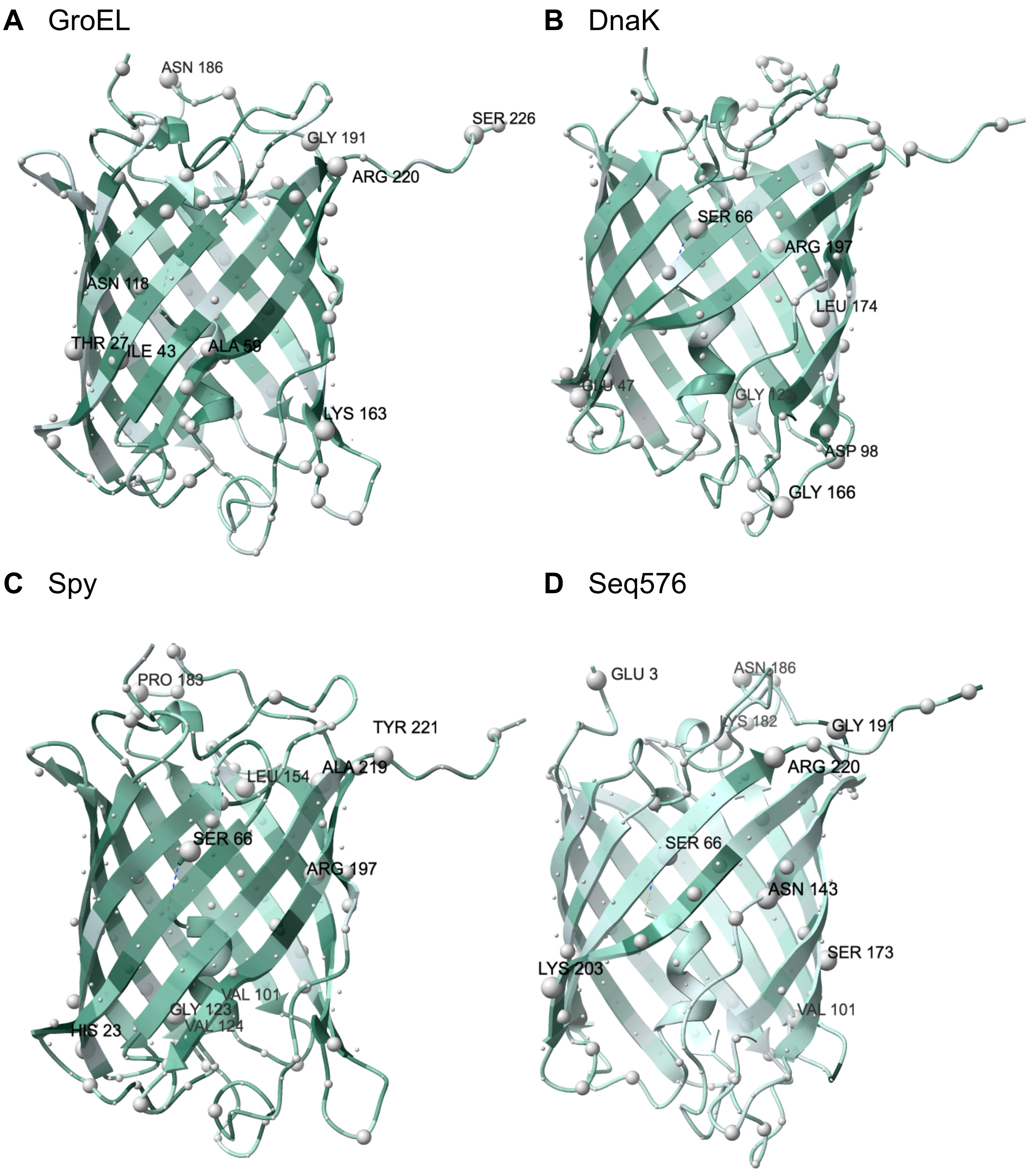


**Fig. S16. Structural distribution of residue-level propagated variability.**
(**A**–**D**) Structural mapping of residue-level sd_log2 under (**A**) GroEL, (**B**) DnaK, (**C**) Spy, and (**D**) Seq576 conditions. Residues were stratified into five distribution-based categories (QD1–QD5), where QD5 represents sd_log2 ≥ $\mu + 2\sigma$ within each condition. Sphere size reflects propagated between-replicate variability, while ribbon coloring represents $\log_2$ratioF. Rotating structural visualizations corresponding to GroEL, DnaK, Spy, and Seq576 conditions are provided in Movies S9–S12, respectively.

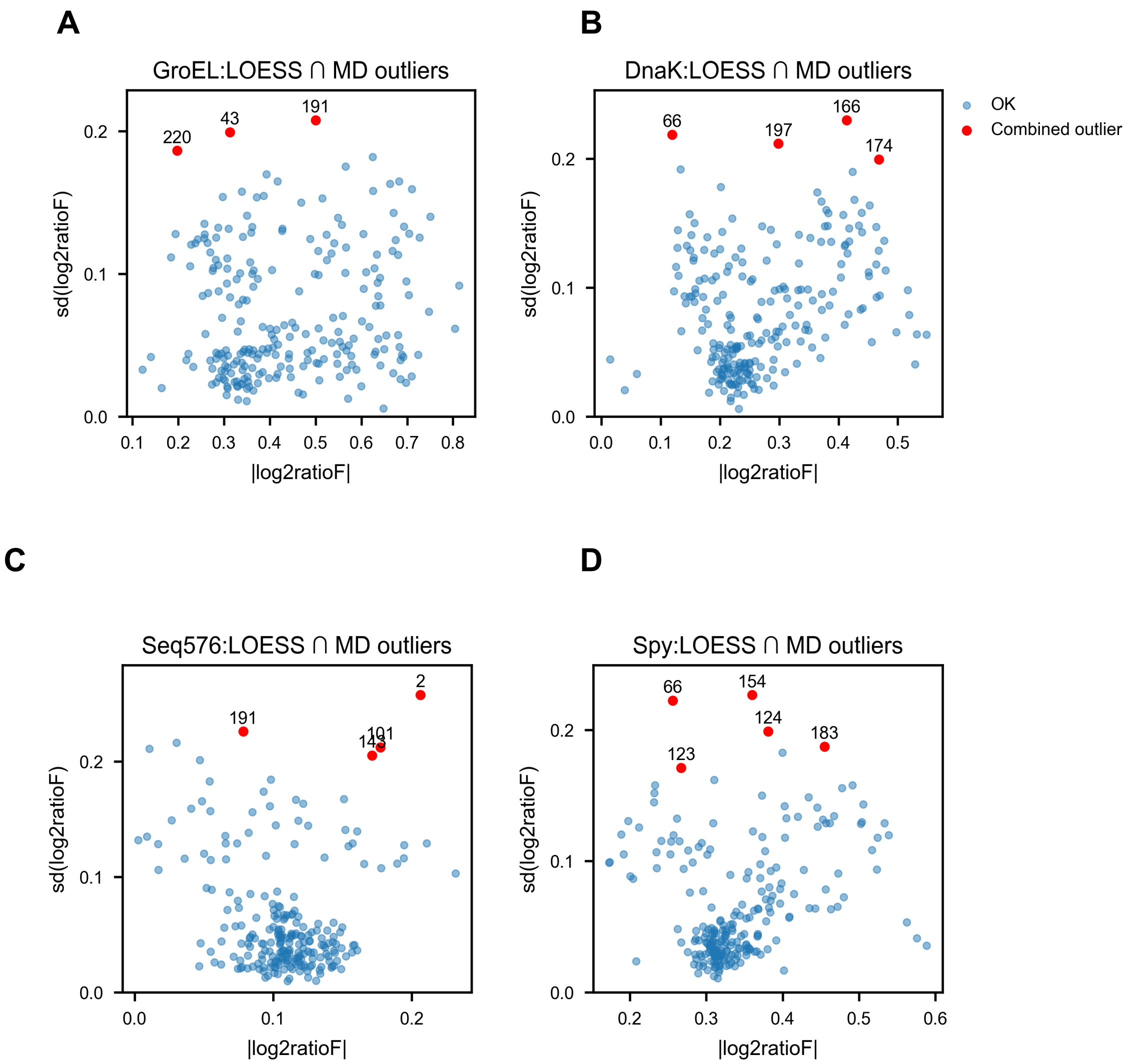


**(Fig. S17. Continued)**

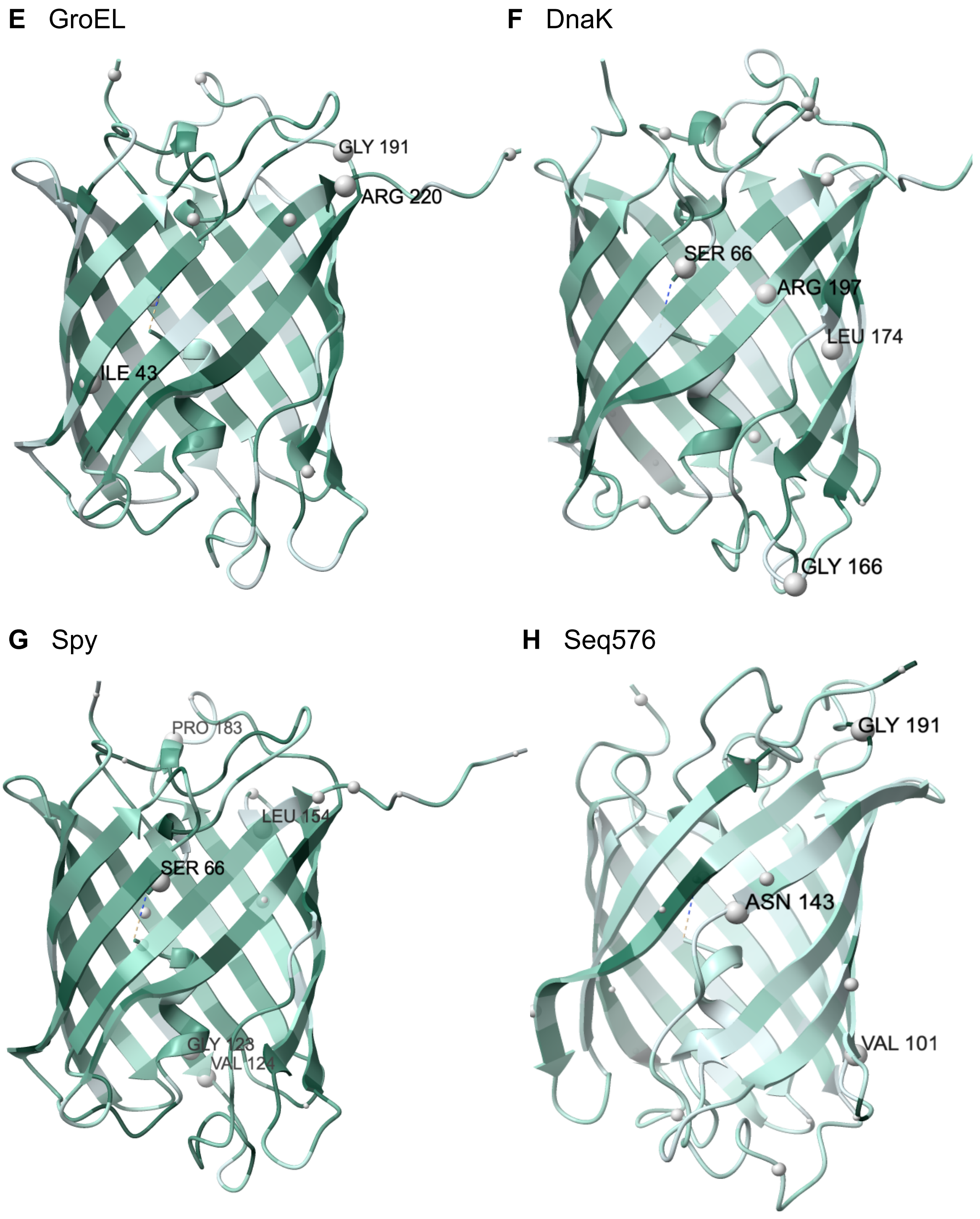


**Fig. S17. Statistical identification of high-variability residues.**
(**A**–**D**) Scatter plots of sd_log2 versus |$\log_2$ratioF| with combined outliers (LOESS ∩ Mahalanobis) highlighted under (**A**) GroEL, (**B**) DnaK, (**C**) Spy, and (**D**) Seq576 conditions. (**E**–**H**) Structural mapping of combined outliers under (**E**) GroEL, (**F**) DnaK, (**G**) Spy, and (**H**) Seq576 conditions. Residues classified as combined outliers exceeded both the LOESS residual threshold (top 5%) and the Mahalanobis distance threshold (97.5$^{th}$ percentile of the $\chi^2$ distribution, df = 2). Rotating structural visualizations of the outlier mappings are provided in Movies S13–S16, respectively.

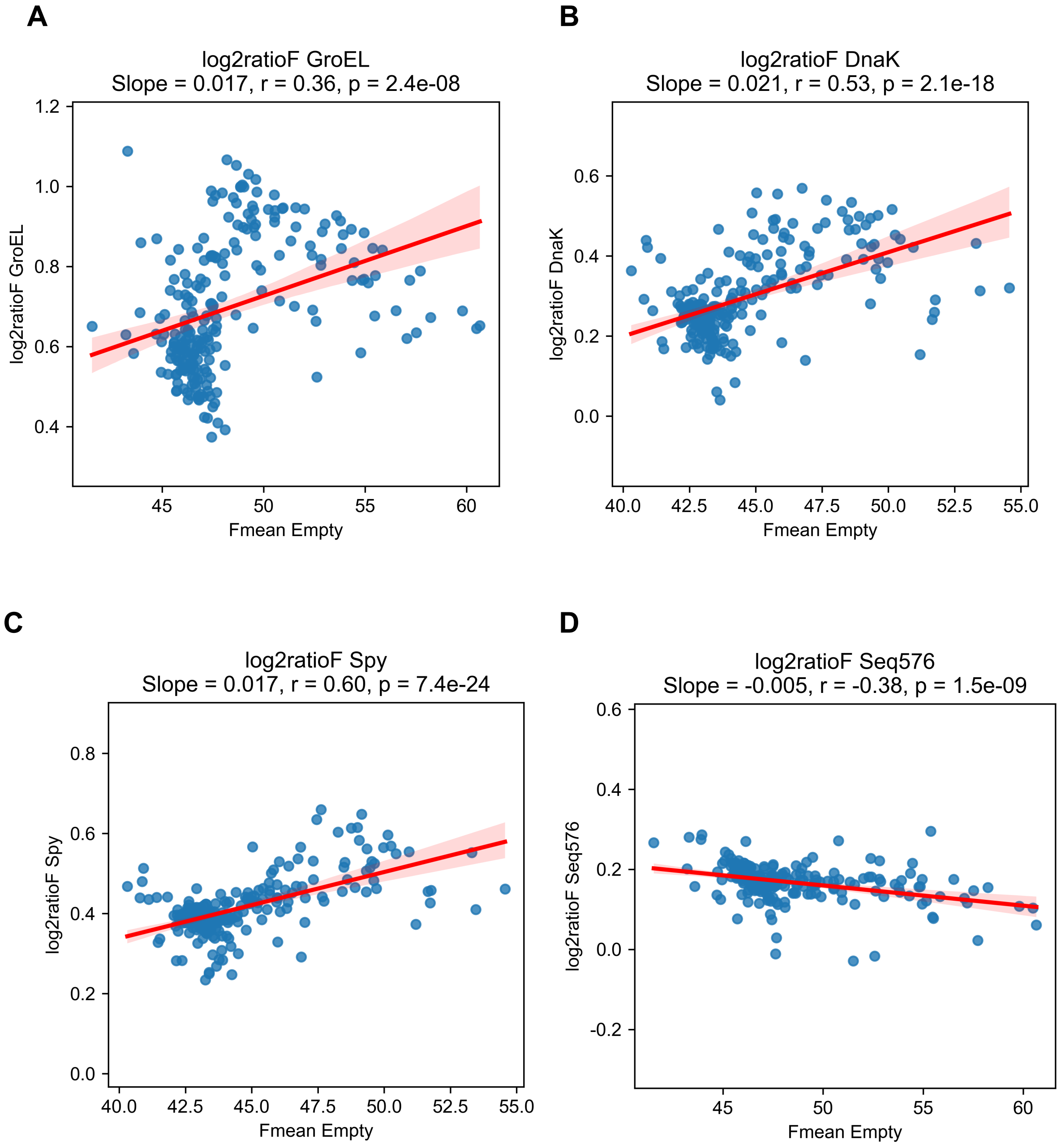


**Fig. S18. Baseline dependence of residue-level folding effects.**
Residue-level $log_2$ratioF values were plotted against baseline folding levels measured under the Empty condition ($Fmean_{Empty}$). Each point represents a single alanine variant. Linear regression lines are shown in red. Protein-based chaperones exhibited positive slopes, indicating larger folding improvements for variants with higher baseline folding levels, whereas Seq576 displayed a weak negative slope, indicating an opposite redistribution trend. Regression slopes, Pearson correlation coefficients, and associated p-values are reported for each panel.

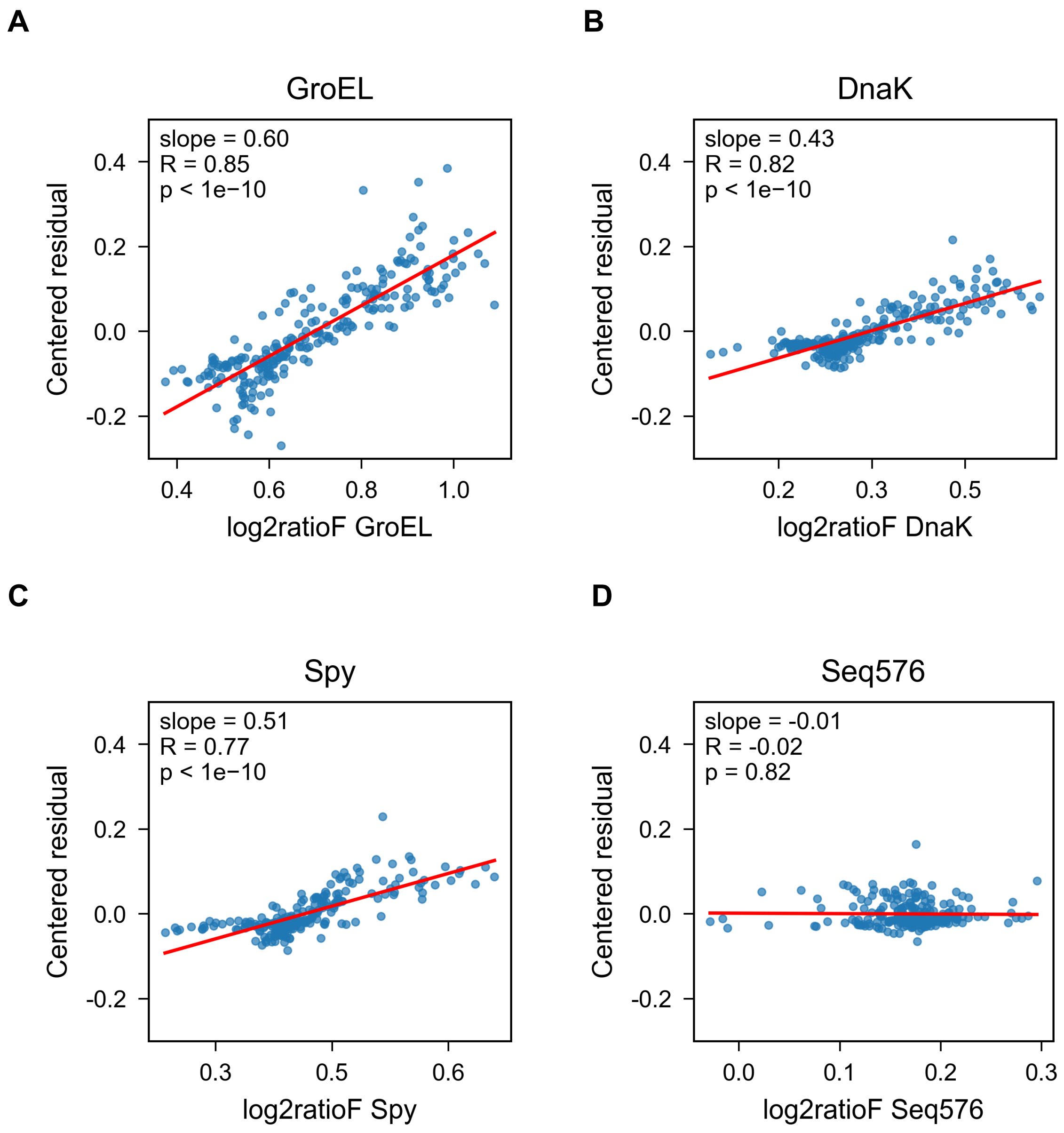


**Fig. S19. Consistency between residual-based and $\log_2$ratioF-based folding metrics.** Centered residual values relative to the Empty baseline were plotted against $\log_2$ratioF for each condition (GroEL (**A**), DnaK (**B**), Spy (**C**), and Seq576 (**D**)). Protein-based chaperones exhibited strong positive correlations, indicating that variants with larger chaperone-dependent folding effects also showed larger positive deviations from the Empty baseline model. Seq576 showed no significant correlation. Red lines indicate linear regression fits. Pearson correlation coefficients and p-values are shown in each panel.

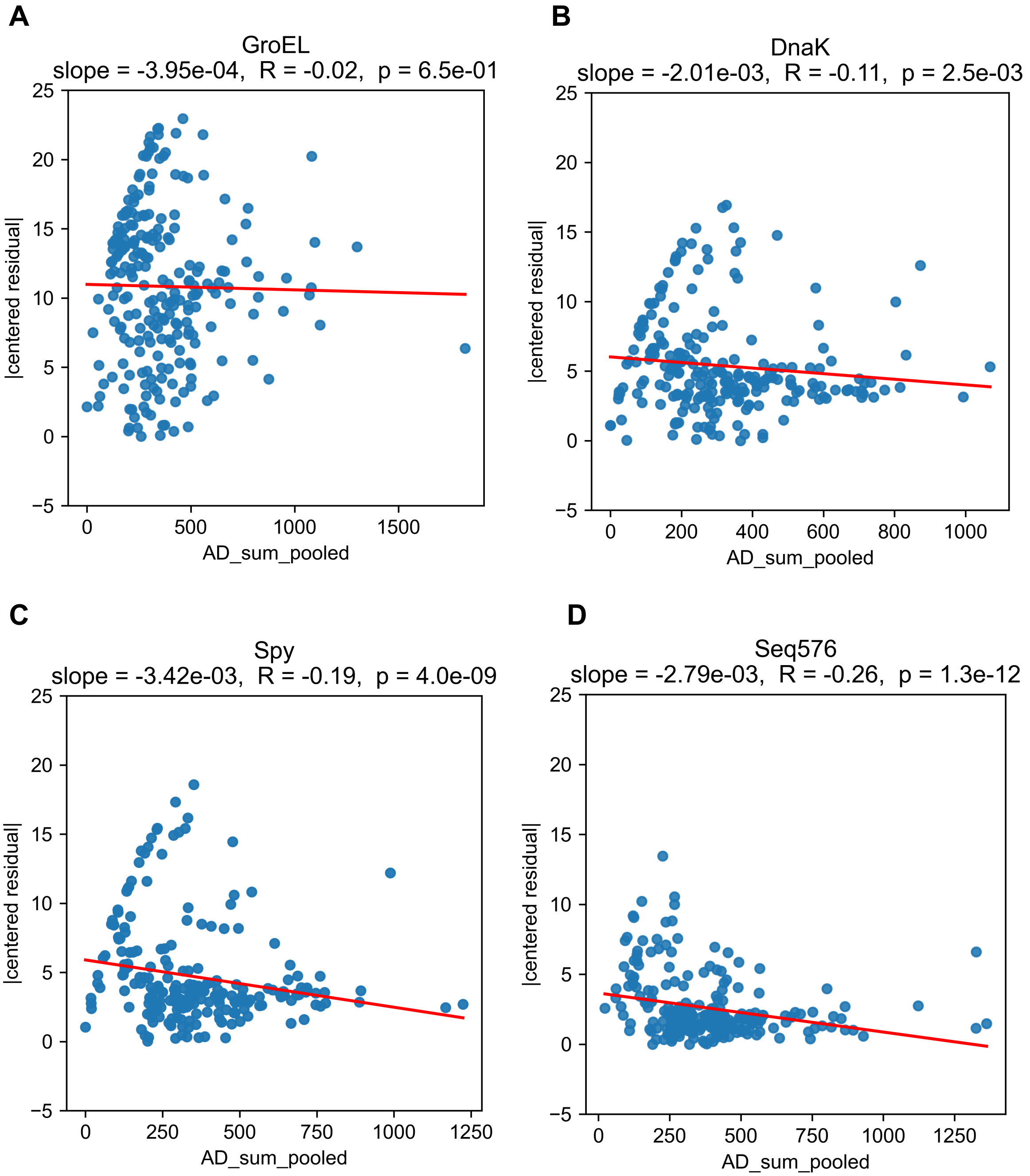


**Fig. S20. Residual dispersion shows little relationship with sequencing depth.**
Absolute centered residuals relative to the Empty baseline were plotted against pooled read depth (AD_sum_pooled) for GroEL (**A**), DnaK (**B**), Spy (**C**), and Seq576 (**D**) conditions. Residuals were mean centered within condition to remove global shifts in fluorescence levels. Linear regression lines and Pearson R values are shown. Correlations were weak, indicating that sequencing depth is unlikely to be a major determinant of the observed residual dispersion.

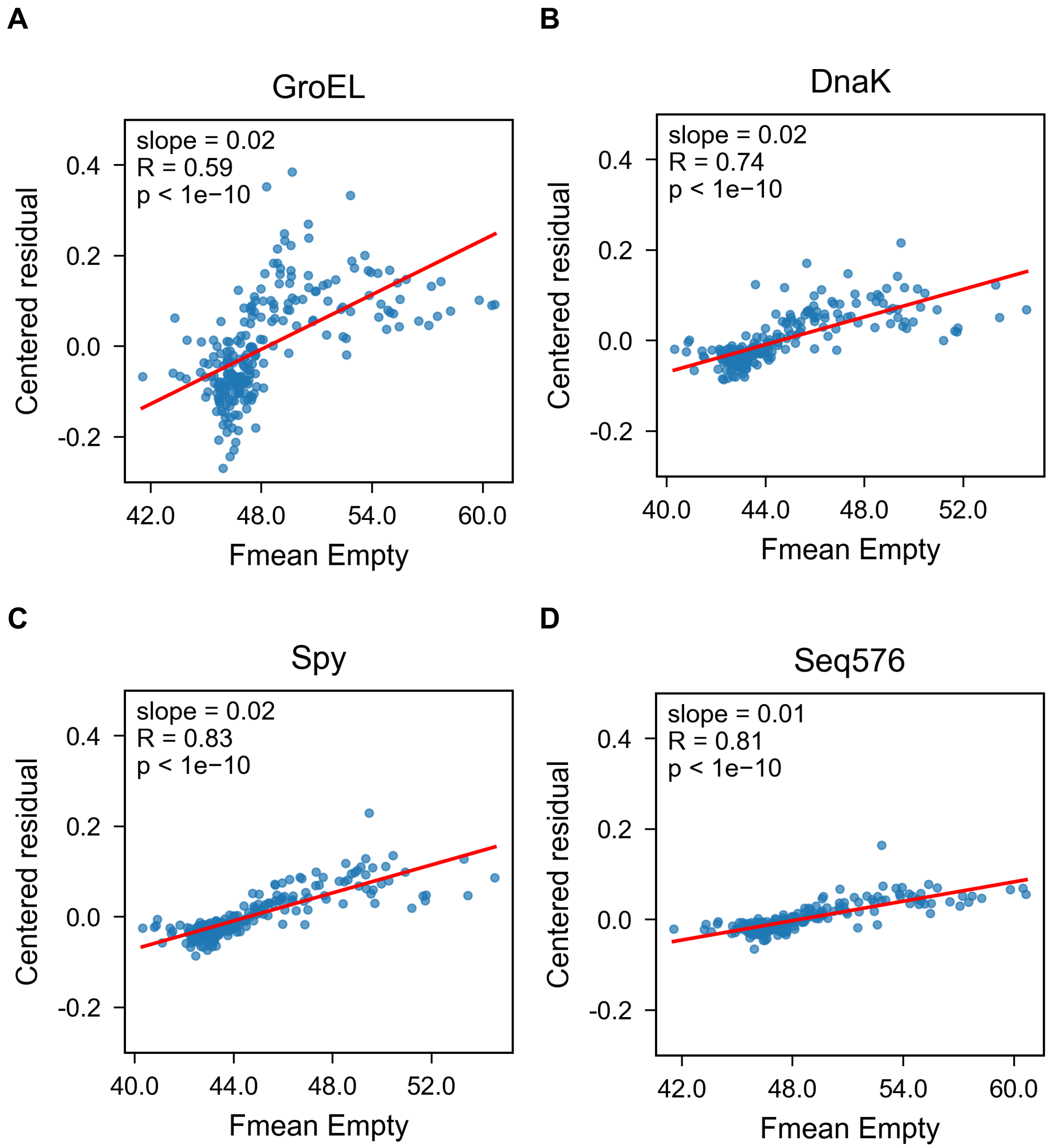


**Fig. S21. Relationship between baseline folding level and Empty-referenced residual deviations.**
Centered residual values relative to the Empty baseline regression were plotted against baseline folding levels (Fmean under the Empty condition) for each chaperone system (GroEL (**A**), DnaK (**B**), Spy (**C**), and Seq576 (**D**)). Residuals were mean centered within each condition to remove global fluorescence shifts. Each point represents the average residual for a single residue across replicates. Linear regression lines are shown in red. Slopes were generally small, indicating only a weak association of residual deviations on baseline foldability.

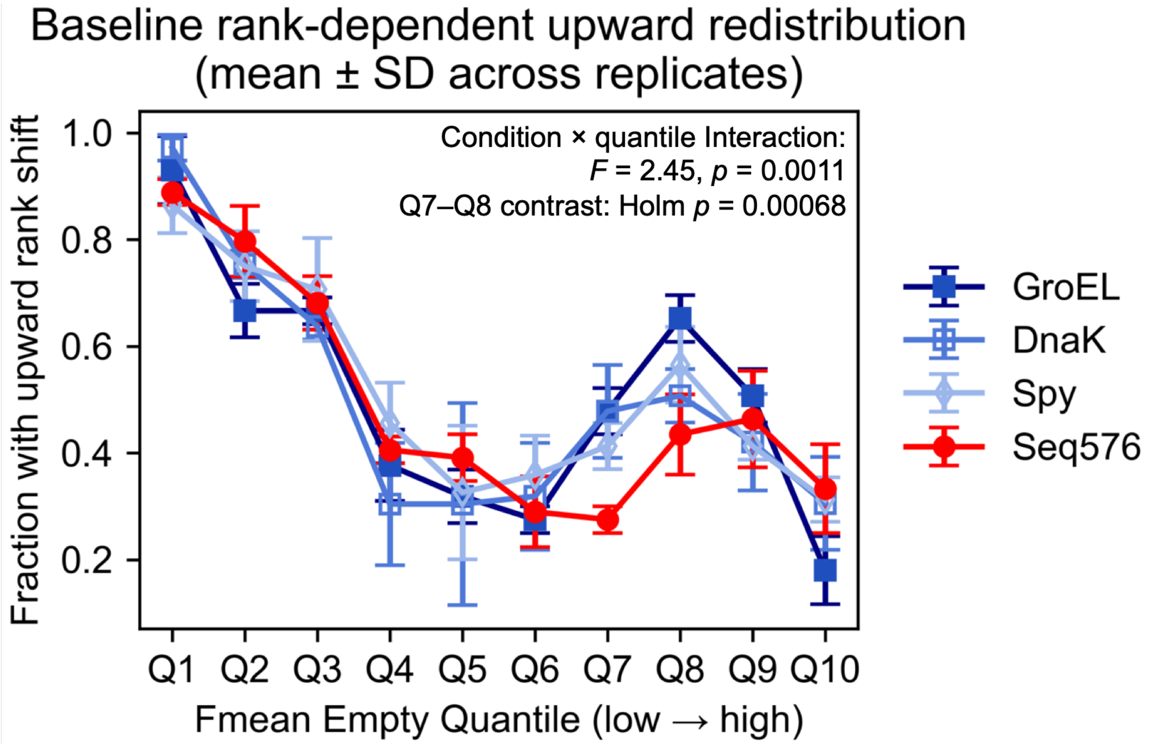


**Fig. S22. High-resolution analysis of rank-dependent redistribution.**
Decile-level (10-bin) upward redistribution fractions plotted across the Empty percentile baseline for each chaperone condition. Protein-based chaperones exhibit a distinct amplification peak within the 60$^{th}$–80$^{th}$ percentile range, whereas Seq576 shows a gradual redistribution pattern without a sharp mid-high spike. A significant condition × quantile interaction was observed ($F$ = 2.45, $p$ = 0.0011). A planned contrast comparing the mean of GroEL, DnaK, and Spy with Seq576 across Q7–Q8 was significant ($t(9)$ = 5.91, Holm-adjusted $p$ = 0.00068). Values represent mean ± SD across replicates.

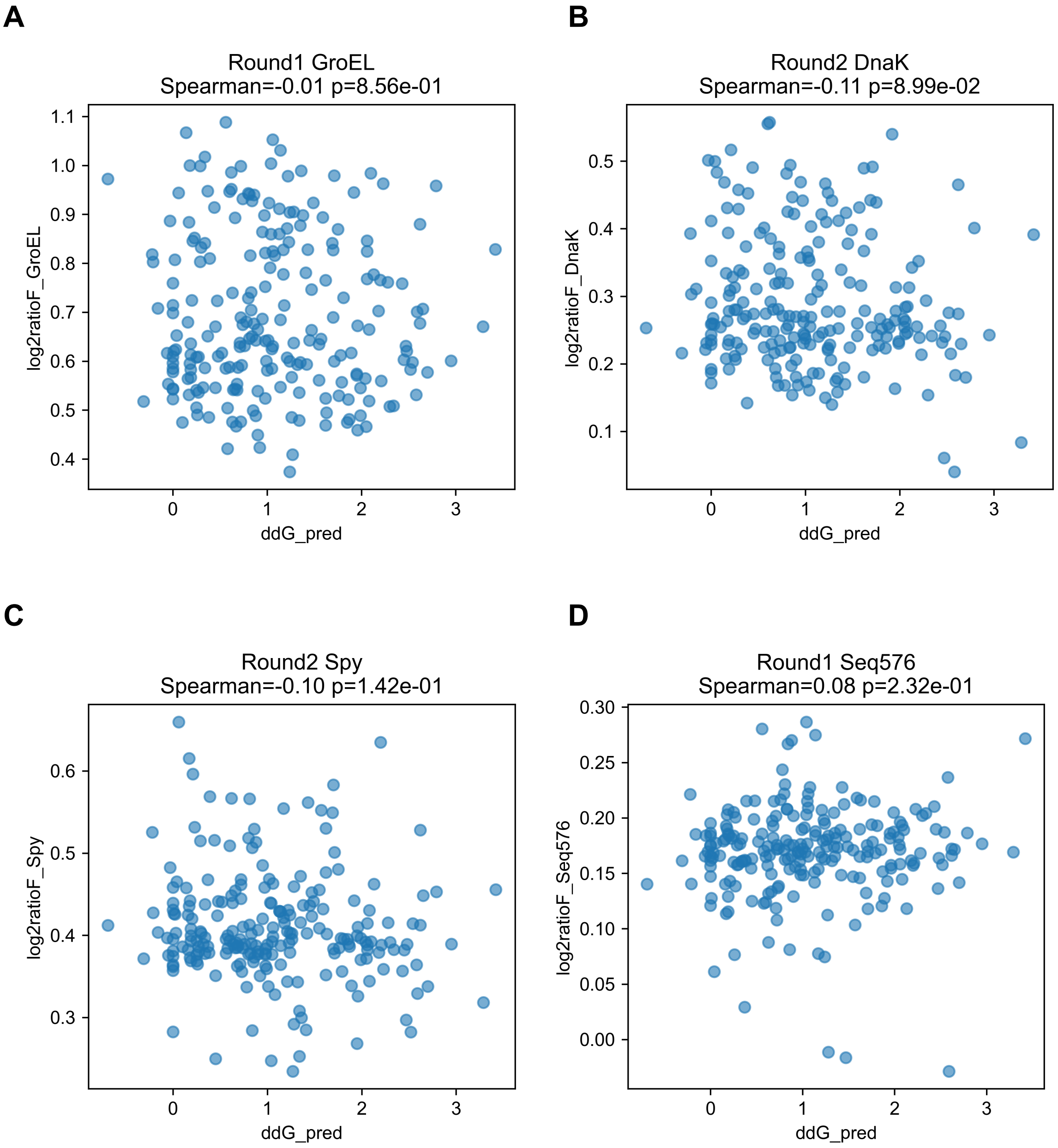


**Fig. S23. Predicted mutation stability does not correlate with chaperone-dependent folding effects.**

Predicted stability changes (ΔΔG_pred) for alanine substitutions were calculated using ThermoMPNN based on the reporter protein structure (PDB ID: 4KGF). Predicted ΔΔG values were compared with experimentally inferred chaperone-dependent folding effects ($log_2$ratioF) under GroEL (**A**), DnaK (**B**), Spy (**C**), and Seq576 (**D**) conditions. Each point represents a single alanine substitution. Spearman correlation coefficients indicate no significant relationship between predicted stability and chaperone-dependent folding effects.

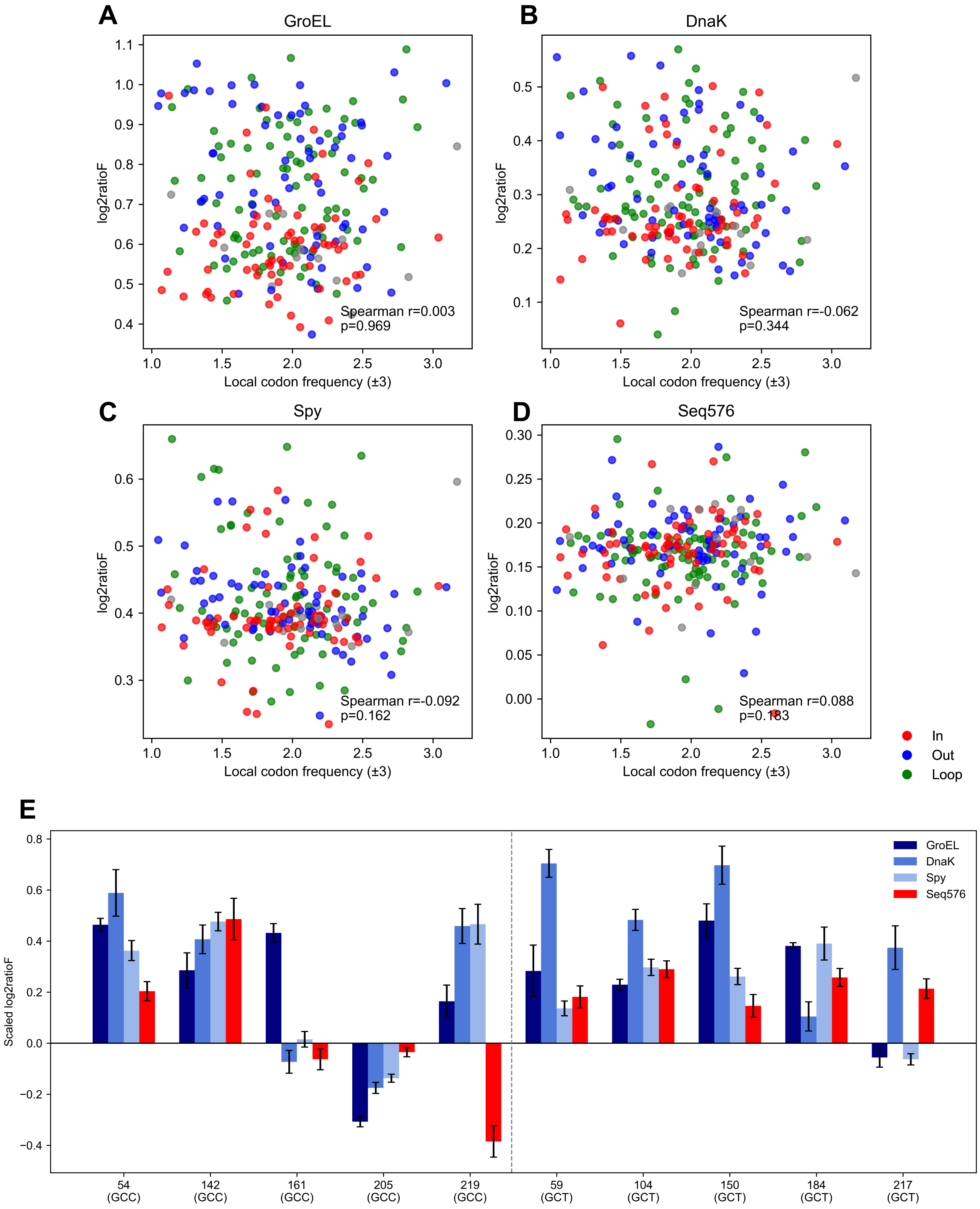


**Fig. S24. Codon usage does not explain chaperone-dependent folding rescue.**
(**A–D**) Scatter plots comparing local codon frequency (±3 residues) with residue-level $log_2$ratioF for GroEL, DnaK, Spy, and Seq576. Points are colored by structural exposure class (In, Out, Loop). Spearman correlation coefficients and p-values are indicated.
(**E**) Comparison of scaled $log_2$ratioF values for native alanine residues encoded by GCC or GCT across the four chaperone conditions. Bars show mean scaled $log_2$ratioF values, with error bars representing SEM.

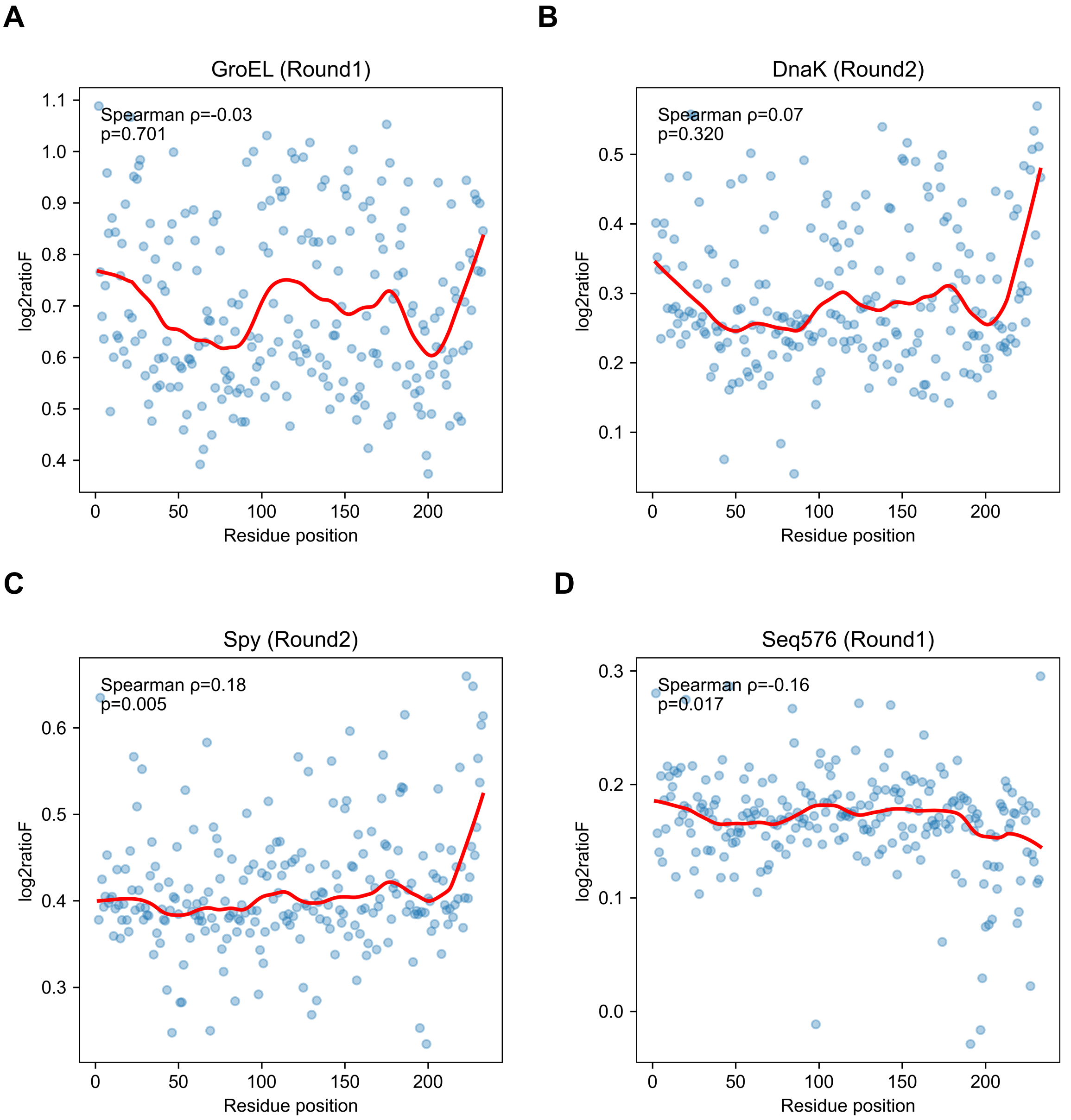


**Fig. S25. Positional trends of chaperone-dependent folding effects along the sequence.** Scatter plots show residue position versus $log_2$ratioF for each condition (GroEL (**A**), DnaK (**B**), Spy (**C**), and Seq576 (**D**)). Each point represents a residue from the alanine-scanning library. Red curves indicate LOWESS smoothing (fraction = 0.18) used to visualize large-scale positional trends. Spearman correlation coefficients and p-values are shown in each panel. Weak but significant positional correlations were observed for Seq576 and Spy, whereas GroEL and DnaK showed no significant positional relationship.

### SI Tables

#### Table S1. Bacterial strains and plasmids

| Strain Number | Strains | Plasmid1 | Marker | Plasmid2 | Marker | References |
|---|---|---|---|---|---|---|
| AS181 | MC4100(DE3) | pBAD/HisD-TagRFP675 | ApR | pBAD33mut-Empty | CmR | (*14*) |
| AS183 | MC4100(DE3) | pBAD/HisD-TagRFP675 | ApR | pBAD33-GroEL | CmR | (*14*) |
| AS197 | MC4100(DE3) | pBAD/HisD-TagRFP675 | ApR | pBAD33mut-Seq576 | CmR | (*14*) |
| AS491 | MC4100(DE3) | pBAD/HisD-TagRFP675 | ApR | pBAD33-DnaK | CmR | (*14*) |
| AS497 | MC4100(DE3) | pBAD/HisD-TagRFP675 | ApR | pBAD33-Spy | CmR | (*14*) |
| AS605 | MC4100(DE3) | pBAD/HisD-TagRFP675<br>Alanine mutation library<br>(hereafter referred to as the “Ala library”) | ApR | pBAD33mut-Empty | CmR | This study |
| AS611 | MC4100(DE3) | Ala library | ApR | pBAD33-GroEL | CmR | This study |
| AS608 | MC4100(DE3) | Ala library | ApR | pBAD33mut-Seq576 | CmR | This study |
| AS883 | MC4100(DE3) | Ala library | ApR | pBAD33-DnaK | CmR | This study |
| AS626 | MC4100(DE3) | Ala library | ApR | pBAD33-Spy | CmR | This study |
| AS632 | MC4100(DE3) | RD1-1 sorted Ala library (Low) | ApR | pBAD33mut-Empty | CmR | This study |
| AS633 | MC4100(DE3) | RD1-1 sorted Ala library (High) | ApR | pBAD33mut-Empty | CmR | This study |
| AS634 | MC4100(DE3) | RD1-1 sorted Ala library (Low) | ApR | pBAD33mut-Seq576 | CmR | This study |
| AS635 | MC4100(DE3) | RD1-1 sorted Ala library (High) | ApR | pBAD33mut-Seq576 | CmR | This study |
| AS636 | MC4100(DE3) | RD1-1 sorted Ala library (Low) | ApR | pBAD33-GroEL | CmR | This study |
| AS637 | MC4100(DE3) | RD1-1 sorted Ala library (High) | ApR | pBAD33-GroEL | CmR | This study |
| AS652 | MC4100(DE3) | RD1-1 re-transformed Ala library (Low) | ApR | pBAD33mut-Empty | CmR | This study |
| AS657 | MC4100(DE3) | RD1-1 re-transformed Ala library (High) | ApR | pBAD33mut-Empty | CmR | This study |
| AS662 | MC4100(DE3) | RD1-1 re-transformed Ala library (Low) | ApR | pBAD33mut-Seq576 | CmR | This study |
| AS666 | MC4100(DE3) | RD1-1 re-transformed Ala library (High) | ApR | pBAD33mut-Seq576 | CmR | This study |
| AS671 | MC4100(DE3) | RD1-1 re-transformed Ala library (Low) | ApR | pBAD33-GroEL | CmR | This study |
| AS676 | MC4100(DE3) | RD1-1 re-transformed Ala library (High) | ApR | pBAD33-GroEL | CmR | This study |
| AS680 | MC4100(DE3) | RD1-2 sorted Ala library (Low) | ApR | pBAD33mut-Empty | CmR | This study |
| AS682 | MC4100(DE3) | RD1-2 sorted Ala library (High) | ApR | pBAD33mut-Empty | CmR | This study |
| AS684 | MC4100(DE3) | RD1-2 sorted Ala library (Low) | ApR | pBAD33mut-Seq576 | CmR | This study |
| AS686 | MC4100(DE3) | RD1-2 sorted Ala library (High) | ApR | pBAD33mut-Seq576 | CmR | This study |
| AS688 | MC4100(DE3) | RD1-2 sorted Ala library (Low) | ApR | pBAD33-GroEL | CmR | This study |
| AS690 | MC4100(DE3) | RD1-2 sorted Ala library (High) | ApR | pBAD33-GroEL | CmR | This study |
| AS692 | MC4100(DE3) | RD1-2 re-transformed Ala library (Low) | ApR | pBAD33mut-Empty | CmR | This study |
| AS697 | MC4100(DE3) | RD1-2 re-transformed Ala library (High) | ApR | pBAD33mut-Empty | CmR | This study |
| AS701 | MC4100(DE3) | RD1-2 re-transformed Ala library (Low) | ApR | pBAD33mut-Seq576 | CmR | This study |
| AS705 | MC4100(DE3) | RD1-2 re-transformed Ala library (High) | ApR | pBAD33mut-Seq576 | CmR | This study |
| AS709 | MC4100(DE3) | RD1-2 re-transformed Ala library (Low) | ApR | pBAD33-GroEL | CmR | This study |
| AS713 | MC4100(DE3) | RD1-2 re-transformed Ala library (High) | ApR | pBAD33-GroEL | CmR | This study |
| AS717 | MC4100(DE3) | RD1-3 sorted Ala library (Low) | ApR | pBAD33mut-Empty | CmR | This study |
| AS720 | MC4100(DE3) | RD1-3 sorted Ala library (High) | ApR | pBAD33mut-Empty | CmR | This study |
| AS723 | MC4100(DE3) | RD1-3 sorted Ala library (Low) | ApR | pBAD33mut-Seq576 | CmR | This study |
| AS726 | MC4100(DE3) | RD1-3 sorted Ala library (High) | ApR | pBAD33mut-Seq576 | CmR | This study |

**(Table S1. Continued)**

| Strain Number | Strains | Plasmid1 | Marker | Plasmid2 | Marker | References |
|---|---|---|---|---|---|---|
| AS729 | MC4100(DE3) | RD1-3 sorted Ala library (Low) | ApR | pBAD33-GroEL | CmR | This study |
| AS732 | MC4100(DE3) | RD1-3 sorted Ala library (High) | ApR | pBAD33-GroEL | CmR | This study |
| AS741 | MC4100(DE3) | RD1-3 re-transformed Ala library (Low) | ApR | pBAD33mut-Empty | CmR | This study |
| AS745 | MC4100(DE3) | RD1-3 re-transformed Ala library (High) | ApR | pBAD33mut-Empty | CmR | This study |
| AS749 | MC4100(DE3) | RD1-3 re-transformed Ala library (Low) | ApR | pBAD33mut-Seq576 | CmR | This study |
| AS753 | MC4100(DE3) | RD1-3 re-transformed Ala library (High) | ApR | pBAD33mut-Seq576 | CmR | This study |
| AS757 | MC4100(DE3) | RD1-3 re-transformed Ala library (Low) | ApR | pBAD33-GroEL | CmR | This study |
| AS761 | MC4100(DE3) | RD1-3 re-transformed Ala library (High) | ApR | pBAD33-GroEL | CmR | This study |
| AS956 | MC4100(DE3) | RD2-1 sorted Ala library (Low) | ApR | pBAD33mut-Empty | CmR | This study |
| AS959 | MC4100(DE3) | RD2-1 sorted Ala library (High) | ApR | pBAD33mut-Empty | CmR | This study |
| AS962 | MC4100(DE3) | RD2-1 sorted Ala library (Low) | ApR | pBAD33-DnaK | CmR | This study |
| AS965 | MC4100(DE3) | RD2-1 sorted Ala library (High) | ApR | pBAD33-DnaK | CmR | This study |
| AS968 | MC4100(DE3) | RD2-1 sorted Ala library (Low) | ApR | pBAD33-Spy | CmR | This study |
| AS971 | MC4100(DE3) | RD2-1 sorted Ala library (High) | ApR | pBAD33-Spy | CmR | This study |
| AS980 | MC4100(DE3) | RD2-2 sorted Ala library (Low) | ApR | pBAD33mut-Empty | CmR | This study |
| AS983 | MC4100(DE3) | RD2-2 sorted Ala library (High) | ApR | pBAD33mut-Empty | CmR | This study |
| AS986 | MC4100(DE3) | RD2-2 sorted Ala library (Low) | ApR | pBAD33-DnaK | CmR | This study |
| AS989 | MC4100(DE3) | RD2-2 sorted Ala library (High) | ApR | pBAD33-DnaK | CmR | This study |
| AS992 | MC4100(DE3) | RD2-2 sorted Ala library (Low) | ApR | pBAD33-Spy | CmR | This study |
| AS995 | MC4100(DE3) | RD2-2 sorted Ala library (High) | ApR | pBAD33-Spy | CmR | This study |
| AS1010 | MC4100(DE3) | RD2-3 sorted Ala library (Low) | ApR | pBAD33mut-Empty | CmR | This study |
| AS1013 | MC4100(DE3) | RD2-3 sorted Ala library (High) | ApR | pBAD33mut-Empty | CmR | This study |
| AS1016 | MC4100(DE3) | RD2-3 sorted Ala library (Low) | ApR | pBAD33-DnaK | CmR | This study |
| AS1019 | MC4100(DE3) | RD2-3 sorted Ala library (High) | ApR | pBAD33-DnaK | CmR | This study |
| AS1022 | MC4100(DE3) | RD2-3 sorted Ala library (Low) | ApR | pBAD33-Spy | CmR | This study |
| AS1025 | MC4100(DE3) | RD2-3 sorted Ala library (High) | ApR | pBAD33-Spy | CmR | This study |
| AS1038 | MC4100(DE3) | RD2-4 sorted Ala library (Low) | ApR | pBAD33mut-Empty | CmR | This study |
| AS1041 | MC4100(DE3) | RD2-4 sorted Ala library (High) | ApR | pBAD33mut-Empty | CmR | This study |
| AS1044 | MC4100(DE3) | RD2-4 sorted Ala library (Low) | ApR | pBAD33-DnaK | CmR | This study |
| AS1047 | MC4100(DE3) | RD2-4 sorted Ala library (High) | ApR | pBAD33-DnaK | CmR | This study |
| AS1050 | MC4100(DE3) | RD2-4 sorted Ala library (Low) | ApR | pBAD33-Spy | CmR | This study |
| AS1053 | MC4100(DE3) | RD2-4 sorted Ala library (High) | ApR | pBAD33-Spy | CmR | This study |
| AS1066 | MC4100(DE3) | RD2-5 sorted Ala library (Low) | ApR | pBAD33mut-Empty | CmR | This study |
| AS1069 | MC4100(DE3) | RD2-5 sorted Ala library (High) | ApR | pBAD33mut-Empty | CmR | This study |
| AS1072 | MC4100(DE3) | RD2-5 sorted Ala library (Low) | ApR | pBAD33-DnaK | CmR | This study |
| AS1075 | MC4100(DE3) | RD2-5 sorted Ala library (High) | ApR | pBAD33-DnaK | CmR | This study |
| AS1078 | MC4100(DE3) | RD2-5 sorted Ala library (Low) | ApR | pBAD33-Spy | CmR | This study |
| AS1081 | MC4100(DE3) | RD2-5 sorted Ala library (High) | ApR | pBAD33-Spy | CmR | This study |
| AS1094 | MC4100(DE3) | RD2-2 re-transformed Ala library (Low) | ApR | pBAD33mut-Empty | CmR | This study |
| AS1097 | MC4100(DE3) | RD2-2 re-transformed Ala library (High) | ApR | pBAD33mut-Empty | CmR | This study |

**(Table S1. Continued)**

| Strain Number | Strains | Plasmid1 | Marker | Plasmid2 | Marker | References |
|---|---|---|---|---|---|---|
| AS1100 | MC4100(DE3) | RD2-3 re-transformed Ala library (Low) | ApR | pBAD33mut-Empty | CmR | This study |
| AS1103 | MC4100(DE3) | RD2-3 re-transformed Ala library (High) | ApR | pBAD33mut-Empty | CmR | This study |
| AS1106 | MC4100(DE3) | RD2-4 re-transformed Ala library (Low) | ApR | pBAD33mut-Empty | CmR | This study |
| AS1109 | MC4100(DE3) | RD2-4 re-transformed Ala library (High) | ApR | pBAD33mut-Empty | CmR | This study |
| AS1112 | MC4100(DE3) | RD2-1 re-transformed Ala library (Low) | ApR | pBAD33-DnaK | CmR | This study |
| AS1115 | MC4100(DE3) | RD2-1 re-transformed Ala library (High) | ApR | pBAD33-DnaK | CmR | This study |
| AS1118 | MC4100(DE3) | RD2-2 re-transformed Ala library (Low) | ApR | pBAD33-DnaK | CmR | This study |
| AS1121 | MC4100(DE3) | RD2-2 re-transformed Ala library (High) | ApR | pBAD33-DnaK | CmR | This study |
| AS1124 | MC4100(DE3) | RD2-4 re-transformed Ala library (Low) | ApR | pBAD33-DnaK | CmR | This study |
| AS1127 | MC4100(DE3) | RD2-4 re-transformed Ala library (High) | ApR | pBAD33-DnaK | CmR | This study |
| AS1136 | MC4100(DE3) | RD2-2 re-transformed Ala library (Low) | ApR | pBAD33-Spy | CmR | This study |
| AS1139 | MC4100(DE3) | RD2-2 re-transformed Ala library (High) | ApR | pBAD33-Spy | CmR | This study |
| AS1142 | MC4100(DE3) | RD2-3 re-transformed Ala library (Low) | ApR | pBAD33-Spy | CmR | This study |
| AS1145 | MC4100(DE3) | RD2-3 re-transformed Ala library (High) | ApR | pBAD33-Spy | CmR | This study |
| AS1148 | MC4100(DE3) | RD2-4 re-transformed Ala library (Low) | ApR | pBAD33-Spy | CmR | This study |
| AS1151 | MC4100(DE3) | RD2-4 re-transformed Ala library (High) | ApR | pBAD33-Spy | CmR | This study |
| AS1154 | MC4100(DE3) | RD2-5 re-transformed Ala library (Low) | ApR | pBAD33-Spy | CmR | This study |
| AS1157 | MC4100(DE3) | RD2-5 re-transformed Ala library (High) | ApR | pBAD33-Spy | CmR | This study |
| AS1284 | MC4100(DE3) | pBAD/HisD-TagRFP675(H10A) | ApR | pBAD33mut-Empty | CmR | This study |
| AS1286 | MC4100(DE3) | pBAD/HisD-TagRFP675(H10A) | ApR | pBAD33-GroEL | CmR | This study |
| AS1288 | MC4100(DE3) | pBAD/HisD-TagRFP675(H10A) | ApR | pBAD33-DnaK | CmR | This study |
| AS1290 | MC4100(DE3) | pBAD/HisD-TagRFP675(H10A) | ApR | pBAD33-Spy | CmR | This study |
| AS1292 | MC4100(DE3) | pBAD/HisD-TagRFP675(H10A) | ApR | pBAD33mut-Seq576 | CmR | This study |
| AS1296 | MC4100(DE3) | pBAD/HisD-TagRFP675(M11A) | ApR | pBAD33mut-Empty | CmR | This study |
| AS1298 | MC4100(DE3) | pBAD/HisD-TagRFP675(M11A) | ApR | pBAD33-GroEL | CmR | This study |
| AS1300 | MC4100(DE3) | pBAD/HisD-TagRFP675(M11A) | ApR | pBAD33-DnaK | CmR | This study |
| AS1302 | MC4100(DE3) | pBAD/HisD-TagRFP675(M11A) | ApR | pBAD33-Spy | CmR | This study |
| AS1304 | MC4100(DE3) | pBAD/HisD-TagRFP675(M11A) | ApR | pBAD33mut-Seq576 | CmR | This study |
| AS1308 | MC4100(DE3) | pBAD/HisD-TagRFP675(N20A) | ApR | pBAD33mut-Empty | CmR | This study |
| AS1310 | MC4100(DE3) | pBAD/HisD-TagRFP675(N20A) | ApR | pBAD33-GroEL | CmR | This study |
| AS1312 | MC4100(DE3) | pBAD/HisD-TagRFP675(N20A) | ApR | pBAD33-DnaK | CmR | This study |
| AS1314 | MC4100(DE3) | pBAD/HisD-TagRFP675(N20A) | ApR | pBAD33-Spy | CmR | This study |
| AS1316 | MC4100(DE3) | pBAD/HisD-TagRFP675(N20A) | ApR | pBAD33mut-Seq576 | CmR | This study |
| AS1320 | MC4100(DE3) | pBAD/HisD-TagRFP675(S28A) | ApR | pBAD33mut-Empty | CmR | This study |
| AS1322 | MC4100(DE3) | pBAD/HisD-TagRFP675(S28A) | ApR | pBAD33-GroEL | CmR | This study |
| AS1324 | MC4100(DE3) | pBAD/HisD-TagRFP675(S28A) | ApR | pBAD33-DnaK | CmR | This study |
| AS1326 | MC4100(DE3) | pBAD/HisD-TagRFP675(S28A) | ApR | pBAD33-Spy | CmR | This study |
| AS1328 | MC4100(DE3) | pBAD/HisD-TagRFP675(S28A) | ApR | pBAD33mut-Seq576 | CmR | This study |
| AS1332 | MC4100(DE3) | pBAD/HisD-TagRFP675(G37A) | ApR | pBAD33mut-Empty | CmR | This study |
| AS1334 | MC4100(DE3) | pBAD/HisD-TagRFP675(G37A) | ApR | pBAD33-GroEL | CmR | This study |

**(Table S1. Continued)**

| Strain Number | Strains | Plasmid1 | Marker | Plasmid2 | Marker | References |
|---|---|---|---|---|---|---|
| AS1336 | MC4100(DE3) | pBAD/HisD-TagRFP675(G37A) | ApR | pBAD33-DnaK | CmR | This study |
| AS1338 | MC4100(DE3) | pBAD/HisD-TagRFP675(G37A) | ApR | pBAD33-Spy | CmR | This study |
| AS1340 | MC4100(DE3) | pBAD/HisD-TagRFP675(G37A) | ApR | pBAD33mut-Seq576 | CmR | This study |
| AS1344 | MC4100(DE3) | pBAD/HisD-TagRFP675(S66A) | ApR | pBAD33mut-Empty | CmR | This study |
| AS1346 | MC4100(DE3) | pBAD/HisD-TagRFP675(S66A) | ApR | pBAD33-GroEL | CmR | This study |
| AS1348 | MC4100(DE3) | pBAD/HisD-TagRFP675(S66A) | ApR | pBAD33-DnaK | CmR | This study |
| AS1350 | MC4100(DE3) | pBAD/HisD-TagRFP675(S66A) | ApR | pBAD33-Spy | CmR | This study |
| AS1352 | MC4100(DE3) | pBAD/HisD-TagRFP675(S66A) | ApR | pBAD33mut-Seq576 | CmR | This study |
| AS1356 | MC4100(DE3) | pBAD/HisD-TagRFP675(Y96A) | ApR | pBAD33mut-Empty | CmR | This study |
| AS1358 | MC4100(DE3) | pBAD/HisD-TagRFP675(Y96A) | ApR | pBAD33-GroEL | CmR | This study |
| AS1360 | MC4100(DE3) | pBAD/HisD-TagRFP675(Y96A) | ApR | pBAD33-DnaK | CmR | This study |
| AS1362 | MC4100(DE3) | pBAD/HisD-TagRFP675(Y96A) | ApR | pBAD33-Spy | CmR | This study |
| AS1364 | MC4100(DE3) | pBAD/HisD-TagRFP675(Y96A) | ApR | pBAD33mut-Seq576 | CmR | This study |
| AS1368 | MC4100(DE3) | pBAD/HisD-TagRFP675(D98A) | ApR | pBAD33mut-Empty | CmR | This study |
| AS1370 | MC4100(DE3) | pBAD/HisD-TagRFP675(D98A) | ApR | pBAD33-GroEL | CmR | This study |
| AS1372 | MC4100(DE3) | pBAD/HisD-TagRFP675(D98A) | ApR | pBAD33-DnaK | CmR | This study |
| AS1374 | MC4100(DE3) | pBAD/HisD-TagRFP675(D98A) | ApR | pBAD33-Spy | CmR | This study |
| AS1376 | MC4100(DE3) | pBAD/HisD-TagRFP675(D98A) | ApR | pBAD33mut-Seq576 | CmR | This study |
| AS1380 | MC4100(DE3) | pBAD/HisD-TagRFP675(A104A) | ApR | pBAD33mut-Empty | CmR | This study |
| AS1382 | MC4100(DE3) | pBAD/HisD-TagRFP675(A104A) | ApR | pBAD33-GroEL | CmR | This study |
| AS1384 | MC4100(DE3) | pBAD/HisD-TagRFP675(A104A) | ApR | pBAD33-DnaK | CmR | This study |
| AS1386 | MC4100(DE3) | pBAD/HisD-TagRFP675(A104A) | ApR | pBAD33-Spy | CmR | This study |
| AS1388 | MC4100(DE3) | pBAD/HisD-TagRFP675(A104A) | ApR | pBAD33mut-Seq576 | CmR | This study |
| AS1392 | MC4100(DE3) | pBAD/HisD-TagRFP675(T108A) | ApR | pBAD33mut-Empty | CmR | This study |
| AS1394 | MC4100(DE3) | pBAD/HisD-TagRFP675(T108A) | ApR | pBAD33-GroEL | CmR | This study |
| AS1396 | MC4100(DE3) | pBAD/HisD-TagRFP675(T108A) | ApR | pBAD33-DnaK | CmR | This study |
| AS1398 | MC4100(DE3) | pBAD/HisD-TagRFP675(T108A) | ApR | pBAD33-Spy | CmR | This study |
| AS1400 | MC4100(DE3) | pBAD/HisD-TagRFP675(T108A) | ApR | pBAD33mut-Seq576 | CmR | This study |
| AS1404 | MC4100(DE3) | pBAD/HisD-TagRFP675(L110A) | ApR | pBAD33mut-Empty | CmR | This study |
| AS1406 | MC4100(DE3) | pBAD/HisD-TagRFP675(L110A) | ApR | pBAD33-GroEL | CmR | This study |
| AS1408 | MC4100(DE3) | pBAD/HisD-TagRFP675(L110A) | ApR | pBAD33-DnaK | CmR | This study |
| AS1410 | MC4100(DE3) | pBAD/HisD-TagRFP675(L110A) | ApR | pBAD33-Spy | CmR | This study |
| AS1412 | MC4100(DE3) | pBAD/HisD-TagRFP675(L110A) | ApR | pBAD33mut-Seq576 | CmR | This study |
| AS1416 | MC4100(DE3) | pBAD/HisD-TagRFP675(Q111A) | ApR | pBAD33mut-Empty | CmR | This study |
| AS1418 | MC4100(DE3) | pBAD/HisD-TagRFP675(Q111A) | ApR | pBAD33-GroEL | CmR | This study |
| AS1420 | MC4100(DE3) | pBAD/HisD-TagRFP675(Q111A) | ApR | pBAD33-DnaK | CmR | This study |
| AS1422 | MC4100(DE3) | pBAD/HisD-TagRFP675(Q111A) | ApR | pBAD33-Spy | CmR | This study |
| AS1424 | MC4100(DE3) | pBAD/HisD-TagRFP675(Q111A) | ApR | pBAD33mut-Seq576 | CmR | This study |
| AS1428 | MC4100(DE3) | pBAD/HisD-TagRFP675(C114A) | ApR | pBAD33mut-Empty | CmR | This study |
| AS1430 | MC4100(DE3) | pBAD/HisD-TagRFP675(C114A) | ApR | pBAD33-GroEL | CmR | This study |

**(Table S1. Continued)**

| Strain Number | Strains | Plasmid1 | Marker | Plasmid2 | Marker | References |
|---|---|---|---|---|---|---|
| AS1432 | MC4100(DE3) | pBAD/HisD-TagRFP675(C114A) | ApR | pBAD33-DnaK | CmR | This study |
| AS1434 | MC4100(DE3) | pBAD/HisD-TagRFP675(C114A) | ApR | pBAD33-Spy | CmR | This study |
| AS1436 | MC4100(DE3) | pBAD/HisD-TagRFP675(C114A) | ApR | pBAD33mut-Seq576 | CmR | This study |
| AS1440 | MC4100(DE3) | pBAD/HisD-TagRFP675(K120A) | ApR | pBAD33mut-Empty | CmR | This study |
| AS1442 | MC4100(DE3) | pBAD/HisD-TagRFP675(K120A) | ApR | pBAD33-GroEL | CmR | This study |
| AS1444 | MC4100(DE3) | pBAD/HisD-TagRFP675(K120A) | ApR | pBAD33-DnaK | CmR | This study |
| AS1446 | MC4100(DE3) | pBAD/HisD-TagRFP675(K120A) | ApR | pBAD33-Spy | CmR | This study |
| AS1448 | MC4100(DE3) | pBAD/HisD-TagRFP675(K120A) | ApR | pBAD33mut-Seq576 | CmR | This study |
| AS1452 | MC4100(DE3) | pBAD/HisD-TagRFP675(S128A) | ApR | pBAD33mut-Empty | CmR | This study |
| AS1454 | MC4100(DE3) | pBAD/HisD-TagRFP675(S128A) | ApR | pBAD33-GroEL | CmR | This study |
| AS1456 | MC4100(DE3) | pBAD/HisD-TagRFP675(S128A) | ApR | pBAD33-DnaK | CmR | This study |
| AS1458 | MC4100(DE3) | pBAD/HisD-TagRFP675(S128A) | ApR | pBAD33-Spy | CmR | This study |
| AS1460 | MC4100(DE3) | pBAD/HisD-TagRFP675(S128A) | ApR | pBAD33mut-Seq576 | CmR | This study |
| AS1464 | MC4100(DE3) | pBAD/HisD-TagRFP675(N158A) | ApR | pBAD33mut-Empty | CmR | This study |
| AS1466 | MC4100(DE3) | pBAD/HisD-TagRFP675(N158A) | ApR | pBAD33-GroEL | CmR | This study |
| AS1468 | MC4100(DE3) | pBAD/HisD-TagRFP675(N158A) | ApR | pBAD33-DnaK | CmR | This study |
| AS1470 | MC4100(DE3) | pBAD/HisD-TagRFP675(N158A) | ApR | pBAD33-Spy | CmR | This study |
| AS1472 | MC4100(DE3) | pBAD/HisD-TagRFP675(N158A) | ApR | pBAD33mut-Seq576 | CmR | This study |
| AS1476 | MC4100(DE3) | pBAD/HisD-TagRFP675(R197A) | ApR | pBAD33mut-Empty | CmR | This study |
| AS1478 | MC4100(DE3) | pBAD/HisD-TagRFP675(R197A) | ApR | pBAD33-GroEL | CmR | This study |
| AS1480 | MC4100(DE3) | pBAD/HisD-TagRFP675(R197A) | ApR | pBAD33-DnaK | CmR | This study |
| AS1482 | MC4100(DE3) | pBAD/HisD-TagRFP675(R197A) | ApR | pBAD33-Spy | CmR | This study |
| AS1484 | MC4100(DE3) | pBAD/HisD-TagRFP675(R197A) | ApR | pBAD33mut-Seq576 | CmR | This study |
| AS1488 | MC4100(DE3) | pBAD/HisD-TagRFP675(L232A) | ApR | pBAD33mut-Empty | CmR | This study |
| AS1490 | MC4100(DE3) | pBAD/HisD-TagRFP675(L232A) | ApR | pBAD33-GroEL | CmR | This study |
| AS1492 | MC4100(DE3) | pBAD/HisD-TagRFP675(L232A) | ApR | pBAD33-DnaK | CmR | This study |
| AS1494 | MC4100(DE3) | pBAD/HisD-TagRFP675(L232A) | ApR | pBAD33-Spy | CmR | This study |
| AS1496 | MC4100(DE3) | pBAD/HisD-TagRFP675(L232A) | ApR | pBAD33mut-Seq576 | CmR | This study |
| AS1502 | BL21(DE3) | pBAD/HisD-TagRFP675(Q111A) | ApR | | | This study |
| AS1506 | BL21(DE3) | pBAD/HisD-TagRFP675(L232A) | ApR | | | This study |

**Table S2. Summary of validation outcomes and replicate selection in Round 2 for downstream NGS analysis.**
Summary of validation outcomes for five independent sorting iterations in Round 2 across different chaperone conditions (Empty, DnaK, and Spy). Yellow boxes indicate replicates that passed all fluorescence-based validation criteria, including consistent separation between low- and high-folding populations in bulk fluorescence measurements and FACS profiles, and were selected for downstream next-generation sequencing (NGS) analysis. White boxes indicate replicates that failed validation or were not included in downstream analysis. Final replicate selection was based solely on quality control outcomes and not on iteration order.

| Sample / Cell sorting | Empty | DnaK | Spy |
|---|---|---|---|
| 2-1st | | DnaK 1 | |
| 2-2nd | Empty 2 | DnaK 2 | Spy 2 |
| 2-3rd | Empty 3 | | Spy 3 |
| 2-4th | Empty 4 | DnaK 4 | Spy 4 |
| 2-5th | | | Spy 5 |

**Table S3. Sequencing yield and quality metrics for CHAP-SEQ libraries included in downstream analysis.** (A) Round 1: Empty, GroEL, and Seq576 conditions (three biological replicates per condition). (B) Round 2: Empty and DnaK conditions, with three biological replicates per condition, and Spy, with four biological replicates. High- and low-fluorescence populations from each biological replicate were sequenced separately. Overall values represent yield-weighted summaries across the samples listed within each round.

**A. Round 1**

| Sorting batch | Condition | Sorted bin | Read pairs | Yield (Mbases) | Mean quality score | Bases ≥ Q30 (%) |
|---|---|---|---|---|---|---|
| 1-1st | Empty | Low | 4425877 | 1328 | 34.96 | 88.34 |
| | Empty | High | 4787567 | 1436 | 34.84 | 87.74 |
| | GroEL | Low | 3883625 | 1165 | 35.07 | 88.93 |
| | GroEL | High | 4305632 | 1292 | 35.02 | 88.64 |
| | Seq576 | Low | 3767810 | 1130 | 34.93 | 88.23 |
| | Seq576 | High | 4962643 | 1489 | 34.66 | 86.89 |
| 1-2nd | Empty | Low | 4560530 | 1368 | 35.05 | 88.82 |
| | Empty | High | 5036532 | 1511 | 34.95 | 88.3 |
| | GroEL | Low | 5538727 | 1662 | 34.81 | 87.63 |
| | GroEL | High | 5665523 | 1700 | 34.53 | 86.24 |
| | Seq576 | Low | 4615202 | 1385 | 34.88 | 87.98 |
| | Seq576 | High | 5328547 | 1599 | 34.9 | 88.09 |
| 1-3rd | Empty | Low | 4911297 | 1473 | 34.94 | 88.26 |
| | Empty | High | 5326660 | 1598 | 34.91 | 88.11 |
| | GroEL | Low | 5449088 | 1635 | 35.13 | 89.24 |
| | GroEL | High | 3979394 | 1194 | 35.03 | 88.74 |
| | Seq576 | Low | 4657039 | 1397 | 34.85 | 87.82 |
| | Seq576 | High | 4763097 | 1429 | 34.85 | 87.86 |
| **Overall** | | | **85,964,790** | **25,791** | **34.90** | **88.07** |

**B. Round 2**

| Sorting batch | Condition | Sorted bin | Read pairs | Yield (Mbases) | Mean quality score | Bases ≥ Q30 (%) |
|---|---|---|---|---|---|---|
| 2-1st | DnaK | Low | 2821915 | 847 | 37.81 | 89.26 |
| | DnaK | High | 2372299 | 712 | 37.68 | 88.6 |
| 2-2nd | Empty | Low | 2650491 | 795 | 37.73 | 88.87 |
| | Empty | High | 3018063 | 905 | 37.73 | 88.89 |
| | DnaK | Low | 2507260 | 752 | 37.77 | 89.09 |
| | DnaK | High | 2800567 | 840 | 37.75 | 89.01 |
| | Spy | Low | 2566466 | 770 | 37.94 | 89.8 |
| | Spy | High | 2659312 | 798 | 37.92 | 89.67 |
| 2-3rd | Empty | Low | 2723767 | 817 | 38.01 | 90.11 |
| | Empty | High | 2872706 | 862 | 37.83 | 89.26 |
| | Spy | Low | 2687775 | 806 | 37.8 | 89.21 |
| | Spy | High | 2496878 | 749 | 37.73 | 88.85 |
| 2-4th | Empty | Low | 2696903 | 809 | 37.88 | 89.47 |
| | Empty | High | 2298534 | 690 | 37.76 | 88.94 |
| | DnaK | Low | 2616672 | 785 | 38.15 | 90.79 |
| | DnaK | High | 2636813 | 791 | 37.84 | 89.28 |
| | Spy | Low | 2560838 | 768 | 37.59 | 88.24 |
| | Spy | High | 2723355 | 817 | 37.89 | 89.69 |
| 2-5th | Spy | Low | 2697562 | 809 | 37.99 | 90 |
| | Spy | High | 2643585 | 793 | 38.04 | 90.25 |
| **Overall** | | | **53,051,761** | **15,915** | **37.84** | **89.37** |

**Table S4. Sequencing depth stability across rounds and conditions.**
Depth ratios ($DP_H/DP_L$) were calculated for each variant within each replicate. Summary statistics are shown by round. The fraction of variants with zero effective depth (DP = 0) in either fluorescence bin is reported for each round-condition combination. No additional exclusion criteria were applied.

| **Depth ratio summary (round-level)** | | | | | |
|---|---|---|---|---|---|
| **Round** | **N variants** | **Mean ($DP_H/DP_L$)** | **SD** | **Median** | **5–95% range** |
| R1 | 1679 | 1.0069 | 0.0198 | 1.0018 | 0.993–1.046 |
| R2 | 1950 | 1.0075 | 0.0696 | 0.9993 | 0.911–1.189 |

| **Zero-depth fraction by condition** | | | | |
|---|---|---|---|---|
| **Round** | **Condition** | **frac_DP0** | **n_DP0** | **Total** |
| R1 | Empty | 0.125 | 87 | 696 |
| R1 | GroEL | 0.336 | 234 | 696 |
| R1 | Seq576 | 0.126 | 88 | 696 |
| R2 | Empty | 0.079 | 55 | 696 |
| R2 | DnaK | 0.228 | 159 | 696 |
| R2 | Spy | 0.168 | 156 | 928 |

**Table S5. Effect of quantile-based ε stabilization on pHigh estimates.**
Stabilized probabilities ($pHigh^{stabilized}$) were compared to raw probabilities ($pHigh^{raw}$) within each round. Spearman and Pearson correlation coefficients, as well as mean and maximum absolute differences, are reported. Stabilization was applied uniformly across variants prior to inversion of the log-normal gating model. Rows highlighted in yellow indicate quantile ranges where stabilized and raw pHigh estimates show highly consistent agreement (high correlations and minimal absolute differences), representing stable regimes of the ε stabilization procedure.

| Quantile | Condition | Spearman | Pearson | mean_abs_diff | max_abs_diff |
|---|---|---|---|---|---|
| Q1.25 | ALL | 0.998 | 0.986 | 0.047 | 0.272 |
| Q1.25 | Empty | 0.998 | 0.983 | 0.035 | 0.271 |
| Q1.25 | GroEL | 0.989 | 0.990 | 0.071 | 0.266 |
| Q1.25 | Seq576 | 0.998 | 0.982 | 0.036 | 0.272 |
| Q2.5 | ALL | 0.997 | 0.984 | 0.051 | 0.285 |
| Q2.5 | Empty | 0.997 | 0.980 | 0.037 | 0.284 |
| Q2.5 | GroEL | 0.988 | 0.989 | 0.077 | 0.279 |
| Q2.5 | Seq576 | 0.998 | 0.980 | 0.039 | 0.285 |
| Q5.0 | ALL | 0.997 | 0.979 | 0.059 | 0.314 |
| Q5.0 | Empty | 0.997 | 0.974 | 0.044 | 0.313 |
| Q5.0 | GroEL | 0.985 | 0.985 | 0.090 | 0.308 |
| Q5.0 | Seq576 | 0.997 | 0.973 | 0.045 | 0.314 |
| Q7.5 | ALL | 0.996 | 0.973 | 0.067 | 0.337 |
| Q7.5 | Empty | 0.995 | 0.968 | 0.049 | 0.336 |
| Q7.5 | GroEL | 0.981 | 0.981 | 0.102 | 0.331 |
| Q7.5 | Seq576 | 0.997 | 0.966 | 0.050 | 0.337 |
| Q10 | ALL | 0.995 | 0.970 | 0.072 | 0.350 |
| Q10 | Empty | 0.995 | 0.964 | 0.052 | 0.349 |
| Q10 | GroEL | 0.978 | 0.978 | 0.110 | 0.344 |
| Q10 | Seq576 | 0.996 | 0.961 | 0.053 | 0.350 |

**Table S6. Pairwise replicate correlation statistics for inferred Fmean values.**
Pairwise comparisons of inferred Fmean values between biological replicates for each round and condition. For each replicate pair (repA vs repB), Pearson correlation coefficient (*r*) and Spearman rank correlation coefficient (ρ) are reported. All correlations were statistically significant ($p < 10^{-25}$), indicating strong reproducibility across replicates. Rows highlighted in yellow indicate representative replicate comparisons demonstrating the consistently high reproducibility observed across rounds and conditions.

| Condition | Fmean metrics | repA | repB | Pearson *r* | Pearson *p* | Spearman ρ | Spearman *p* |
|---|---|---|---|---|---|---|---|
| Empty | pHigh_Q1.25 | 1 | 2 | 0.744 | 3.76E-42 | 0.666 | 3.65E-31 |
| Empty | pHigh_Q1.25 | 1 | 3 | 0.726 | 3.23E-39 | 0.652 | 1.63E-29 |
| Empty | pHigh_Q1.25 | 2 | 3 | 0.793 | 2.23E-51 | 0.694 | 1.27E-34 |
| GroEL | pHigh_Q1.25 | 1 | 2 | 0.924 | 7.03E-98 | 0.883 | 1.81E-77 |
| GroEL | pHigh_Q1.25 | 1 | 3 | 0.904 | 6.59E-87 | 0.876 | 8.91E-75 |
| GroEL | pHigh_Q1.25 | 2 | 3 | 0.934 | 4.11E-105 | 0.911 | 2.01E-90 |
| Seq576 | pHigh_Q1.25 | 1 | 2 | 0.762 | 3.01E-45 | 0.658 | 3.70E-30 |
| Seq576 | pHigh_Q1.25 | 1 | 3 | 0.616 | 1.16E-25 | 0.616 | 1.34E-25 |
| Seq576 | pHigh_Q1.25 | 2 | 3 | 0.681 | 5.50E-33 | 0.615 | 1.57E-25 |
| Empty | pHigh_Q2.5 | 1 | 2 | 0.747 | 1.07E-42 | 0.667 | 3.29E-31 |
| Empty | pHigh_Q2.5 | 1 | 3 | 0.730 | 6.88E-40 | 0.653 | 1.27E-29 |
| Empty | pHigh_Q2.5 | 2 | 3 | 0.797 | 2.95E-52 | 0.694 | 1.14E-34 |
| GroEL | pHigh_Q2.5 | 1 | 2 | 0.926 | 3.54E-99 | 0.884 | 7.60E-78 |
| GroEL | pHigh_Q2.5 | 1 | 3 | 0.907 | 3.63E-88 | 0.877 | 3.86E-75 |
| GroEL | pHigh_Q2.5 | 2 | 3 | 0.936 | 1.70E-106 | 0.912 | 9.73E-91 |
| Seq576 | pHigh_Q2.5 | 1 | 2 | 0.765 | 7.83E-46 | 0.659 | 2.95E-30 |
| Seq576 | pHigh_Q2.5 | 1 | 3 | 0.625 | 1.71E-26 | 0.617 | 1.02E-25 |
| Seq576 | pHigh_Q2.5 | 2 | 3 | 0.687 | 1.10E-33 | 0.614 | 1.90E-25 |
| Empty | pHigh_Q5.0 | 1 | 2 | 0.755 | 5.27E-44 | 0.668 | 2.39E-31 |
| Empty | pHigh_Q5.0 | 1 | 3 | 0.740 | 1.70E-41 | 0.656 | 6.22E-30 |
| Empty | pHigh_Q5.0 | 2 | 3 | 0.806 | 2.27E-54 | 0.696 | 5.99E-35 |
| GroEL | pHigh_Q5.0 | 1 | 2 | 0.930 | 3.53E-102 | 0.885 | 2.16E-78 |
| GroEL | pHigh_Q5.0 | 1 | 3 | 0.912 | 3.94E-91 | 0.879 | 8.62E-76 |
| GroEL | pHigh_Q5.0 | 2 | 3 | 0.940 | 8.80E-110 | 0.913 | 9.73E-92 |
| Seq576 | pHigh_Q5.0 | 1 | 2 | 0.772 | 3.08E-47 | 0.662 | 1.25E-30 |
| Seq576 | pHigh_Q5.0 | 1 | 3 | 0.644 | 1.46E-28 | 0.619 | 5.88E-26 |
| Seq576 | pHigh_Q5.0 | 2 | 3 | 0.699 | 2.19E-35 | 0.614 | 1.96E-25 |
| Empty | pHigh_Q7.5 | 1 | 2 | 0.761 | 4.22E-45 | 0.669 | 1.70E-31 |
| Empty | pHigh_Q7.5 | 1 | 3 | 0.748 | 7.45E-43 | 0.659 | 2.82E-30 |
| Empty | pHigh_Q7.5 | 2 | 3 | 0.814 | 3.62E-56 | 0.699 | 2.62E-35 |
| GroEL | pHigh_Q7.5 | 1 | 2 | 0.934 | 1.28E-104 | 0.887 | 5.44E-79 |
| GroEL | pHigh_Q7.5 | 1 | 3 | 0.917 | 1.36E-93 | 0.880 | 1.99E-76 |
| GroEL | pHigh_Q7.5 | 2 | 3 | 0.944 | 1.50E-112 | 0.915 | 1.53E-92 |
| Seq576 | pHigh_Q7.5 | 1 | 2 | 0.779 | 1.98E-48 | 0.665 | 5.87E-31 |
| Seq576 | pHigh_Q7.5 | 1 | 3 | 0.660 | 2.13E-30 | 0.622 | 3.46E-26 |
| Seq576 | pHigh_Q7.5 | 2 | 3 | 0.710 | 7.36E-37 | 0.613 | 2.22E-25 |
| Empty | pHigh_Q10 | 1 | 2 | 0.765 | 9.21E-46 | 0.671 | 1.13E-31 |
| Empty | pHigh_Q10 | 1 | 3 | 0.753 | 1.12E-43 | 0.661 | 1.53E-30 |
| Empty | pHigh_Q10 | 2 | 3 | 0.818 | 2.91E-57 | 0.700 | 1.52E-35 |
| GroEL | pHigh_Q10 | 1 | 2 | 0.936 | 4.65E-106 | 0.888 | 1.33E-79 |
| GroEL | pHigh_Q10 | 1 | 3 | 0.919 | 4.54E-95 | 0.882 | 3.97E-77 |
| GroEL | pHigh_Q10 | 2 | 3 | 0.946 | 3.09E-114 | 0.916 | 3.64E-93 |
| Seq576 | pHigh_Q10 | 1 | 2 | 0.782 | 3.73E-49 | 0.667 | 3.60E-31 |
| Seq576 | pHigh_Q10 | 1 | 3 | 0.670 | 1.49E-31 | 0.624 | 2.07E-26 |
| Seq576 | pHigh_Q10 | 2 | 3 | 0.716 | 9.09E-38 | 0.614 | 1.81E-25 |

**Table S7. Seq576 hotspot clustering and permutation analyses.** Sliding window and permutation-based hotspot statistics. Residues within the lower 5–25% of $log_2$ratioF values were analyzed across multiple percentile thresholds to evaluate sequence clustering robustness. Sliding-window enrichment (window size = 10 residues) was tested using a binomial survival test with Benjamini-Hochberg correction (FDR < 0.05), and longest-run significance was evaluated by permutation testing (10,000 permutations). Statistically significant fractions are highlighted in yellow. Across the tested percentile thresholds (5–25%), the 15% cutoff was selected as the representative threshold.

| Fraction | Condition | Threshold | Number of strong | Longest run length | Longest run regions | Permutation *p* value | Number of significant windows | Significant window regions |
|---|---|---|---|---|---|---|---|---|
| 0.05 | GroEL | 0.4755 | 12 | 2 | 199–200 | 0.4443 | 0 | |
| 0.05 | Seq576 | 0.1043 | 12 | 2 | 197–198; 219–220 | 0.4511 | 7 | 191–207 |
| 0.05 | DnaK | 0.1682 | 12 | 1 | 43, 46, 66, 77, 85, 98, 130, 157, 162, 169, 178, 204; | 1.0000 | 0 | |
| 0.05 | Spy | 0.3043 | 12 | 2 | 51–52 | 0.4549 | 0 | |
| 0.1 | GroEL | 0.5043 | 24 | 2 | 199–200 | 0.9306 | 0 | |
| 0.1 | Seq576 | 0.1212 | 24 | 4 | 197–200 | 0.0197 | 4 | 195–207 |
| 0.1 | DnaK | 0.1836 | 24 | 3 | 162–164 | 0.2022 | 0 | |
| 0.1 | Spy | 0.3448 | 24 | 3 | 51–53 | 0.1857 | 0 | |
| 0.15 | GroEL | 0.5312 | 35 | 3 | 63–65; 156–158 | 0.4993 | 0 | |
| 0.15 | Seq576 | 0.1353 | 35 | 6 | 197–202 | 0.0021 | 9 | 193–210 |
| 0.15 | DnaK | 0.2057 | 35 | 3 | 162–164 | 0.4926 | 0 | |
| 0.15 | Spy | 0.3627 | 35 | 3 | 51–53 | 0.4940 | 0 | |
| 0.2 | GroEL | 0.5459 | 47 | 4 | 156–159 | 0.2571 | 0 | |
| 0.2 | Seq576 | 0.1433 | 47 | 6 | 197–202 | 0.0085 | 6 | 195–209 |
| 0.2 | DnaK | 0.2206 | 47 | 4 | 199–202 | 0.2498 | 0 | |
| 0.2 | Spy | 0.3721 | 47 | 3 | 51–53 | 0.8119 | 0 | |
| 0.25 | GroEL | 0.5700 | 58 | 5 | 192–196 | 0.1416 | 2 | 192–202 |
| 0.25 | Seq576 | 0.1537 | 58 | 6 | 197–202 | 0.0347 | 4 | 196–208 |
| 0.25 | DnaK | 0.2287 | 58 | 4 | 199–202 | 0.4920 | 0 | |
| 0.25 | Spy | 0.3779 | 58 | 3 | 51–53 | 0.9600 | 0 | |

**Table S8. Relationships between structural dynamics (RMSD), folding metrics, and Seq576 hotspot enrichment.** (**A**) Residue-level correlations between RMSD and folding metrics. Spearman and Pearson correlations between residue RMSD values and experimentally inferred folding metrics (Fmean and $log_2$ratioF) were calculated across conditions. (**B**) Enrichment of structurally dynamic residues showing strong Seq576-dependent folding effects. Overlap analysis comparing residues within the top RMSD fraction and residues within the bottom $log_2$ratioF fraction across multiple percentile thresholds (5–25%). Statistical significance was evaluated using Fisher's exact test and permutation testing. Rows highlighted in yellow indicate statistically significant enrichment. Among the tested percentile thresholds, the 10% cutoff was selected as the representative threshold for subsequent analyses. (**C**) Sliding-window spatial association between RMSD and folding metrics. Sliding-window analysis was performed using a window size of 10 residues. For each window, the mean RMSD, mean $log_2$ratioF, fraction of residues within the top 10% of RMSD values, and fraction of residues within the bottom 10% of $log_2$ratioF values were calculated. Spearman correlations were calculated between mean RMSD and mean $log_2$ratioF and, separately, between the top-10% RMSD fraction and the bottom-10% $log_2$ratioF fraction.

**A. Residue-level correlation (RMSD vs folding)**

| Round | Type | Condition | Spearman *ρ* | Spearman *p* | Pearson *r* | Pearson *p* |
|---|---|---|---|---|---|---|
| Round1 | Fmean | Empty | 0.249 | 0.00017 | 0.256 | 0.000111 |
| Round1 | Fmean | GroEL | 0.243 | 0.00025 | 0.322 | 0.000001 |
| Round1 | Fmean | Seq576 | 0.182 | 0.00638 | 0.197 | 0.003201 |
| Round2 | Fmean | Empty | 0.216 | 0.00119 | 0.241 | 0.000289 |
| Round2 | Fmean | DnaK | 0.273 | 0.00004 | 0.255 | 0.000116 |
| Round2 | Fmean | Spy | 0.279 | 0.00002 | 0.299 | 0.000006 |

| Round | Type | Condition | Spearman *ρ* | Spearman *p* | Pearson *r* | Pearson *p* |
|---|---|---|---|---|---|---|
| Round1 | $log_2$ratioF | GroEL | 0.236 | 0.00039 | 0.273 | 0.000036 |
| Round1 | $log_2$ratioF | Seq576 | -0.096 | 0.15488 | -0.174 | 0.009110 |
| Round2 | $log_2$ratioF | DnaK | 0.244 | 0.00023 | 0.218 | 0.001060 |
| Round2 | $log_2$ratioF | Spy | 0.251 | 0.00015 | 0.293 | 0.000008 |

**B. Enrichment analysis**

| Q | Condition | High-RMSD residues, *n* | Low-$log_2$ratioF residues, *n* | Overlap, *n* | Fisher's exact *p* | Permutation *p* |
|---|---|---|---|---|---|---|
| 0.05 | $log_2$ratioF GroEL | 12 | 12 | 0 | 1 | 1 |
| 0.05 | $log_2$ratioF Seq576 | 12 | 12 | 3 | 0.020 | 0.019 |
| 0.05 | $log_2$ratioF DnaK | 12 | 12 | 0 | 1 | 1 |
| 0.05 | $log_2$ratioF Spy | 12 | 12 | 0 | 1 | 1 |
| 0.1 | $log_2$ratioF GroEL | 23 | 23 | 1 | 0.929 | 0.926 |
| 0.1 | $log_2$ratioF Seq576 | 23 | 23 | 6 | 0.019 | 0.018 |
| 0.1 | $log_2$ratioF DnaK | 23 | 23 | 0 | 1 | 1 |
| 0.1 | $log_2$ratioF Spy | 23 | 23 | 0 | 1 | 1 |
| 0.15 | $log_2$ratioF GroEL | 34 | 34 | 2 | 0.982 | 0.980 |
| 0.15 | $log_2$ratioF Seq576 | 34 | 34 | 8 | 0.117 | 0.121 |
| 0.15 | $log_2$ratioF DnaK | 34 | 34 | 1 | 0.998 | 0.998 |
| 0.15 | $log_2$ratioF Spy | 34 | 34 | 2 | 0.982 | 0.979 |
| 0.2 | $log_2$ratioF GroEL | 45 | 45 | 2 | 1 | 1 |
| 0.2 | $log_2$ratioF Seq576 | 45 | 45 | 14 | 0.037 | 0.037 |
| 0.2 | $log_2$ratioF DnaK | 45 | 45 | 3 | 0.999 | 0.998 |
| 0.2 | $log_2$ratioF Spy | 45 | 45 | 2 | 1 | 1 |
| 0.25 | $log_2$ratioF GroEL | 56 | 56 | 6 | 0.999 | 1 |
| 0.25 | $log_2$ratioF Seq576 | 56 | 56 | 19 | 0.059 | 0.059 |
| 0.25 | $log_2$ratioF DnaK | 56 | 56 | 8 | 0.992 | 0.992 |
| 0.25 | $log_2$ratioF Spy | 56 | 56 | 6 | 0.999 | 1 |

**(Table S6. Continued)**

**C. Sliding window correlation**

| Condition | Window-level RMSD metric (X) | Window-level folding metric (Y) | Spearman ρ | Spearman p | Number of windows |
|---|---|---|---|---|---|
| $log_2$ratioF GroEL | mean RMSD | mean $log_2$ratioF | 0.179 | 0.0086 | 214 |
| $log_2$ratioF Seq576 | mean RMSD | mean $log_2$ratioF | -0.386 | 5.06E-09 | 214 |
| $log_2$ratioF DnaK | mean RMSD | mean $log_2$ratioF | 0.260 | 0.0001 | 214 |
| $log_2$ratioF Spy | mean RMSD | mean $log_2$ratioF | 0.348 | 1.80E-07 | 214 |
| $log_2$ratioF GroEL | Frac_RMSD_top10 | frac_log2_bottom10 | -0.048 | 0.4845 | 214 |
| $log_2$ratioF Seq576 | Frac_RMSD_top10 | frac_log2_bottom10 | 0.326 | 1.07E-06 | 214 |
| $log_2$ratioF DnaK | Frac_RMSD_top10 | frac_log2_bottom10 | -0.309 | 3.98E-06 | 214 |
| $log_2$ratioF Spy | Frac_RMSD_top10 | frac_log2_bottom10 | -0.216 | 0.0015 | 214 |

**Table S9. Quantitative metrics of Q3–Q4 amplification and rich-getting-richer (RGR) onset location.**

The table reports replicate-level mean upward redistribution in Q3 (40$^{th}$–60$^{th}$ percentile) and Q4 (60$^{th}$–80$^{th}$ percentile), Q4–Q3 jump magnitude (mean ± SD), and the percentile corresponding to the maximal local slope (RGR onset candidate).

| Condition | Q3 | Q4 | Jump Δ(Q4–Q3) | Jump sd | Max slope (%) | Max slope |
|---|---|---|---|---|---|---|
| GroEL | 0.340 | 0.464 | 0.123 | 0.044 | 0.70 | 2.249 |
| Seq576 | 0.348 | 0.399 | 0.051 | 0.018 | 0.62 | 1.786 |
| DnaK | 0.319 | 0.446 | 0.127 | 0.010 | 0.60 | 2.679 |
| Spy | 0.319 | 0.464 | 0.147 | 0.089 | 0.56 | 2.679 |

**Movie S1.** Rotating structural visualizations of $log_2$ratioF values under GroEL condition (corresponding to Fig. S12A).
**Movie S2.** Rotating structural visualizations of $log_2$ratioF values under DnaK condition (corresponding to Fig. S12B).
**Movie S3.** Rotating structural visualizations of $log_2$ratioF values under Spy condition (corresponding to Fig. S12C).
**Movie S4.** Rotating structural visualizations of $log_2$ratioF values under Seq576 condition (corresponding to Figs. 2F and S12D).

**Movie S5.** Rotating visualizations of the TagRFP675 structural ensemble used for RMSD analysis (corresponding to Fig. S14A).
**Movie S6.** Rotating structural visualizations of residue-level RMSD values mapped onto TagRFP675 (corresponding to Fig. S14B).

**Movie S7.** Rotating structural visualization of Seq576 (G4)-specific effectiveness outliers (corresponding to Fig. 4A). Sphere size reflects the effectiveness-score (ES) category (ES3–ES5).
**Movie S8.** Rotating structural visualization of protein-based-chaperone-specific and all-three-concordant effectiveness outliers (corresponding to Fig. 4B). Sphere size reflects the effectiveness-score (ES) category (ES3–ES5).

**Movie S9.** Rotating structural visualizations of residue-level sd_log2 distribution under GroEL condition (corresponding to Fig. S16A). Sphere size reflects propagated between-replicate variability (QD1–QD5 stratification), and ribbon coloring represents $log_2$ratioF.
**Movie S10.** Rotating structural visualizations of residue-level sd_log2 distribution under DnaK condition (corresponding to Fig. S16B). Sphere size reflects propagated between-replicate (QD1–QD5 stratification), and ribbon coloring represents $log_2$ratioF.
**Movie S11.** Rotating structural visualizations of residue-level sd_log2 distribution under Spy condition (corresponding to Fig. S16C). Sphere size reflects propagated between-replicate (QD1–QD5 stratification), and ribbon coloring represents $log_2$ratioF.
**Movie S12.** Rotating structural visualizations of residue-level sd_log2 distribution under Seq576 condition (corresponding to Fig. S16D). Sphere size reflects propagated between-replicate (QD1–QD5 stratification), and ribbon coloring represents $log_2$ratioF.

**Movie S13.** Rotating structural visualizations of statistically defined combined outliers under GroEL condition (corresponding to Fig. S17). Combined outliers satisfy both the LOESS residual (top 5%) and Mahalanobis distance threshold (97.5th percentile of the $\chi^2$ distribution, df = 2).
**Movie S14.** Rotating structural visualizations of statistically defined combined outliers under DnaK condition (corresponding to Fig. S17F). Combined outliers satisfy both the LOESS residual (top 5%) and Mahalanobis distance threshold (97.5th percentile of the $\chi^2$ distribution, df = 2).
**Movie S15.** Rotating structural visualizations of statistically defined combined outliers under Spy condition (corresponding to Fig. S17G). Combined outliers satisfy both the LOESS residual (top 5%) and Mahalanobis distance threshold (97.5th percentile of the $\chi^2$ distribution, df = 2).
**Movie S16.** Rotating structural visualizations of statistically defined combined outliers under Seq576 condition (corresponding to Fig. S17H). Combined outliers satisfy both the LOESS residual (top 5%) and Mahalanobis distance threshold (97.5th percentile of the $\chi^2$ distribution, df = 2).

**Dataset S1A–S1H. Tabulated sequencing and folding datasets.** These datasets include raw and processed sequencing counts, pHigh values, inferred mean fluorescence values (Fmean; internally designed Fhigh in the analysis file), and residue-level folding metrics (including Fmean, log2ratioF, and SD) for all variants across experimental conditions.
Dataset S1A (step0_Round1_processed_raw_data.xlsx). Raw sequencing counts for Round 1 samples prior to filtering.
Dataset S1B (step0_Round2_processed_raw_data.xlsx). Raw sequencing counts for Round 2 samples prior to filtering.
Dataset S1C (step1_StrictGCGfiltering_Round1.csv). Filtered counts for Round 1 samples after strict alanine (GCG) codon validation and coverage filtering.

Dataset S1D (step1_StrictGCGfiltering_Round2.csv). Filtered counts for Round 2 samples after strict alanine (GCG) codon validation and coverage filtering.
Dataset S1E (step2_Fhigh_from_pHigh_Round1.csv). Inferred fluorescence (Fhigh) values for Round 1 variants derived from pHigh values.
Dataset S1F (step2_Fhigh_from_pHigh_Round2.csv). Inferred fluorescence (Fhigh) values for Round 2 variants derived from pHigh values.
Dataset S1G (Step3_Residue_Folding_Metrics_Round1.csv). Residue-level folding metrics for Round 1, including Fmean, log2ratioF, SD, and derived analysis metrics.
Dataset S1H (Step3_Residue_Folding_Metrics_Round2.csv). Residue-level folding metrics for Round 2, including Fmean, log2ratioF, SD, and derived analysis metrics.

**Dataset S2. Structure files used for visualization and structural analysis.** This dataset includes PDB files with residue-level mapping of folding metrics (including log2ratioF and SD) for each chaperone condition, RMSD-derived structural datasets, and a CIF file containing AlphaFold3 structural models. Dataset S2 is publicly available through Zenodo (https://doi.org/10.5281/zenodo.20274818).
